# Growth-driven mechanical buckling as a mechanism of age-related tortuosity in human cerebral arteries

**DOI:** 10.64898/2026.11.04.764114

**Authors:** Tomoaki Akiyama, Koji Yoshimoto, Takashi Miura

## Abstract

Human cerebral arteries become increasingly, sometimes abnormally, winding with age, but the mechanical basis of this change remains incompletely understood. To examine how arterial bends form and what sets their spacing, we modeled a cerebral artery as a slender elastic vessel supported by surrounding tissue. The distance between its ends remains constant, and the vessel can bend only in a plane tangent to the brain surface. As its natural length increases, axial compression builds until the vessel buckles. Bending partly relieves the compression, and the resulting waves grow in amplitude as elongation continues. Arterial bending stiffness and tissue support determine the wavelength, which is much larger than and proportional to vessel diameter, with comparatively weak dependence on other parameters. Numerical simulations captured a pre-buckling phase, rapid amplification of waves, and slow post-buckling growth during continued elongation. We evaluated these predictions using three-dimensional centerlines of cerebral arteries reconstructed from magnetic resonance angiography in healthy adults. Older age was associated with greater arterial length and excess-length ratio. For arterial waves oriented mostly parallel to the brain surface, wavelength increased with vessel diameter. Both this relationship and the observed wavelength-to-diameter ratio were consistent with model predictions. However, wave amplitude was not significantly associated with age. Overall, the modeling and imaging results support growth-driven buckling and post-buckling deformation under spatial constraint as a plausible mechanical explanation for age-related cerebral arterial tortuosity. This framework links vascular aging to growth-driven instability and pattern formation, with potential applications to diseased arteries beyond the brain.

**Significance:** Why cerebral arteries become increasingly winding with age remains incompletely understood. We present a minimal mechanical framework in which constrained arterial elongation generates compression and leads to buckling. Bending relieves compression, while continued elongation drives further deformation. The model predicts that arterial bending stiffness and surrounding-tissue support determine a wavelength proportional to vessel diameter. Imaging of healthy adults reveals higher arterial excess-length ratios at older ages and supports predictions for wavelength scaling with vessel diameter and the wavelength-to-diameter ratio. These findings link vascular aging to growth-driven instability and pattern formation. Beyond the brain, the model may be extended to predict how vessel properties, wall geometry, and mechanical conditions shape healthy and pathological arteries.

## 1 Introduction

Straight arterial paths facilitate efficient blood transport to distal organs and tissues (1). However, arterial paths must also accommodate the anatomy of the body and individual organs. Cerebral arteries, including major arteries and their cortical branches, generally course along the brain surface, giving rise to fine branches that penetrate and supply the brain tissue (2). The human cerebrum is characterized by numerous ridges (gyri) and grooves (sulci), as well as deep fissures such as the Sylvian and interhemispheric fissures. Arteries thus follow complex paths along the surface and within these sulci and fissures (3, 4). Consequently, these vascular trajectories complicate both the three-dimensional (3D) analysis of arterial geometry and the interpretation of the results.

Vascular geometry also plays a crucial role in hemodynamics, and geometric alterations can contribute to vascular pathologies. Bends in a tortuous artery alter local blood flow. As a result, wall shear stress (the frictional force per unit area that blood exerts on the vessel wall) differs between the inner and outer sides of a bend (1). Disturbed wall shear stress is associated with arterial wall remodeling and the development of atherosclerosis (5, 6). In clinical practice, increased cerebral arterial tortuosity is frequently observed in older adults (Figure 1) (7–10). Greater cerebral arterial tortuosity has also been associated with several cerebrovascular diseases, including intracranial atherosclerosis, ischemic stroke, and cerebral aneurysm, as well as with Alzheimer’s disease (11–13). Aging, hypertension, and other vascular risk factors, as well as genetic predisposition, have been identified as potential contributors to vascular tortuosity (1, 14).

**Figure 1.**
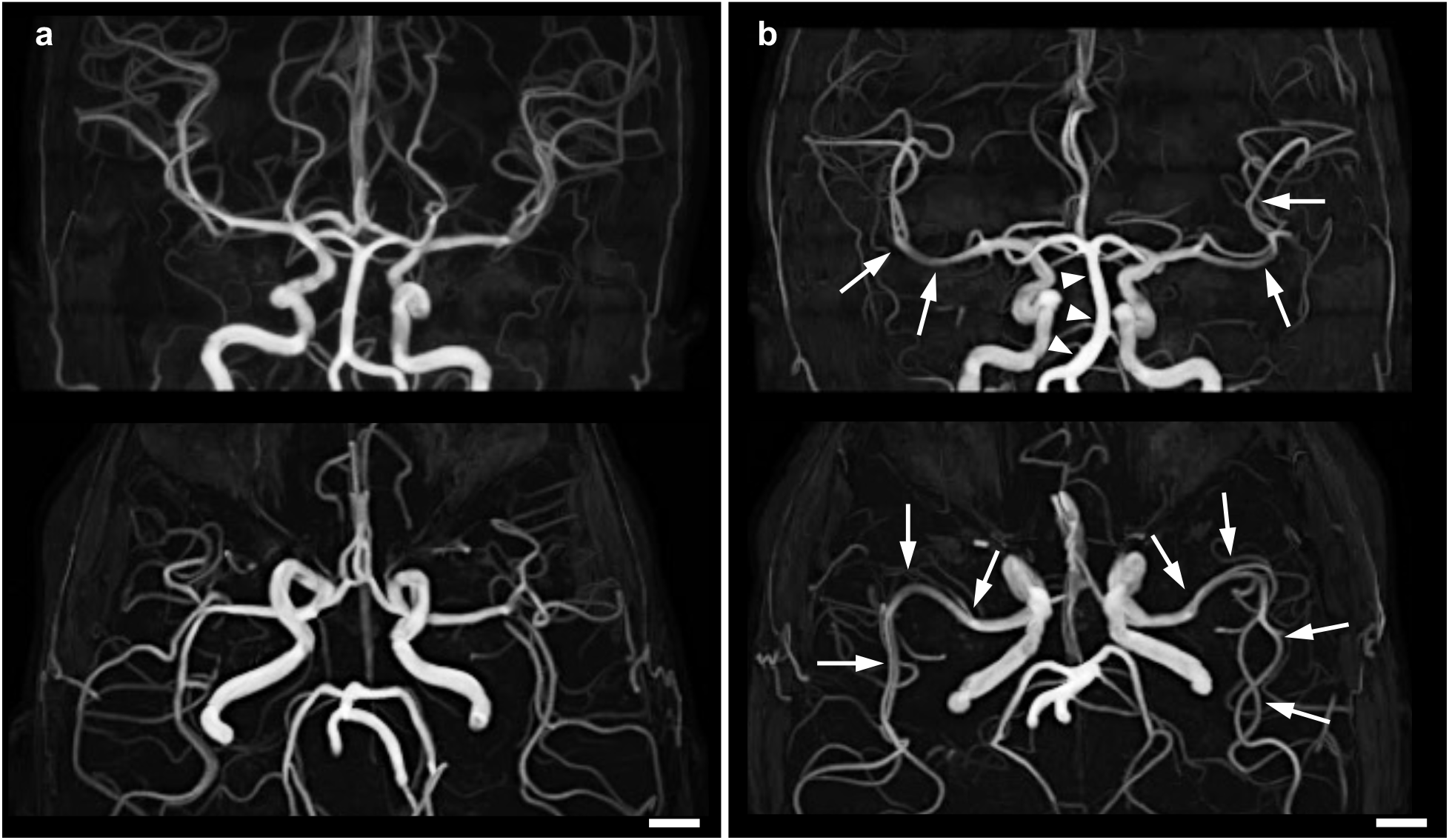
Maximum intensity projection images of magnetic resonance angiography in representative healthy adults. Top row: coronal views; bottom row: axial views. (a) 24-year-old participant; (b) 70-year-old participant. Compared with the younger participant, the older participant shows greater tortuosity of the middle cerebral arteries (arrows) and the basilar artery (arrowheads). The arterial waves have apparent wavelengths on the order of 10^1^ mm, as judged qualitatively from these projections. Scale bars, 10 mm.

Han developed theoretical models identifying elastic buckling under internal pressure as a possible mechanism of vascular tortuosity (15–18). These analyses predict critical internal pressures for buckling and corresponding buckling modes, and show that the critical pressure decreases as axial tension is reduced (1). Weiss et al. further showed in a computational model of mouse aortic growth and remodeling that mechanosensitive cellular responses can drive axial elongation. With the end-to-end distance fixed, these processes can gradually reduce axial tension and lead to progressive, persistent tortuosity in structurally vulnerable arteries (19). Age-related increases in the length of adult human cerebral arteries have been reported (10, 20, 21), but the associated *in vivo* axial loading is poorly characterized.

Building on this work, we developed a time-dependent one-dimensional (1D) buckling model of a cerebral artery supported by surrounding tissue. The artery is treated as an elastic column, and deformation is restricted to a plane approximating the local brain surface. The model describes how vessel elongation (an increase in the natural, stress-free length) at a fixed end-to-end distance generates compression and how lateral bending relieves it. We used mathematical analyses and numerical simulations to examine the onset of buckling and the subsequent evolution of wave amplitude during continued elongation and to predict the relationship between wavelength and vessel diameter. We evaluated the model’s geometric predictions using quantitative 3D measurements of arterial tortuosity, wavelength, and amplitude in human cerebral arteries reconstructed from magnetic resonance angiography (MRA).

## 2 Model

### 2.1 Euler–Bernoulli model with growth-induced axial compression

To describe age-related arterial tortuosity at the scale of the vessel centerline, we reduce the system to a 1D elastic column. This reduction is appropriate when the vessel diameter is small compared with the characteristic axial length scale of bending and the dominant deformation can be represented by a transverse displacement *h*(*x, t*) in a local plane. Here, *x* is the axial coordinate and *t* is time. We assume small slopes, linear elasticity, and Euler–Bernoulli kinematics, in which cross-sections initially normal to the centerline remain plane and normal to the deformed centerline. For a uniform column under a compressive axial force *P >* 0 and no distributed transverse load, the static transverse force balance per unit length is (22):

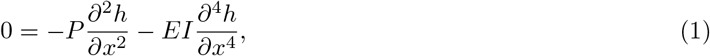

where *EI* is the effective flexural rigidity, with *E* denoting Young’s modulus of the arterial wall in a physiologically pressurized state and *I* the second moment of area. This classical Euler–Bernoulli balance provides the mechanical basis for the buckling model developed below.

We apply this framework to cerebral arteries. These vessels often follow the brain surface and course through fissures and sulci (3, 4) (Figure 2a). Locally, arteries lie between the brain surface and the overlying arachnoid membrane or between opposing brain surfaces within fissures and sulci (Figure 2b,c). Such anatomical surroundings can geometrically constrain surface-normal motion, so that the dominant tortuous deformation may be approximated locally as planar (Figure 2d). With aging, cerebral arteries undergo axial elongation (10, 20, 21). This elongation is also subject to anatomical constraints: proximal arteries are constrained at their dural entry sites (23, 24), whereas more distal segments can be partially confined by surrounding meningeal structures and by the complex geometry of the brain surface. When the increase in natural length is not fully accommodated by a corresponding increase in the projected span, a mechanical mismatch arises (19). In the present model, we prescribe an increase in natural length at a fixed projected span and represent the resulting mismatch as an effective compressive axial force *P* (*t*). We therefore model cerebral artery tortuosity as growth-driven buckling of an elastic column under anatomical confinement. Growth-induced compression and buckling under spatial constraint have also been studied using morphoelastic rod models (25, 26).

**Figure 2.**
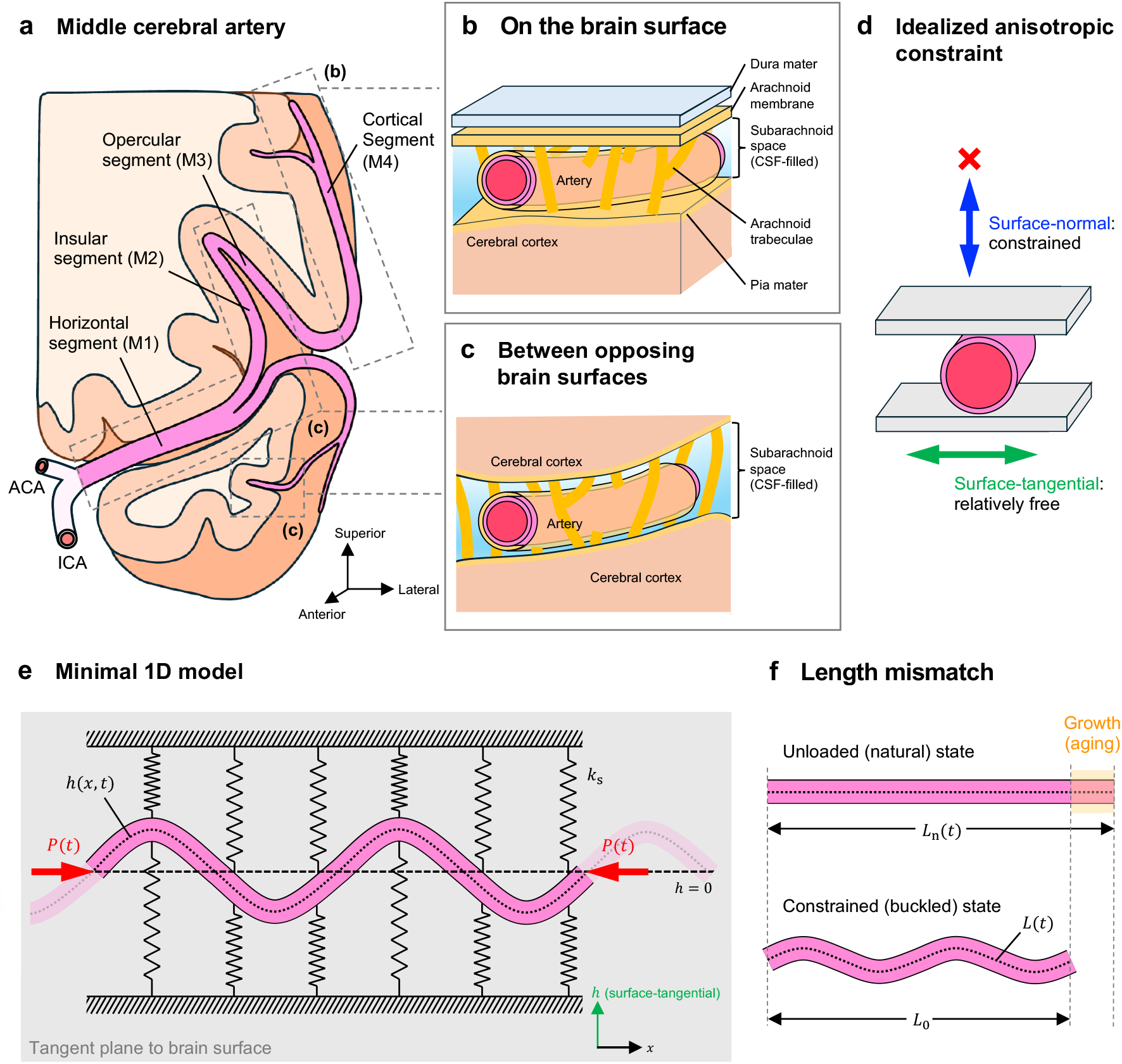
Anatomical basis and formulation of the surface-constrained 1D buckling model. (a) Schematic course of the middle cerebral artery, which runs along the brain surface within the subarachnoid space. Dashed boxes indicate representative regions shown in (b) and (c). (b) An artery between the brain surface and the overlying arachnoid membrane. (c) An artery between opposing brain surfaces. Arachnoid trabeculae are illustrated as local supporting structures in (b) and (c). (d) Idealized anisotropic constraint: surface-normal displacement is assumed to be restricted, whereas surface-tangential displacement perpendicular to the vessel axis is less geometrically constrained. (e) Minimal 1D model of an elastic column in the local tangent plane to the brain surface. *h*(*x, t*), surface-tangential transverse displacement of the vessel centerline (dotted line) from *h* = 0 (dashed line); *P* (*t*), effective compressive axial force. Hatched lines and springs represent the surrounding tissue as a linear elastic foundation of stiffness *k*_s_. The schematic depicts interior buckling with end effects neglected; the ends do not represent physical fixation points. (f) Growth-induced length mismatch. *L*_n_(*t*), natural (axially unloaded) length, which increases with aging; *L*_0_, fixed projected span; *L*(*t*), centerline arc length in the buckled state. Shading indicates the growth-induced length mismatch *L*_n_(*t*) −*L*_0_, accommodated by axial compression and by buckling. *1D, one-dimensional; ACA, anterior cerebral artery; ICA, internal carotid artery; CSF, cerebrospinal fluid*.

On the slow timescale of age-related arterial elongation, both axial and transverse inertial effects are neglected. The transverse dynamics are modeled as an overdamped force balance per unit length. The model also does not resolve blood flow or pulsatile fluid–structure interactions. The surrounding tissue is represented as a linear elastic foundation, whose restoring force per unit length is proportional to the local displacement *h* (17, 27) (Figure 2e). Incorporating effective damping and this restoring force into the static Euler–Bernoulli equation, Eq. 1, yields the governing equation for the slow time evolution of the transverse displacement:

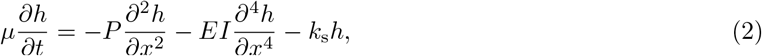

where *µ* is the effective damping coefficient and *k*_s_ is the foundation stiffness of the surrounding tissue. The cross-sectional geometry and mechanical coefficients are held fixed.

To couple natural-length elongation with transverse deformation, we determine *P* (*t*) from geometric compatibility between the time-dependent natural length *L*_n_(*t*) and the deformed centerline arc length *L*(*t*). Here, *L*(*t*) is the model counterpart of the measured centerline arc length *L*, and *L*_0_ denotes the projected span in the model (Figure 2f). To examine the mechanical consequences of growth-induced length mismatch, the reference model omits explicit internal pressure loading. Similarly, although arteries are normally axially prestretched *in vivo* (1), we adopt a straight reference configuration without initial axial stretch (*L*_n_(0) = *L*_0_). Under the small-slope approximation, |*∂h/∂x*|*≪*1, we approximate the deformed arc length 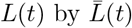, retaining the leading-order geometric nonlinearity:

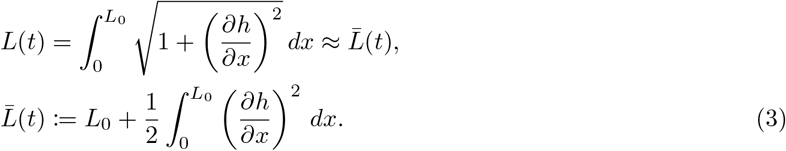

Using 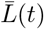, the effective compressive axial strain is

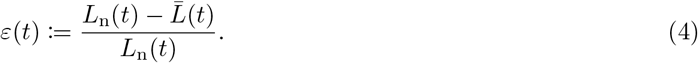

Assuming linear elasticity in the axial direction, *P* (*t*) is formulated as

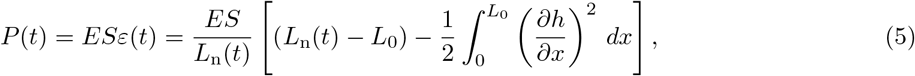

where *S* is the cross-sectional area of the vessel wall. The quadratic geometric term retained here is consistent with the von Kármán approximation (28).

Substituting the axial-force relation (Eq. 5) into the governing equation (Eq. 2) yields the equivalent nonlocal nonlinear evolution equation:

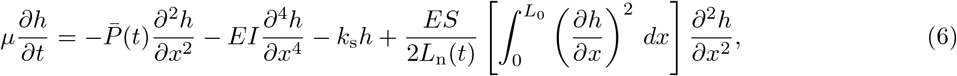

where 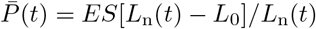 is the growth-induced axial force in the straight configuration (*h* = 0) obtained from Eq. 5. The last term in Eq. 6 constitutes the leading-order geometric nonlinearity associated with the deformation-induced reduction of the effective compressive force. Increasing transverse deformation reduces the effective axial compression, thereby acting as a stabilizing mechanism that limits wave amplification and permits a finite-amplitude post-buckling state. Continued elongation can drive further deformation. The nonlocal coupling between axial force and transverse deformation has the same mathematical structure as that in the Woinowsky-Krieger model (28).

An extended model incorporating internal pressure loading and initial axial stretch is described in *SI Appendix*, Section S1.1. In the nomenclature (Table 1 and *SI Appendix*, Table S1), superscripts (·)^sim^ and (·)^img^ denote simulation-derived quantities and quantities of the MRA-imaged arteries, respectively, and are omitted when the context is unambiguous.

**Table 1.**
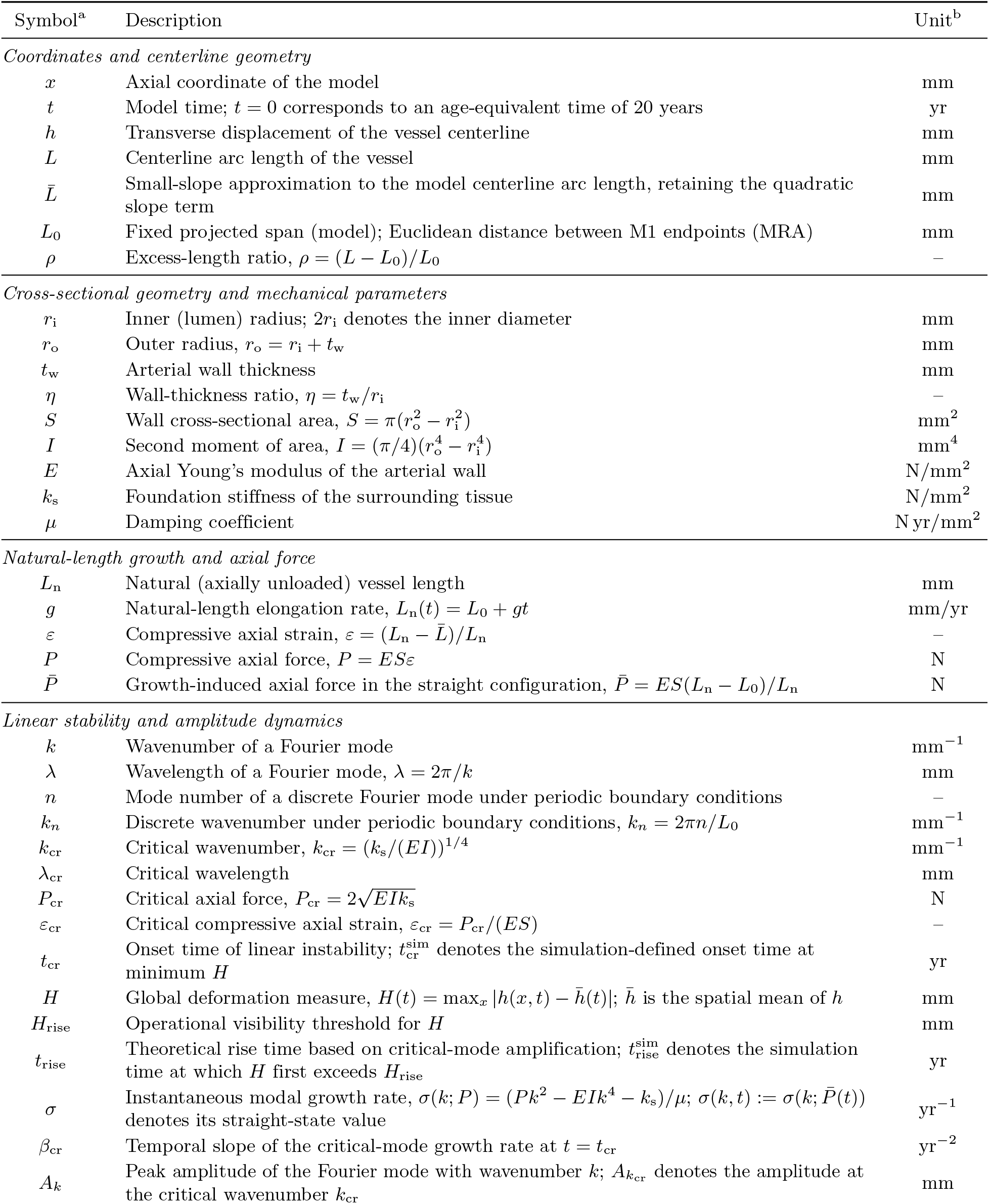
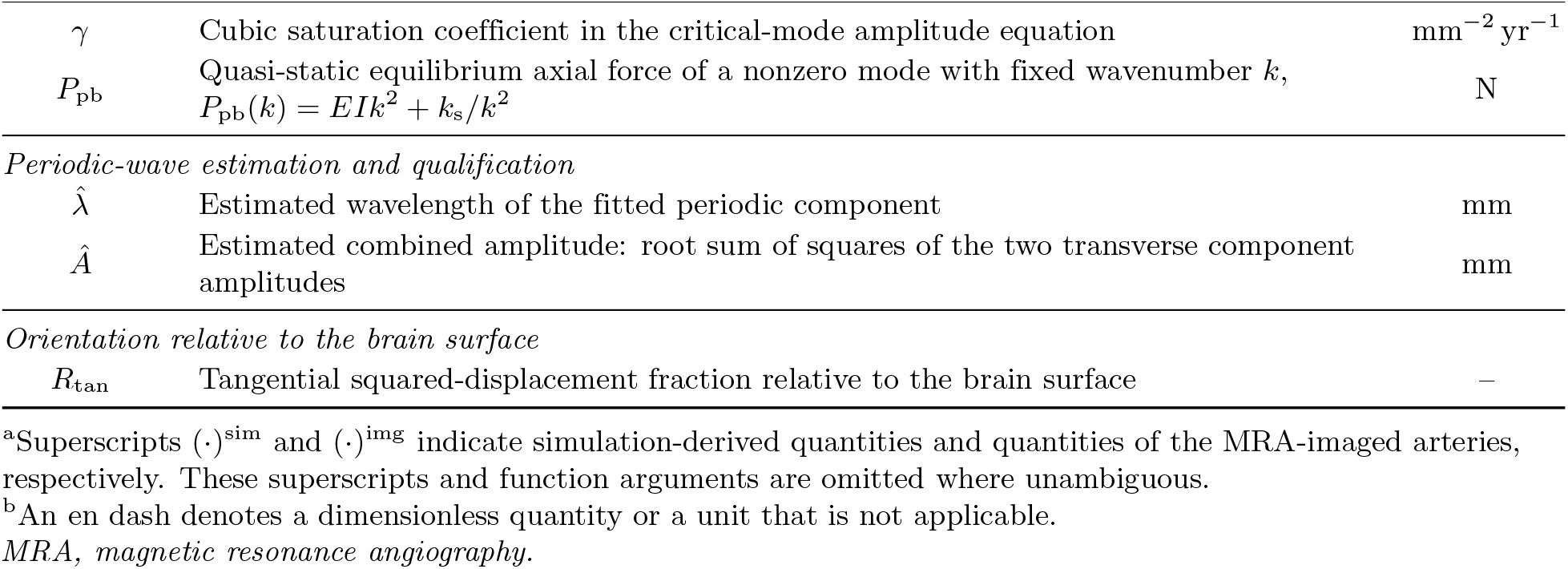
List of variables and parameters.

| Symbol<sup>a</sup> | Description | Unit<sup>b</sup> |
|---|---|---|
| *Coordinates and centerline geometry* |  |  |
| x | Axial coordinate of the model | mm |
| t | Model time; t = 0 corresponds to an age-equivalent time of 20 years | yr |
| h | Transverse displacement of the vessel centerline | mm |
| L | Centerline arc length of the vessel | mm |
| L̄ | Small-slope approximation to the model centerline arc length, retaining the quadratic slope term | mm |
| L<sub>0</sub> | Fixed projected span (model); Euclidean distance between M1 endpoints (MRA) | mm |
| ρ | Excess-length ratio, ρ = (L − L<sub>0</sub>)/L<sub>0</sub> | – |
| *Cross-sectional geometry and mechanical parameters* |  |  |
| r<sub>i</sub> | Inner (lumen) radius; 2r<sub>i</sub> denotes the inner diameter | mm |
| r<sub>o</sub> | Outer radius, r<sub>o</sub> = r<sub>i</sub> + t<sub>w</sub> | mm |
| t<sub>w</sub> | Arterial wall thickness | mm |
| η | Wall-thickness ratio, η = t<sub>w</sub>/r<sub>i</sub> | – |
| S | Wall cross-sectional area, S = π(r<sub>o</sub><sup>2</sup> − r<sub>i</sub><sup>2</sup>) | mm<sup>2</sup> |
| I | Second moment of area, I = (π/4)(r<sub>o</sub><sup>4</sup> − r<sub>i</sub><sup>4</sup>) | mm<sup>4</sup> |
| E | Axial Young’s modulus of the arterial wall | N/mm<sup>2</sup> |
| k<sub>s</sub> | Foundation stiffness of the surrounding tissue | N/mm<sup>2</sup> |
| μ | Damping coefficient | N yr/mm<sup>2</sup> |
| *Natural-length growth and axial force* |  |  |
| L<sub>n</sub> | Natural (axially unloaded) vessel length | mm |
| g | Natural-length elongation rate, L<sub>n</sub>(t) = L<sub>0</sub> + gt | mm/yr |
| ε | Compressive axial strain, ε = (L<sub>n</sub> − L̄)/L<sub>n</sub> | – |
| P | Compressive axial force, P = ESε | N |
| P̄ | Growth-induced axial force in the straight configuration, P̄ = ES(L<sub>n</sub> − L<sub>0</sub>)/L<sub>n</sub> | N |
| *Linear stability and amplitude dynamics* |  |  |
| k | Wavenumber of a Fourier mode | mm<sup>−1</sup> |
| λ | Wavelength of a Fourier mode, λ = 2π/k | mm |
| n | Mode number of a discrete Fourier mode under periodic boundary conditions | – |
| k<sub>n</sub> | Discrete wavenumber under periodic boundary conditions, k<sub>n</sub> = 2πn/L<sub>0</sub> | mm<sup>−1</sup> |
| k<sub>cr</sub> | Critical wavenumber, k<sub>cr</sub> = (k<sub>s</sub>/(EI))<sup>1/4</sup> | mm<sup>−1</sup> |
| λ<sub>cr</sub> | Critical wavelength | mm |
| P<sub>cr</sub> | Critical axial force, $P_\mathrm{cr} = 2\sqrt{EIk_\mathrm{s}}$ | N |
| ε<sub>cr</sub> | Critical compressive axial strain, ε<sub>cr</sub> = P<sub>cr</sub>/(ES) | – |
| t<sub>cr</sub> | Onset time of linear instability; t<sub>cr</sub><sup>sim</sup> denotes the simulation-defined onset time at minimum H | yr |
| H | Global deformation measure, H(t) = max<sub>x</sub> \|h(x, t) − h̄(t)\|; h̄ is the spatial mean of h | mm |
| H<sub>rise</sub> | Operational visibility threshold for H | mm |
| t<sub>rise</sub> | Theoretical rise time based on critical-mode amplification; t<sub>rise</sub><sup>sim</sup> denotes the simulation time at which H first exceeds H<sub>rise</sub> | yr |
| σ | Instantaneous modal growth rate, σ(k; P) = (Pk<sup>2</sup> − EIk<sup>4</sup> − k<sub>s</sub>)/μ; σ(k, t) := σ(k; P̄(t)) denotes its straight-state value | yr<sup>−1</sup> |
| β<sub>cr</sub> | Temporal slope of the critical-mode growth rate at t = t<sub>cr</sub> | yr<sup>−2</sup> |
| A<sub>k</sub> | Peak amplitude of the Fourier mode with wavenumber k; A<sub>k<sub>cr</sub></sub> denotes the amplitude at the critical wavenumber k<sub>cr</sub> | mm |
| γ | Cubic saturation coefficient in the critical-mode amplitude equation | mm<sup>−2</sup> yr<sup>−1</sup> |
| P<sub>pb</sub> | Quasi-static equilibrium axial force of a nonzero mode with fixed wavenumber k,<br>P<sub>pb</sub>(k) = EIk<sup>2</sup> + k<sub>s</sub>/k<sup>2</sup> | N |
| *Periodic-wave estimation and qualification* |  |  |
| λ̂ | Estimated wavelength of the fitted periodic component | mm |
| Â | Estimated combined amplitude: root sum of squares of the two transverse component amplitudes | mm |
| *Orientation relative to the brain surface* |  |  |
| R<sub>tan</sub> | Tangential squared-displacement fraction relative to the brain surface | – |

<sup>a</sup>Superscripts (·)<sup>sim</sup> and (·)<sup>img</sup> indicate simulation-derived quantities and quantities of the MRA-imaged arteries, respectively. These superscripts and function arguments are omitted where unambiguous.
<sup>b</sup>An en dash denotes a dimensionless quantity or a unit that is not applicable.
*MRA, magnetic resonance angiography.*

## 3 Results

### 3.1 Numerical simulations of the model

We examined the dynamical behavior of the proposed growth-driven buckling model using numerical simulations of Eq. 6. The parameters used in the simulations are summarized in Table 2. The spatiotemporal evolution is illustrated by the representative vessel profiles and the space–time map (Figure 3a,b). The corresponding coupled evolution of the axial force, global deformation, and modal amplitudes revealed three successive dynamic stages: pre-buckling, super-exponential growth, and slow growth (Figure 3c,d).

**Table 2.** M1-informed parameters used for analytical and numerical analyses.

| Symbol | Description | Value | Source |
|---|---|---|---|
| *Anatomical and material parameters* |  |  |  |
| *r*<sub>i</sub> | Inner (lumen) radius (mm) | 1.6 | Young-adult MRA data<sup>a</sup> |
| *t*<sub>w</sub> | Arterial wall thickness (mm) | 0.2 | Literature-based (47) |
| *E* | Axial Young's modulus (N/mm<sup>2</sup>) | 2.0 | Literature-based (29) |
| *Effective model parameters* |  |  |  |
| *k*<sub>s</sub> | Foundation stiffness (N/mm<sup>2</sup>) | 0.001 | Theoretical estimate (SI, S2.1) |
| *μ* | Damping coefficient (N yr/mm<sup>2</sup>) | 0.005 | Model-based reference value (SI, S2.2) |
| *g* | Natural-length elongation rate (mm/yr) | 0.35 | MRA-informed reference value (SI, S2.2) |
| *Derived geometric parameters* |  |  |  |
| *r*<sub>o</sub> | Outer radius, *r*<sub>o</sub> = *r*<sub>i</sub> + *t*<sub>w</sub> (mm) | 1.8 | – |
| *I* | Second moment of area, *I* = (π/4)(*r*<sub>o</sub><sup>4</sup> − *r*<sub>i</sub><sup>4</sup>) (mm<sup>4</sup>) | ≈ 3.10 | – |
| *S* | Wall cross-sectional area, *S* = π(*r*<sub>o</sub><sup>2</sup> − *r*<sub>i</sub><sup>2</sup>) (mm<sup>2</sup>) | ≈ 2.14 | – |
| *Numerical simulation settings* |  |  |  |
| *L*<sub>0</sub> | Fixed projected span (computational domain length) (mm) | 100 | Model setting |
| *t*<sub>max</sub> | Simulation time horizon (yr) | 60 | Adult-life simulation horizon |
| *N* | Number of spatial grid points | 1000 | – |
| Δ*t* | Time step (yr) | 0.01 | – |
| – | Initial arc length increment (% of *L*<sub>0</sub>) | 0.01 | – |
| – | Random seed | 1 | – |
| – | Output interval (steps) | 10 | – |

<sup>a</sup>The inner radius *r*<sub>i</sub> was set from the median M1 inner radius measured on the MRA data in young adults (< 35 years).
*MRA, magnetic resonance angiography.*

**Figure 3.**
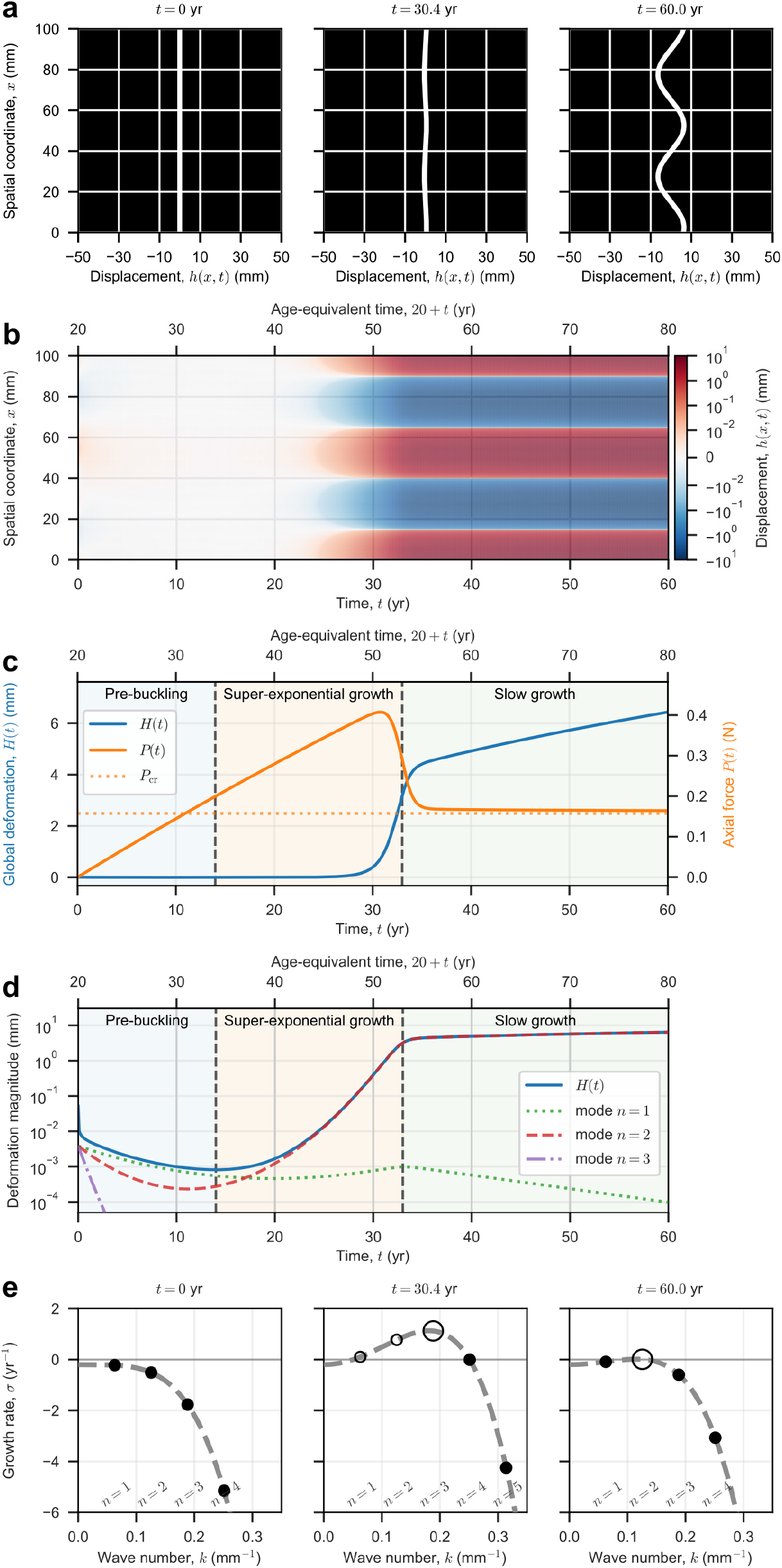
Numerical simulations of growth-driven buckling dynamics. (a)Representative configurations of the displacement field *h*(*x, t*) at selected time points, showing the temporal development of tortuosity. The left panel shows the initial configuration at *t* = 0 yr. The middle panel shows the configuration at *t* = 30.4 yr, the first time at which the global deformation measure *H*(*t*) exceeded the operational visibility threshold *H*_rise_ = 0.5 mm. The right panel shows the final configuration at *t* = 60 yr. (b)Spatiotemporal evolution of *h*(*x, t*), illustrating the emergence and growth of deformation over time. Displacement is shown using a symmetric logarithmic color scale. The lower axis shows model time *t*, and the upper axis shows age-equivalent time 20 + *t*, with *t* = 0 corresponding to 20 years of age. (c) Time evolution of *H*(*t*) and the axial force *P* (*t*). *P* (*t*) (orange line) initially increased due to elongation, reached a peak, and then relaxed toward a lower post-buckling value, while *H*(*t*) (blue line) continued to increase. The horizontal dotted orange line indicates the critical axial force from the linear stability analysis, *P*_cr_. Colored backgrounds and vertical dashed lines in (c) and (d) indicate the pre-buckling, super-exponential growth, and slow-growth stages. The first boundary marks the amplitude-based onset at *t* = 14.0 yr, when *H*(*t*) reached its minimum. The second boundary, at *t* = 33.0 yr, was defined as the time after the peak of *d*(log *H*)*/dt* at which *d*^2^(log *H*)*/dt*^2^ attained its minimum. (d) Time evolution of *H*(*t*) and the amplitudes of the selected Fourier modes *n* = 1, 2, and 3 on a logarithmic scale. (e) Instantaneous growth-rate spectra at the representative times shown in (a). The dashed curve indicates the continuum dispersion relation *σ*(*k*; *P*) evaluated using the simulated *P* (*t*), whereas circles indicate the discrete wavenumbers *k*_*n*_ = 2*πn/L*_0_ allowed by the periodic boundary conditions. Filled circles denote stable modes with *σ <* 0, and open circles denote unstable modes with *σ >* 0. The largest open circle identifies the discrete mode with the highest positive growth rate at each time.

#### Pre-buckling stage

The pre-buckling stage was defined as the interval before the amplitude-based simulation onset, 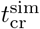, at which the global deformation measure 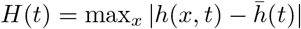reached its minimum. Here, 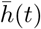 is the spatial mean of *h*(*x, t*). During this stage, continued elongation increased the axial force *P* (*t*). However, transverse deformation did not develop, and each modal amplitude remained small (Figure 3c,d).

#### Super-exponential growth stage

After the amplitude-based onset, *H*(*t*) initially showed an increasing slope on a log-linear plot, indicating super-exponential growth (Figure 3d). The first modal instability occurred before the *H*(*t*) onset. The amplitude of the *n* = 2 mode (*λ*^sim^ = 50.0 mm) reached its minimum at *t* = 11.2 yr, while *H*(*t*) reached its minimum later at 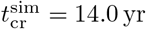 (an age-equivalent time of 34.0 years). This offset reflects the combined contributions of growing and decaying modes to *H*(*t*). Because *H*(*t*) initially remained small despite this growth, we defined the rise time, 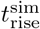, as the first time at which *H*(*t*) exceeded the operational visibility threshold *H*_rise_ = 0.5 mm. This threshold was first exceeded at 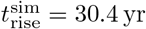, corresponding to an age-equivalent time of 50.4 years. Thus, due to the initially slow yet accelerating growth of *H*(*t*), visually apparent deformation emerged much later than either onset. During this stage, *P* (*t*) continued to increase under progressive elongation and reached a transient peak near its end (Figure 3c).

For the dominant *n* = 2 mode, the wavelength-to-outer-diameter ratio, *λ/*(2*r*_o_), was 13.9, supporting the slenderness assumption underlying the Euler–Bernoulli theory.

#### Slow-growth stage

Following the super-exponential growth stage, the system entered a slow-growth post-buckling stage. The axial force *P* (*t*) relaxed toward a post-buckling level, while the global deformation measure *H*(*t*) continued to increase (Figure 3c). This behavior indicates that the additional imposed elongation was mainly accommodated by lateral deformation rather than a further increase in axial force. In the reference simulation, once deformation became visually apparent, the first unstable mode, *n* = 2, remained dominant throughout the subsequent evolution (Figure 3b,d).

However, because the small-slope condition was no longer satisfied late in the slow-growth stage, the simulated deformation during this period should be regarded as a qualitative extrapolation of this model rather than as a quantitative prediction (*SI Appendix*, Section S3.1).

### 3.2 Mathematical analysis of the model

**Linear stability analysis**

We conducted a linear stability analysis to predict the onset of instability and the characteristic wavelength of vessel buckling (*SI Appendix*, Section S4.1). For a spatially uniform axial force *P*, the instantaneous growth rate of a Fourier mode with wavenumber *k* is

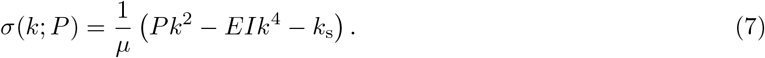

This dispersion relation distinguishes growing (*σ >* 0) from decaying (*σ <* 0) modes. The curves in Figure 3e were evaluated using the actual axial force *P* (*t*) from the nonlinear simulation. In contrast, linearizing about the straight state gives 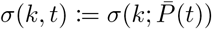. By imposing the condition that the maximum of *σ*(*k, t*) over continuous wavenumbers first reaches zero, the analysis identifies the critical wavelength *λ*_cr_ and the corresponding critical axial force *P*_cr_ at the onset of instability:

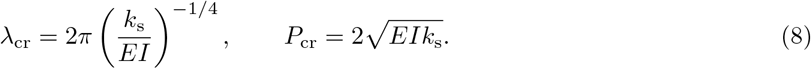

The corresponding critical wavenumber is *k*_cr_ = 2*π/λ*_cr_. With an inner (lumen) radius *r*_i_ and a wall-thickness ratio *η* = *t*_w_*/r*_i_, the critical wavelength in Eq. 8 can be written as

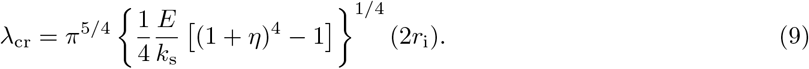

These expressions indicate that the balance between arterial bending stiffness and surrounding-tissue support selects a characteristic wavelength proportional to the inner diameter 2*r*_i_ at fixed *η* and *k*_s_*/E*. The dependence on these ratios is relatively weak, approaching fourth-root scaling for *η* in the thin-wall limit (*η≪*1) and following an inverse fourth-root law for *k*_s_*/E*.

The corresponding critical axial strain for the straight state is

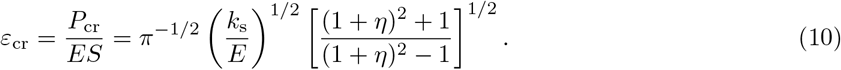

With the projected span fixed, this threshold characterizes the elongation mismatch required for the straight state to become unstable.

Using the M1-informed parameters summarized in Table 2, the predicted critical wavelength is *λ*_cr_ ≈ 55.7 mm. This wavelength is approximately 17.4 times the inner diameter (2*r*_i_), indicating broad bends rather than closely spaced waves. Under periodic boundary conditions with *L*_0_ = 100 mm, the first unstable admissible mode is *n* = 2, with a wavelength of 50 mm, matching the dominant post-buckling mode in the numerical simulations (Figure 3b,d). This wavelength scale is qualitatively consistent with the apparent arterial wavelengths on the order of 10^1^ mm seen in the MRA projections (Figure 1). The median M1 Euclidean distance (18.0 mm; *SI Appendix*, Table S2) was shorter than the simulated half-wavelength (25.0 mm). At this length scale, M1 could show a single broad C-shaped or partial S-shaped bend rather than repeated short waves, consistent with reported age-related shifts toward more tortuous M1 configurations with C- or S-shaped projections (7, 9, 30).

The predicted instability onset is *t*_cr_ ≈ 10.9 yr for continuous wavenumbers and approximately 11.2 yr for the admissible *n* = 2 mode. The time-dependent linear analysis also explains the delay between the onset of instability and visible deformation in the simulations. Near the critical time (*t* ≥ *t*_cr_), the amplitude of the critical mode is approximated as

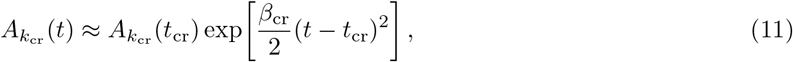

where *β*_cr_ *>* 0 is the temporal slope of the critical-mode growth rate at *t* = *t*_cr_. This accelerating amplification from a minute amplitude provides a theoretical basis for the delay before visible deformation and the early part of the super-exponential growth stage in the simulations (Figure 3c,d).

For an approximate comparison with the simulations, we used the ratio of the operational visibility threshold *H*_rise_ = 0.5 mm to the minimum simulated *H*(*t*) as an effective amplification factor in the continuous-wavenumber theory (*SI Appendix*, Section S4.1). This yielded *t*_rise_ ≈ 30.0 yr, in reasonable agreement with the simulation-defined rise time 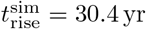.

#### Weakly nonlinear analysis

To determine how force relaxation caused by increasing vessel tortuosity affects further amplitude growth near the onset of instability, we performed a weakly nonlinear analysis (*SI Appendix*, Section S4.2). Under a single-mode approximation, projecting Eq. 6 onto the critical mode reduces the dynamics to an amplitude equation of the form

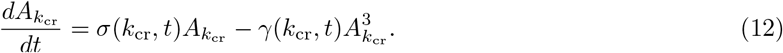

Here, 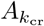 is the peak amplitude of the critical mode, *σ*(*k*_cr_, *t*) is the linear growth rate obtained by evaluating Eq. 7 at 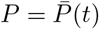, and *γ*(*k*_cr_, *t*) *>* 0 is the cubic saturation coefficient arising from the nonlinear geometric constraint. With *σ* and *γ* treated as time-independent parameters, the corresponding equation for a signed amplitude has the normal form of a supercritical pitchfork bifurcation (31).

The nontrivial equilibrium branch satisfies

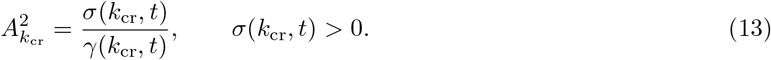

Along this equilibrium branch near the onset, the small-slope approximation relates the modal amplitude to the excess-length ratio *ρ* = (*L* − *L*_0_)*/L*_0_:

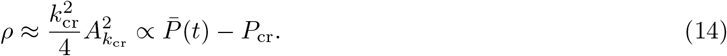

This result directly links the supercritical pitchfork bifurcation to the geometrically measurable quantity *ρ*. As 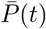 increases with vessel growth, the equilibrium amplitude rises along this quasi-static branch.

The weakly nonlinear analysis also provides a mechanical interpretation of the force peak observed in the simulations (Figure 3c). Under the single-mode approximation, the axial force is approximated as

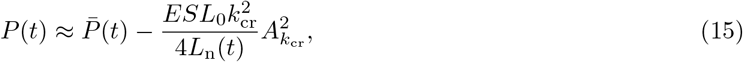

showing that the growing buckling amplitude converts part of the growth-induced length mismatch into additional arc length, thereby reducing *P* (*t*) relative to the instantaneous straight-state force 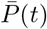. At the force peak, the rate of geometric force relaxation balances the rate of increase in 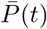.

#### Post-buckling quasi-static equilibrium and modal dynamics under continued growth

To explain why post-buckling deformation continued to increase as the axial force approached a near-critical value in the simulations (Figure 3c), we performed a quasi-static analysis and examined the corresponding modal dynamics (*SI Appendix*, Section S4.3). For a nonzero single-mode equilibrium with fixed wavenumber *k*, the quasi-static force balance gives

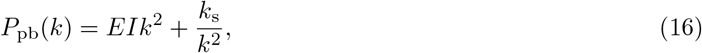

where *P*_pb_(*k*) denotes the equilibrium axial force associated with wavenumber *k*. This force is independent of the mode amplitude and natural length; thus, along the quasi-static equilibrium branch, additional growth is accommodated by an increase in amplitude rather than in axial force. Evaluating this expression at *k* = *k*_cr_ gives *P*_pb_(*k*_cr_) = *P*_cr_.

Away from equilibrium, the peak amplitude of mode *k, A*_*k*_(*t*), evolves according to

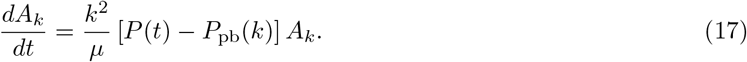

This amplitude equation shows that a mode grows when the actual axial force exceeds its equilibrium force. Late in the slow-growth stage in the reference simulation, *P* (*t*) remained slightly above the equilibrium force of the dominant *n* = 2 mode, allowing its amplitude to continue growing slowly (Figure 3c,d). The equilibrium force of this discrete mode is slightly higher than *P*_cr_ = *P*_pb_(*k*_cr_), the minimum critical axial force over continuous wavenumbers. Because the first unstable mode has the lowest equilibrium force among the admissible modes, other modes cannot grow while *P* (*t*) remains near this level. This explains why the first unstable mode remained dominant in the slow-growth stage (Figure 3b,d).

### 3.3 Geometric analysis of MRA data from healthy adults

#### Vascular elongation and tortuosity with aging in M1 segments

We analyzed 3-T MRA and structural T1-weighted images from 154 adults aged 20–81 years in the IXI (Information eXtraction from Images) Dataset (32). To assess whether age-related differences in arterial geometry are consistent with the model, we focused on the M1 segment of the middle cerebral artery (MCA; Figure 2a,c). Its relatively straight course in younger adults (7, 9, 30) (Figure 1) approximates the initially straight configuration assumed in the model. Accordingly, the Euclidean distance between its anatomically defined endpoints can serve as a geometric counterpart of the model span (see *SI Appendix*, Section S5.4 for the M1 endpoint definitions). We measured the centerline length *L*, Euclidean distance *L*_0_, and excess-length ratio *ρ* (Figure 4a–c). This cross-sectional MRA analysis included 349 anatomically defined M1 segments from 154 participants; segment-level morphological parameters are summarized in *SI Appendix*, Table S2.

**Figure 4.**
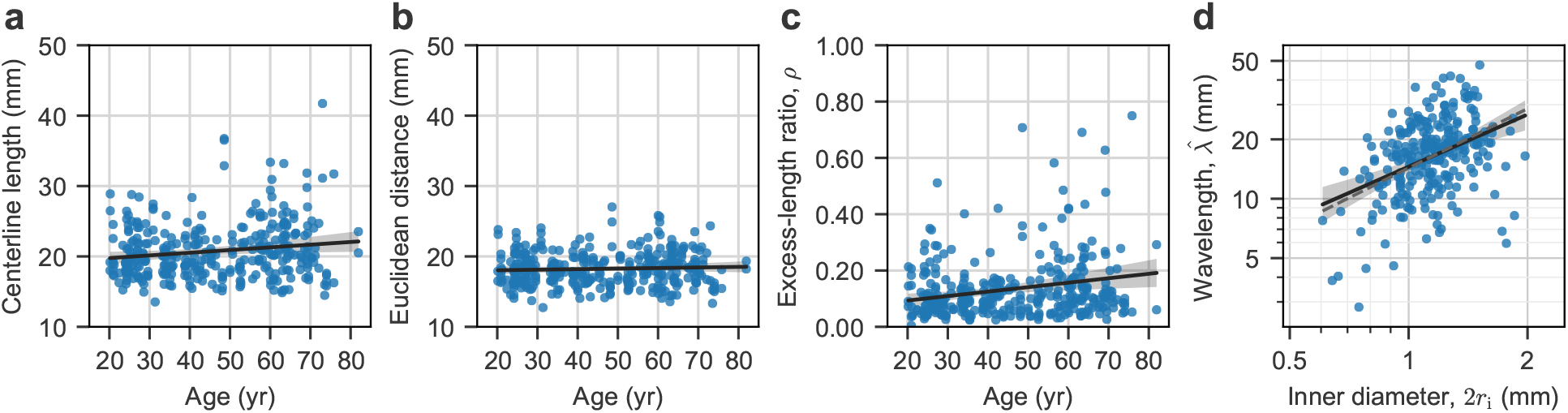
Geometric characteristics of middle cerebral arteries in healthy adults. (a–c) M1 morphological measurements versus age (*n* = 349 segments from 154 participants): (a) centerline length, (b) Euclidean distance, and (c) excess-length ratio *ρ*. (d) Estimated wavelength versus inner diameter on log–log axes in middle cerebral artery intervals with predominantly tangential periodic deviations (*n* = 275 intervals from 133 participants). The dashed line is a slope-1 reference passing through the means of the log-transformed values, representing the proportional scaling 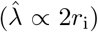 predicted by the idealized linear stability analysis. Black solid lines indicate univariable regression fits, and gray shaded bands indicate pointwise 95% confidence intervals calculated using participant-level cluster-robust standard errors.

In multivariable ordinary least-squares (OLS) regression analyses adjusting for demographic and anatomical covariates and using participant-level cluster-robust standard errors (*SI Appendix*, Table S3), older age was associated with greater M1 centerline length (*β* = 0.063 mm*/*yr; 95% confidence interval [CI], 0.031 to 0.095; *P* < 0.001). The Euclidean distance was also positively associated with age (*β* = 0.022 mm*/*yr; 95% CI, 0.004 to 0.039; *P* = 0.014), although the estimated regression coefficient was smaller than that for centerline length. The excess-length ratio *ρ* was greater in older participants (*β* = 0.0021 yr^*−*1^; 95% CI, 0.0009 to 0.0032; *P* < 0.001). These findings are qualitatively consistent with the increase in geometric excess length predicted by our model.

To explore whether the association between age and *ρ* followed a single linear trend across adulthood or showed a transition suggestive of buckling onset or subsequent accelerated amplification, we performed exploratory spline and piecewise linear regression analyses. The adjusted spline and its age derivative, *dρ/d*Age, suggested that the age association became more pronounced from early middle age onward (*SI Appendix*, Figure S1a,b). In the piecewise linear models, slope-change estimates were positive across all evaluated candidate breakpoints but imprecise (*SI Appendix*, Table S4). In participant-level cluster bootstrap analysis, the most frequently selected breakpoint was 35 years, but the selected breakpoints were distributed across the candidate range (*SI Appendix*, Figure S1c). These exploratory analyses did not identify a well-localized age breakpoint.

#### Periodic arterial waves as buckling-like morphology

We then extended the analysis to all analyzable MCA centerline intervals bounded by branch points or distal endpoints. We fitted a smooth quadratic baseline to represent the broad course of each interval. Using the Bishop frame (33) constructed along the fitted baseline, we extracted the transverse deviations of the centerline from the baseline (*SI Appendix*, Figure S2). We applied sinusoid-plus-trend fitting to these deviations to estimate the wavelength 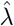 (Figure 4d) and the combined amplitude *Â* (the root sum of squares of the two transverse component amplitudes) of the fitted periodic component (*SI Appendix*, Section S5.5). Because the model assumes waves tangential to the brain surface, we selected intervals that had both an adequate periodic fit and predominantly tangential periodic deviations. This restriction was designed to minimize the extent to which the estimated wavelengths reflect surface-normal waves following brain-surface topography. We assessed two predictions of the model: that wavelength scales in proportion to the inner diameter 2*r*_i_ (Eq. 9) and that wave amplitude increases with age (Figure 3d). Equation 9 predicts the characteristic wavelength at buckling onset. In the reference simulation, the first unstable mode remained dominant in the slow-growth stage (Figure 3b,d), providing a rationale for comparing the onset prediction with observed arterial wavelengths.

Of 11,808 MCA centerline intervals from 154 participants, 275 (2.3%) from 133 participants met these selection criteria (*SI Appendix*, Figure S3). All selected intervals were from M2–M4. Among the 133 participants with included intervals, the median number of intervals per participant was 2 (interquartile range [IQR], 1–3). The median ellipticity, defined as the minor-to-major axis ratio of the ellipse traced by the fitted periodic deviation in Bishop transverse coordinates (*SI Appendix*, Figure S2d), was 0.20 (IQR, 0.08–0.34). This indicates that the periodic deviation was concentrated predominantly along one transverse direction, consistent with the single transverse displacement represented in our minimal buckling model.

In multivariable analysis (*SI Appendix*, Table S5), the estimated wavelength 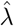 increased with the inner diameter on the log–log scale (*β* = 0.92; 95% CI, 0.61 to 1.24; *P* < 0.001). The 95% CI for the scaling exponent included 1, the value expected for the proportional wavelength–diameter scaling predicted by the model. The median wavelength-to-inner-diameter ratio, 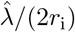, was 14.4 (IQR, 11.4–18.8); the ratio of approximately 17.4 predicted by the linear stability analysis (Eq. 9 with the parameter values in Table 2) lies within this IQR. In noise-free simulations with synthetic centerlines, the estimation and qualification procedure preserved proportional scaling when present and did not generate it when absent (*SI Appendix*, Table S6). As a supplementary analysis, we also examined the association between the wavelength-to-inner-diameter ratio and age. This association was not statistically significant after multivariable adjustment (estimated relative difference per year, 0.24%; 95% CI, −0.08% to 0.56%; *P* = 0.146). After adjustment for log-transformed wavelength, log-transformed inner diameter, and other covariates, we did not detect a positive association between age and wave amplitude (estimated relative difference per year, −0.42%; 95% CI, −0.86% to 0.02%; *P* = 0.064).

## 4 Discussion

We developed a growth-driven, surface-constrained buckling model of age-related cerebral arterial tortuosity and evaluated it through mathematical analyses, numerical simulations, and comparisons with MRA-derived geometric measurements. The model links progressive vessel elongation to an effective compressive force under partial spatial constraint and predicts a characteristic wavelength governed by arterial bending stiffness and surrounding-tissue support. In the cross-sectional MRA data, excess-length ratios were greater at older ages, and periodic waves exhibited diameter-dependent wavelength scaling. These geometric findings are consistent with the model’s predictions for post-buckling deformation and wavelength selection, supporting a buckling-based framework for interpreting age-related cerebral arterial tortuosity. However, an age-related increase in amplitude was not detected, leaving the predicted post-buckling amplitude growth unverified in our MRA data.

### 4.1 Buckling-based framework for vascular tortuosity formation

Buckling instability has been widely used to explain how growth-induced mechanical mismatch can convert slender or layered biological structures into stable geometric patterns rather than simply causing mechanical failure. Such mechanisms have been proposed to shape thin sheets and layered tissues, as in intestinal villification (34, 35), leaf morphogenesis (36), cortical gyrification (37), and fingerprint formation (38). Constrained elongation can generate curved or looped configurations in slender tubes, as in gut looping (39) and the elongated, sinusoidal tracheal tubes of *Drosophila* mutants (40).

In vascular biomechanics, Han formulated arterial tortuosity as a mechanical stability problem involving pressure-induced buckling, with the critical buckling pressure depending on axial stretch, wall properties, and surrounding-tissue support (15, 17, 18). Vascular models have also investigated how growth affects the critical buckling pressure (41, 42). Mechanobiological models incorporating mechanically regulated growth and remodeling have examined the initiation and progression of tortuosity under axial constraint (19, 43). More generally, morphoelastic rod models provide a mathematical framework for relating constrained growth to buckling and subsequent pattern formation (25, 26, 44). Building on this mathematical foundation and these vascular studies, we apply the concepts of buckling instability and pattern formation to age-related morphological changes in adult cerebral arteries by incorporating a time-dependent compressive force arising from vessel elongation under axial constraint (Eq. 5). The resulting nonlinear, nonlocal governing equation (Eq. 6) describes the time evolution of vessel deformation, allowing buckling onset, wavelength selection, and post-buckling amplitude growth to be examined analytically and numerically.

Although this study focused on vessel elongation as a primary trigger, the governing equation also allows for an analysis of how wall properties and surrounding-tissue support affect buckling susceptibility. In the present displacement-controlled framework, growth increases the natural length while the projected span remains fixed. The critical axial strain (Eq. 10) therefore characterizes the elongation mismatch required for buckling onset. Under these conditions, increasing *E* or *η* (i.e., wall stiffening and wall thickening) can lower the critical strain despite increasing the critical axial force (*SI Appendix*, Eq. S23). In contrast, a reduction in the foundation stiffness *k*_s_ lowers both the force and strain thresholds. In addition to vessel elongation (19), arterial growth and remodeling can involve wall thickening and changes in mechanical properties (6). The present model suggests that such changes may modulate susceptibility to growth-driven tortuosity.

Arterial tortuosity includes several phenotypes, such as curving or curling, angulation or kinking, looping, and spiral twisting (1). Among these phenotypes, our 1D buckling model provides a geometric interpretation of long-wavelength curving-type tortuosity, particularly in vessels with restricted displacement normal to anatomical surfaces. The diameter dependence predicted by Eq. 9, together with the weaker sensitivity to *η* and *k*_s_*/E* (a ratio of two stiffnesses that are difficult to quantify separately *in vivo*), may provide a mechanical explanation for the qualitative similarity of tortuous vessel shapes across widely differing calibers (45). Specifically, the scaling suggests that surface-constrained arteries can develop long-wavelength curving with broadly similar wavelength-to-diameter ratios across a wide range of calibers, potentially extending from retinal, superficial temporal, and coronary arteries to iliac arteries and the aorta.

### 4.2 Age-related arterial changes and clinical implications

The proposed buckling mechanism can be placed within a potential sequence of events linking aging to cerebrovascular disease. Arterial remodeling associated with aging and vascular risk factors may contribute to tortuosity formation (1, 14). Once formed, tortuosity can alter local flow patterns, wall shear stress, and intramural stress distributions (1). Such hemodynamic and mechanical changes may contribute to further alterations in vascular wall structure and function (5, 6), potentially creating a feedback loop between vascular geometry, hemodynamics, and wall remodeling (19) (Figure 5). The clinical relevance of tortuosity is supported by its reported associations with various cerebrovascular diseases (11–13). Our study provides a mechanical link between vessel elongation and the geometric changes that may subsequently affect hemodynamics and contribute to vascular pathology, while accounting for the effects of wall properties and surrounding-tissue support on buckling susceptibility. Because direct characterization of vascular tissue is often limited in living patients, this framework offers insight into disease mechanisms by relating imaging-derived vascular geometry to the mechanical processes underlying tortuosity.

**Figure 5.**
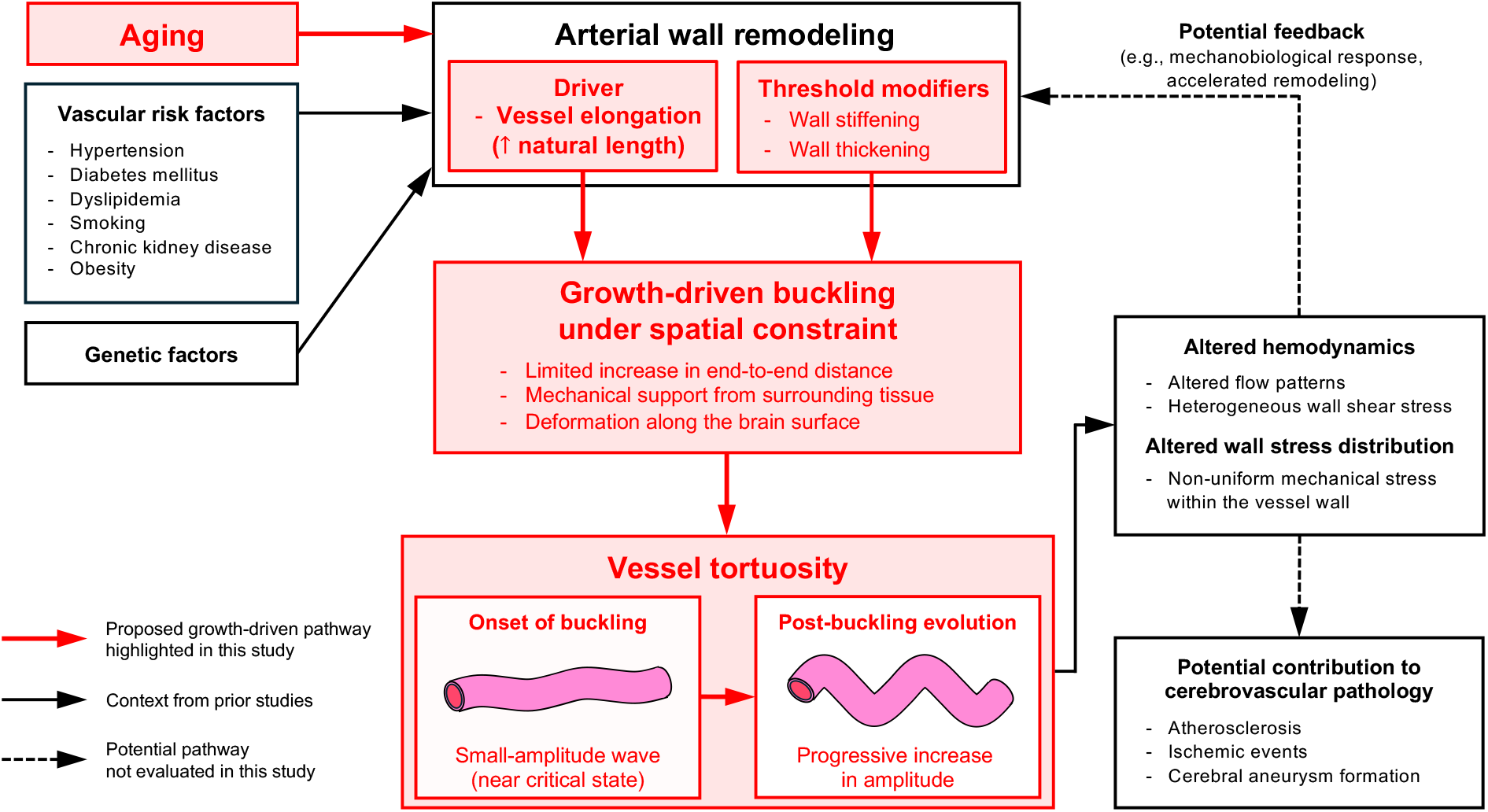
Proposed conceptual model for the initiation and development of arterial tortuosity. Red elements highlight the proposed growth-driven buckling pathway. The mechanical model predicts that a progressive increase in vessel natural length under spatial constraint can initiate buckling and drive post-buckling amplitude growth. Solid black arrows indicate relationships described in prior studies. Dashed arrows indicate potential feedback and downstream pathological consequences that were not evaluated in this study.

### 4.3 Limitations

Several limitations should be considered.

First, our theoretical model relies on simplified assumptions about vascular geometry and wall mechanics. It describes planar buckling of an initially straight, unbranched vessel, whereas real arteries follow complex 3D trajectories along the brain surface, may have pre-existing curvature, and interact with spatially heterogeneous tissue support. The model also does not represent possible remodeling of intrinsic curvature or of the reference configuration of tissue support. Natural-length elongation is prescribed without explicitly modeling the biological processes that produce it. These simplifications isolate the essential physical mechanism but may limit quantitative application to individual vessels.

Second, cross-sectional MRA cannot determine the timing of buckling onset or track subsequent amplitude growth within individual vessels. Distal cerebral arteries, with their small diameters and slow flow, are susceptible to flow-related signal loss on time-of-flight MRA and are less visible at older ages (46). Tortuosity can increase flow resistance (1), and any associated flow reduction may further impair depiction, potentially underrepresenting highly tortuous vessels. Our exploratory analysis showed lower total reconstructed vessel length and fewer detected intervals at older ages (*SI Appendix*, Figure S4). These findings raise the possibility of age-related selection bias in the periodic wave analysis of M2–M4 segments. Such bias could have made it harder to detect any increase in wave amplitude with age. In addition, the inner diameters of distal branches can approach the spatial resolution of time-of-flight MRA, so partial-volume effects may introduce error in diameter measurements.

Future studies should combine longitudinal imaging with model refinement to test the predicted onset and progression of tortuosity and evaluate the clinical relevance of the proposed mechanism. Experimental models in which arterial elongation can be manipulated would provide a more direct test of the proposed causal mechanism.

## 5 Conclusions

This study provides a biomechanical framework in which cerebral arteries become tortuous with age through buckling driven by vessel elongation under axial constraint. In cross-sectional MRA data from healthy adults, greater excess-length ratios at older ages and the wavelength–diameter scaling were consistent with the model’s predictions for post-buckling deformation and wavelength selection. Our model links vascular aging to growth-driven instability and pattern formation and provides a conceptual basis for understanding physiological and pathological changes in vascular geometry both within and beyond the brain.

## 6 Materials and Methods

### 6.1 Setting of numerical simulations

The numerical simulations were performed using Python 3.10. Because the aim was to analyze intrinsic wavelength selection and post-buckling dynamics in a locally repeating vessel segment, periodic boundary conditions were adopted.

Spatial derivatives were discretized using second-order central finite differences on a uniform periodic grid. A semi-implicit Euler scheme was used. At each time step, the axial force *P* (*t*) was first updated explicitly from the current displacement field via the geometric compatibility relation in Eq. 5. Subsequently, with *P* (*t*) held fixed for the duration of the time step, the displacement field was advanced implicitly using the resulting linear spatial operator associated with Eq. 2 to improve numerical stability. Simulations were conducted under the following conditions: projected span *L*_0_ = 100 mm, number of spatial grid points *N* = 1000, total simulation time *t*_max_ = 60 yr, and time step Δ*t* = 0.01 yr. Model time *t* = 0 corresponded to an age-equivalent time of 20 years. The growth-induced natural length was prescribed to increase linearly in time as *L*_n_(*t*) = *L*_0_ + *gt*, with a natural-length elongation rate of *g* = 0.35 mm*/*yr. The model parameters are summarized in Table 2. The initial configuration was a nearly straight centerline perturbed by a band-limited superposition of equal-amplitude Fourier modes. Modes with wavelengths shorter than the outer vessel diameter were excluded. Independent random phases were assigned to the retained modes, and the resulting profile was mean-subtracted. Finally, the profile was rescaled so that the initial value of the quadratic arc-length term 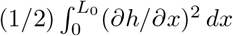 in the axial-force relation (Eq. 5) was 0.01% of the projected span.

### 6.2 MRA data analysis

#### Human participants and data acquisition

This study used the IXI Dataset, which contains magnetic resonance imaging (MRI) data from normal, healthy adults in London, UK (32). Because only anonymized, publicly available data were used, this study was outside the scope of the Japanese Ethical Guidelines for Medical and Biological Research Involving Human Subjects, and ethical approval was not required. All data were used in full compliance with the dataset’s usage policies and the Creative Commons Attribution-ShareAlike 3.0 (CC BY-SA 3.0) license.

We quantified cerebral artery tortuosity and examined its relationships with age and vessel diameter. We used a subset of the IXI Dataset, including MRA and structural T1-weighted images (T1WI), from 180 adult participants scanned at Hammersmith Hospital using a 3-T system. We excluded participants with any of the following: missing demographic information, severe imaging artifacts that compromised vessel visualization, distinct structural abnormalities in the brain, or preprocessing failures (e.g., unsuccessful coregistration between paired volumes or failure of automatic surface reconstruction). Finally, 154 participants were included for analysis in this study (*SI Appendix*, Figure S5). Their ages ranged from 20 to 81 years (*SI Appendix*, Figure S6). The IXI Dataset describes the participants as normal and healthy, although precise definitions of these terms are not provided. Further details of the included participants are provided in *SI Appendix*, Table S7.

In this study, we selected the MCAs as vascular structures of interest because they supply the largest cerebral territory, follow a complex anatomical course, and are critically involved in various cerebrovascular pathologies (3, 4, 12, 47).

#### Image processing pipeline

Three-dimensional vessel reconstruction and geometric feature extraction from MRA were performed using custom pipelines implemented in Python (version 3.10), whereas automated brain segmentation and brain-surface reconstruction from T1WI were performed using Python and the Computational Anatomy Toolbox 12 (CAT12). The image-processing workflow is summarized schematically in *SI Appendix*, Figure S7. We extracted smooth, resampled centerlines from the reconstructed MCA surface meshes using the Vascular Modeling Toolkit (VMTK) (48, 49). Further details are provided in *SI Appendix*, Section S5.3.

#### Tortuosity quantification and statistical analysis

Classical length-based metrics, including centerline length *L*, Euclidean distance *L*_0_, and the excess-length ratio *ρ*, were quantified for M1 segments. For each MCA centerline interval bounded by branch points or distal endpoints, we fitted a quadratic baseline. We used the Bishop (parallel-transport) frame along this baseline to obtain the two transverse components of the centerline deviation. We then fitted a sinusoid-plus-trend model to these components to estimate the wavelength 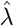 and combined amplitude *Â* (*SI Appendix*, Figure S2). Intervals meeting the periodic-fit qualification criteria, including at least 1.5 fitted cycles, and showing predominantly tangential periodic deviations (*R*_tan_ *>* 0.75, where *R*_tan_ is the tangential squared-displacement fraction relative to the brain surface) were selected for the primary tests of the model predictions. The accuracy of wavelength and amplitude estimation and the effects of estimation and qualification on wavelength–diameter scaling were evaluated using noise-free synthetic centerlines.

Associations were estimated using univariable and multivariable OLS regression with participant-level cluster-robust standard errors to account for multiple segments or intervals per participant. Associations of age with M1 morphological characteristics were evaluated. For periodic arterial waves, wavelength–diameter scaling was evaluated on the log–log scale, and associations of age with the wavelength-to-inner-diameter ratio and combined amplitude were evaluated using log-transformed outcomes. All multivariable models were adjusted for sex, ethnic group, the cube root of total intracranial volume, and side. M1 models additionally included M1 major branch type, and periodic-wave models additionally included anatomical MCA segment. The wavelength–diameter scaling model also included age, and the amplitude model additionally included log-transformed wavelength and log-transformed inner diameter. Exploratory spline and piecewise linear regression analyses examined age associations with the M1 excess-length ratio *ρ*. A two-sided *P* value *<* 0.05 was considered statistically significant. Details of these procedures and the statistical analyses are provided in *SI Appendix*, Sections S5.4, S5.5, and S5.6.

### 6.3 Use of AI tools

The authors used GPT-6 Astra (OpenAI) for language editing and Cursor (Anysphere) with automatic model selection to prepare the analysis code for public release.

## 7 Data, Materials, and Software Availability

All MRI data analyzed in this study were obtained from the publicly available IXI Dataset (https://brain-development.org/ixi-dataset/). The custom code and processed measurements used in this study are available on GitHub at https://github.com/tomo-akiyama/cerebral-artery-tortuosity-buckling.

## 8 Acknowledgments

## Author contributions

T.A. and T.M. designed research; T.A. and T.M. performed research; T.A. and T.M. contributed new analytic tools; T.A. and T.M. analyzed data; and T.A., T.M., and K.Y. wrote the paper.

## Abbreviations

1D: one-dimensional
3D: three-dimensional
ACA: anterior cerebral artery
CI: confidence interval
CSF: cerebrospinal fluid
ICA: internal carotid artery
IQR: interquartile range
M1: horizontal (sphenoidal) segment of the MCA
M2: insular segment of the MCA
M3: opercular segment of the MCA
M4: cortical segment of the MCA
MCA: middle cerebral artery
MRA: magnetic resonance angiography
MRI: magnetic resonance imaging
OLS: ordinary least squares
T1WI: T1-weighted images.

## Supplementary Methods

Unless otherwise specified, variables and parameters follow the definitions given in the main text. In the nomenclature (Table S1), superscripts (·)^sim^ and (·)^img^ denote simulation-derived quantities and quantities of the arteries imaged with magnetic resonance angiography (MRA), respectively, and are omitted when the context is unambiguous.

## S1. Model extension

### S1.1 Incorporating internal pressure loading and initial axial stretch

We extend the reference model to include internal pressure loading and initial axial stretch. Here, *p* denotes the transmural pressure, defined as the internal pressure minus the pressure of the surrounding cerebrospinal fluid (CSF), and is assumed to be spatially uniform and constant in time. The cross-sectional geometry and mechanical coefficients are held fixed, as in the reference model. Adding the transmural pressure contribution 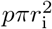 (1) to the axial force defined in Eq. 5 gives the effective axial force

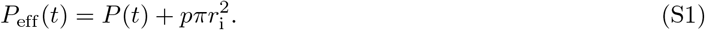

Arteries are normally axially prestretched *in vivo* (2). For an idealized straight initial configuration, we define the initial axial stretch ratio as

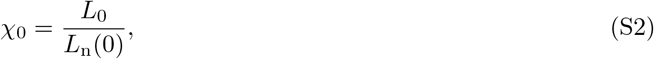

where *χ*_0_ *>* 1 represents this prestretch (3). For a constant natural-length elongation rate *g >* 0, we therefore obtain

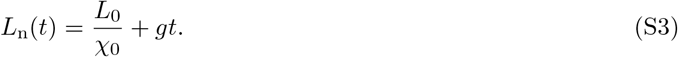

We retain the axial-force relation in Eq. 5, using this expression for *L*_n_(*t*). For this straight configuration, setting *t* = 0 yields

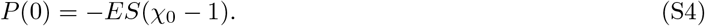

A negative value of *P* (*t*) indicates axial tension. Initial axial stretch is therefore incorporated through *L*_n_(0), without introducing an additional baseline-tension term.

Replacing *P* (*t*) in Eq. 2 with *P*_eff_ (*t*) gives the extended governing equation:

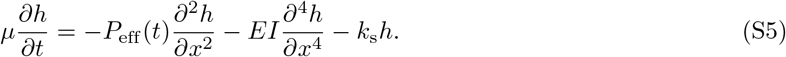

The reference model is recovered by setting *p* = 0 and *χ*_0_ = 1. The pressure contribution 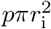 is constant in this extension. Bending reduces *P* (*t*) through the same geometric feedback as in the reference model.

For fixed *EI* and *k*_s_, the linear stability analysis yields the same critical wavelength *λ*_cr_ and critical axial force *P*_cr_ as in the reference model (Eq. 8). However, the time required to reach the instability threshold depends on transmural pressure and initial axial stretch. Provided that 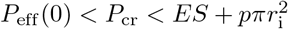, the corresponding critical time is

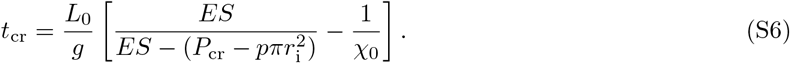

We set *χ*_0_ = 1.10 as a literature-informed reference value based on estimated *in vivo* axial stretch in human anterior cerebral arteries from donors aged 65 years or younger (4); this is not a direct measurement of initial axial stretch in young-adult middle cerebral arteries (MCAs). We assume that the systemic mean arterial pressure (MAP) serves as a surrogate for the mean internal pressure in the M1 segment. Considering the reference values of 90 mmHg for MAP and 15 mmHg for the CSF pressure, we set *p* = 0.01 N*/*mm^2^ (approximately 75 mmHg). Using the M1-informed parameters summarized in Table 2, the initial effective axial force is calculated as *P*_eff_ (0) ≈ −0.347 N. For this parameter set, the onset of instability is delayed to *t*_cr_ ≈ 31.2 yr, compared with approximately 10.9 yr in the reference model without internal pressure loading or initial axial stretch. The extension, nevertheless, preserves the underlying mechanism: progressive natural-length elongation can drive instability and increasing tortuosity, while transverse deformation relaxes effective axial compression through nonlinear geometric feedback.

## S2. Parameter set

### S2.1 Order-of-magnitude estimation of the surrounding foundation stiffness *k*_s_

We performed an order-of-magnitude analysis to estimate the foundation stiffness of the surrounding tissue *k*_s_. Cerebral arteries course within the subarachnoid space (SAS), a heterogeneous fluid–solid environment containing CSF and a network of collagenous arachnoid trabeculae that connect arachnoid and pial structures (Figure 2b,c) (5). We assumed that the surrounding environment resists transverse displacement of the vessel on the brain surface. We modeled this resistance as a local linear Winkler foundation (6), yielding the restoring force per unit vessel length −*k*_s_*h* in Eq. 2.

Following the proportionality assumption used in previous studies (1, 7), we parameterized *k*_s_ as

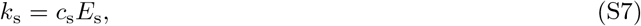

where *E*_s_ is the equivalent Young’s modulus of the homogenized surrounding matrix, and *c*_s_ is a dimensionless factor accounting for geometry, Poisson’s ratio, and load transfer. For the material parameters, we adopted the soft-solid representation of the SAS examined by Saboori and Sadegh (5). In this representation, the SAS was modeled as a homogeneous, isotropic, linear-elastic continuum with a Young’s modulus *E*_s_ = 0.001 N*/*mm^2^ and a Poisson’s ratio *ν*_s_ = 0.48. These values come from a head-impact model; their applicability to slow arterial deformation is assumed here.

To estimate *c*_s_, we considered a local simple-shear idealization of the homogenized SAS layer. We assumed that a relative transverse vessel displacement *h* parallel to the brain surface is accommodated by approximately uniform shear over a characteristic distance *d*_s_ normal to the brain surface. The shear strain in the surrounding tissue is then *h/d*_s_. For an isotropic linear-elastic material, the shear modulus *G*_s_ and shear stress *τ*_s_ are given by

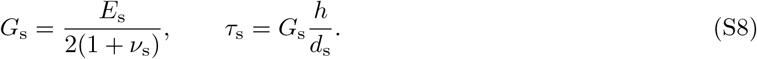

We introduced an effective width *b*_eff_ in the local tangent plane, transverse to the vessel axis. In this idealization, it represents the effective transverse extent of the surrounding SAS over which shear stress contributes to the restoring force. The restoring force per unit vessel length acts opposite to the displacement and is given by

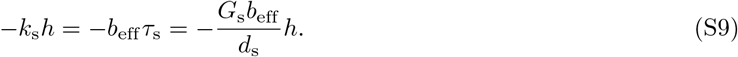

It follows that

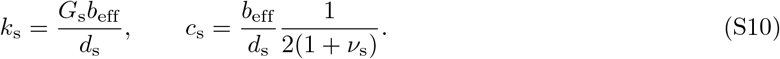

The representative outer vessel diameter (3.6 mm; Table 2) and the millimeter-scale SAS thickness assumed in a previous global head model (5) were taken as geometric reference scales. Within this local simple-shear idealization, we assumed that *b*_eff_ and *d*_s_ are millimeter-scale lengths of comparable magnitude, giving *b*_eff_ */d*_s_ = *O* (1). With *ν*_s_ = 0.48 and 1*/*[2(1 + *ν*_s_)] ≈ 0.34, this scaling gives *c*_s_ = *O* (1). We adopted *c*_s_ = 1 as an order-unity reference value. Together with *E*_s_ = 0.001 N*/*mm^2^, this gives *k*_s_ = 0.001 N*/*mm^2^ in the representative M1 parameter set. The weak dependence of the characteristic wavelength on *k*_s_ is addressed in the Discussion of the main text.

Cerebral perforating arteries arise from parent vessels and penetrate the brain tissue (8). These small arteries may therefore act as discrete anatomical tethers; however, the present model represents the surrounding support as a homogenized elastic foundation and does not explicitly resolve this potential localized restraint.

### S2.2 Selection of the effective natural-length elongation rate *g* **and damping coefficient** *µ*

In the model, the natural length was prescribed as *L*_n_(*t*) = *L*_0_ + *gt*, where *L*_0_ is the fixed projected span, and *g* = *d*(*L*_n_ − *L*_0_)*/dt* is the effective elongation rate (mm*/*yr). To relate the model to the MRA data, let 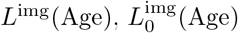 and 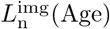 denote age-dependent mean values of the M1 centerline length, Euclidean distance, and unobservable natural length, respectively, with the other covariates held fixed. The age trend in excess natural length relative to the Euclidean distance can then be decomposed as

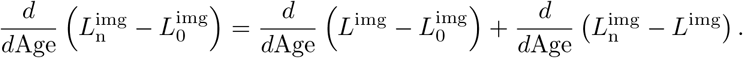

Because the residual mechanical mismatch 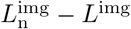 is not observable on MRA, the prescribed rate *g* cannot be uniquely inferred from the MRA data. In the multivariable M1 analysis (Table S3), the age coefficient was *β*_*L*_ = 0.063 mm*/*yr for centerline length and 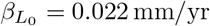 for Euclidean distance. We therefore used the difference between these cross-sectional age coefficients, 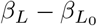, as an MRA-informed reference for selecting the effective elongation rate. Using the median M1 Euclidean distance in young adults (*<* 35 years), 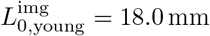, we normalized the difference between the age coefficients:

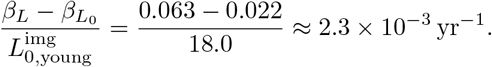

This normalized age association was used as an order-of-magnitude reference for selecting *g/L*_0_, where *L*_0_ = 100 mm is the projected span.

The effective damping coefficient *µ* controls the timescale of transverse deformation. In the linearized dynamics (Eq. S14), the modal growth rate at a given compressive force is inversely proportional to *µ*, whereas the critical axial force *P*_cr_ and the corresponding critical time *t*_cr_ remain independent of *µ* (Eqs. S16 and S21). Like *g, µ* cannot be directly determined from cross-sectional M1 geometry.

We selected *g* = 0.35 mm*/*yr (i.e., *g/L*_0_ = 3.5 × 10^*−*3^ yr^*−*1^) and *µ* = 0.005 N yr*/*mm^2^ as a reference combination to illustrate growth-driven buckling over a 60-year adult-equivalent simulation window (ages 20–80 years). A noticeable increase in deformation around *t* = 30 yr, corresponding to middle adulthood (approximately 50 years on the age-equivalent timescale), was used as a phenomenological timing target. During parameter selection, the third quartile of the pooled segment-level M1 distribution (*ρ*^img^ = 0.16; Table S2) was used as an empirical reference for the terminal deformation magnitude; the reference simulation reached *ρ*^sim^ ≈ 0.15 at *t* = 60 yr.

## S3. Numerical simulations

### S3.1 *A posteriori* assessment of the small-slope approximation

The governing equation (Eq. 6) uses the small-slope assumption |*∂h/∂x*|*≪*1 in both the transverse force balance (Eq. 2) and the axial compatibility relation (Eq. 5). We therefore assessed its *a posteriori* self-consistency by monitoring the maximum slope max_*x*_ |*∂h/∂x*| and the discrepancy induced by the quadratic arc-length approximation used to calculate *P* (*t*). We calculated the geometric excess-length ratio and its quadratic approximation from the same simulated profile:

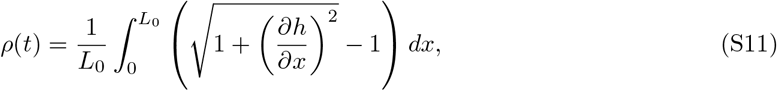

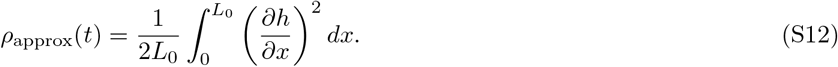

Here, *ρ* is calculated by applying the exact arc-length formula to the profile obtained from the present small-slope model. The quantity *ρ*_approx_ is the quadratic approximation used in Eq. 5: *P* (*t*) = *ES*[(*L*_n_(*t*) − *L*_0_) − *L*_0_*ρ*_approx_(*t*)]*/L*_n_(*t*). For *ρ >* 0, the relative discrepancy was defined as |*ρ*_approx_(*t*) − *ρ*(*t*)|*/ρ*(*t*).

Figure S8 shows the temporal evolution of max_*x*_ |*∂h/∂x*| and the two excess-length ratios, *ρ* and *ρ*_approx_. At the first time point at which *H*(*t*) exceeded the threshold *H*_rise_ = 0.5 mm (*t* = 30.4 yr), max_*x*_ |*∂h/∂x*| = 0.067 and the relative discrepancy was 0.085%. At the onset of the slow-growth stage (*t* = 33.0 yr), these values increased to 0.40 and 2.9%, respectively. This maximum slope corresponded to a maximum inclination angle of approximately 22^*°*^. During the late slow-growth stage, max_*x*_ |*∂h/∂x*| became of order unity, reaching 0.81 at the final time (*t* = 60 yr), when the relative discrepancy was 10.8%. Thus, the small-slope approximation remained internally consistent near buckling onset, whereas the small-slope condition was no longer satisfied during the late slow-growth stage. The late-stage behavior should therefore be interpreted as a qualitative extrapolation of the reduced model rather than as a quantitatively validated finite-amplitude prediction. Within this limitation, the simulation showed that post-buckling tortuosity continued to increase with prescribed elongation, a qualitative trend consistent with the age-associated increase in tortuosity observed in the cross-sectional MRA data.

We also monitored the effective axial strain, 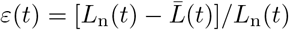. It reached a maximum of 9.5% at *t* = 30.8 yr and subsequently decreased to 3.8% at the final time (*t* = 60 yr). Linear elasticity in the axial direction was assumed over this strain range.

## S4. Mathematical analysis

### S4.1 Linear stability analysis

To characterize the early-stage pattern formation associated with vessel buckling, we performed a linear stability analysis. We consider interior buckling modes of a long arterial segment to isolate the intrinsic instability mechanism.

We first express Eq. 2 in Fourier modes, retaining the spatially uniform axial force *P* (*t*) determined by Eq. 5. For convenience, we represent a Fourier component with wavenumber *k* as *ĥ* (*k, t*)*e*^*ikx*^, where *ĥ* (*k, t*) is a complex modal amplitude. Substituting this Fourier mode into Eq. 2 yields

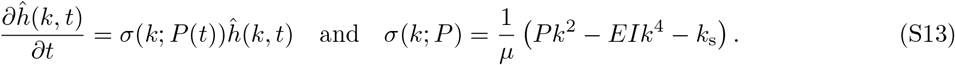

The physical displacement associated with this Fourier component is obtained as *h*_*k*_(*x, t*) = Re [*ĥ* (*k, t*)*e*^*ikx*^].

Around the straight state *h* = 0, the geometric nonlinear contribution to the axial force in Eq. 5 is of order (*∂h/∂x*)^2^ and can be neglected to linear order. The linearized modal evolution is therefore obtained by replacing *P* (*t*) with 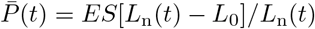:

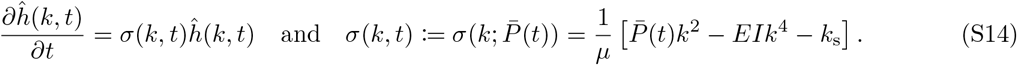

For continuous wavenumbers *k*, the onset of instability is determined by the conditions

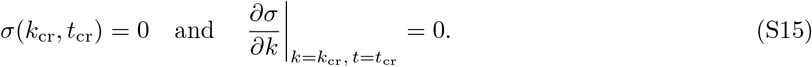

Solving these conditions gives the critical wavenumber, wavelength, and axial force:

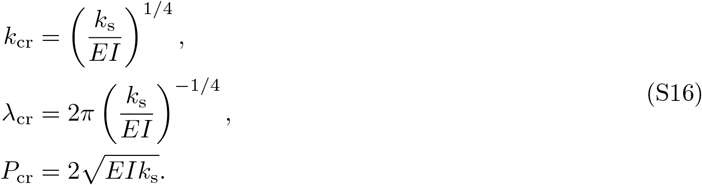

Equivalently, the condition *σ*(*k*_cr_, *t*_cr_) = 0 gives

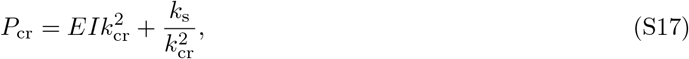

where 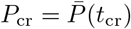.

For the finite periodic domain used in the numerical simulations, the admissible wavenumbers are

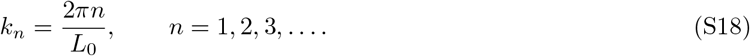

The continuous prediction *k*_cr_ corresponds to the theoretical mode number

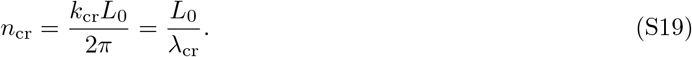

Because 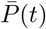 increases monotonically, the first admissible mode to become unstable is the one that minimizes the mode-specific critical axial force 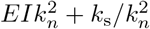.

At the linear instability threshold of the straight reference state, *L*(*t*_cr_) = *L*_0_. Therefore, the critical effective compressive axial strain is

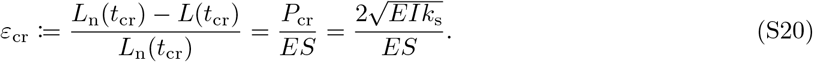

This expression shows that, for continuous wavenumbers, the critical strain does not depend explicitly on the vessel-segment length and is determined by the balance among the flexural rigidity *EI*, axial rigidity *ES*, and foundation stiffness *k*_s_.

Given a constant natural-length elongation rate *g >* 0, we have *L*_n_(*t*_cr_) = *L*_0_ + *gt*_cr_. The critical-strain relation (Eq. S20) therefore gives *ε*_cr_ = *gt*_cr_*/*(*L*_0_ + *gt*_cr_), which yields

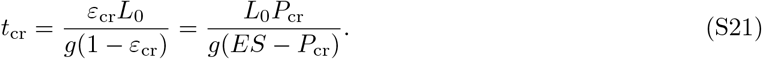

A finite critical time requires *P*_cr_ *< ES*.

We further examine the dependence of the critical wavelength *λ*_cr_ on vessel geometry. With an inner radius *r*_i_ and a wall-thickness ratio *η* = *t*_w_*/r*_i_, the outer radius can be written as *r*_o_ = (1 + *η*)*r*_i_, and the second moment of area is given by 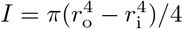. Substituting this into the expression for *λ*_cr_ (Eq. S16) gives

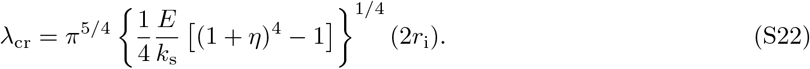

At fixed *η* and *k*_s_*/E*, this scaling relation shows that the critical wavelength is proportional to the inner diameter 2*r*_i_. The dependence on *k*_s_*/E* follows an inverse fourth-root law, while the dependence on *η* approaches fourth-root scaling in the thin-wall limit (*η ≪* 1).

Similarly, substituting the annular cross-sectional area 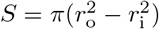 and the above expression for *I* into the expressions for the critical axial force (Eq. S16) and strain (Eq. S20) gives

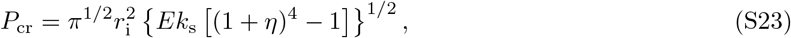

and

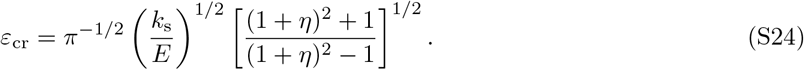

This latter formulation shows that the explicit dependence on the vessel radius cancels out in *ε*_cr_. The dependence on *k*_s_*/E* follows a square-root law, while the dependence on *η* approaches inverse square-root scaling in the thin-wall limit, in contrast to the corresponding fourth-root scaling of the critical wavelength (Eq. S22).

We next examine the amplitude growth near the critical time. For the critical mode *k* = *k*_cr_, Eq. S17 gives 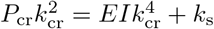. Substituting this relation into Eq. S14 yields

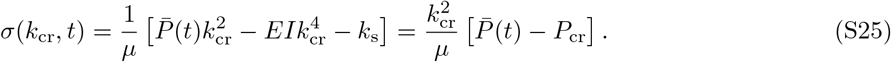

Near *t* = *t*_cr_, a first-order Taylor expansion of 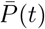 gives

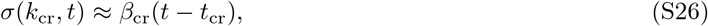

where

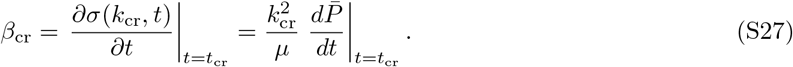

Here, *β*_cr_ is the temporal slope of the modal growth rate of the critical mode at *t* = *t*_cr_. For the linear natural-length elongation protocol used in the simulations, *L*_n_(*t*) = *L*_0_ + *gt*, the growth-induced axial force in the straight configuration (*h* = 0) is 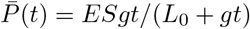. Substituting this expression into Eq. S27 gives

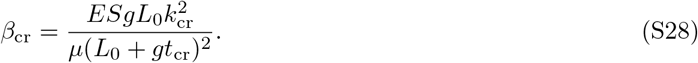

Let 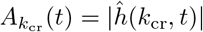 (*t*) = |*ĥ* (*k*_cr_, *t*)| denote the physical modal amplitude associated with the critical Fourier mode.

Since the growth rate *σ*(*k*_cr_, *t*) is real, this amplitude satisfies

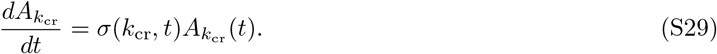

For *t* ≥ *t*_cr_, sufficiently close to *t*_cr_, substituting the linearly increasing growth rate (Eq. S26) into this amplitude equation gives

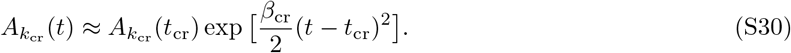

Thus, near the critical time, the exponent increases approximately quadratically in time. When *β*_cr_(*t* − *t*_cr_)^2^*/*2 *≪* 1, the amplitude remains close to its small residual value at *t* = *t*_cr_ and is therefore nearly unobservable. Once this exponent becomes of order unity, the amplitude increases rapidly, eventually leading to visible deformation.

To quantify the delay between the critical threshold and the emergence of observable deformation, we define the characteristic rise time *t*_rise_ as the time at which the modal amplitude reaches *R* times its value at *t* = *t*_cr_:

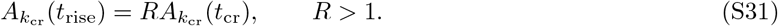

Using Eq. S30, we obtain

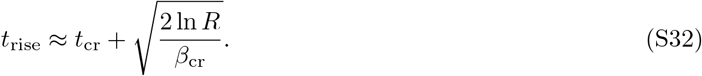

For comparison with the numerical simulations (main text, Section 3.1), we used the global deformation measure 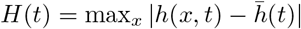. In the simulation time series, *H*(*t*) reached its minimum at 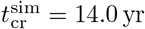, with 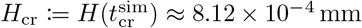. We also defined an operational rise time 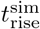 at which *H*(*t*) first exceeded the operational visibility threshold *H*_rise_ = 0.5 mm. Because *H*_cr_ represents the global deformation at 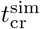 rather than the critical-mode amplitude at the analytical critical time *t*_cr_, we used *H*_rise_*/H*_cr_ ≈ 6.15 × 10^2^ as an effective value of *R* for an approximate comparison. We evaluated Eq. S32 using this value of *R* and the analytical values of *t*_cr_ and *β*_cr_. This yielded *t*_rise_≈ 30.0 yr, in reasonable agreement with the simulation-defined rise time 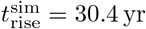.

### S4.2 Weakly nonlinear analysis

To further characterize the finite-amplitude behavior near the onset of instability, we performed a weakly nonlinear analysis of Eq. 6. Following the notation used in the linear stability analysis, the real-valued physical displacement associated with a Fourier mode of wavenumber *k* is written as

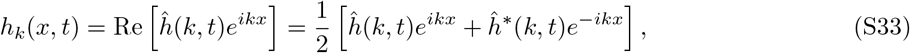

where *ĥ* (*k, t*) *∈*ℂ is the complex Fourier amplitude of the mode. With this normalization, *A*_*k*_(*t*) = |*ĥ* (*k, t*) | is the physical peak amplitude of the mode.

Near the onset of instability, we approximate the total displacement by the contribution of the critical mode with wavenumber *k*_cr_:

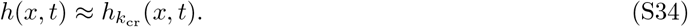

For notational simplicity, we write *h* for 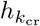 (*x, t*) and *ĥ* for *ĥ* (*k*_cr_, *t*) in the following calculation. From Eq. S33 with *k* = *k*_cr_, we obtain

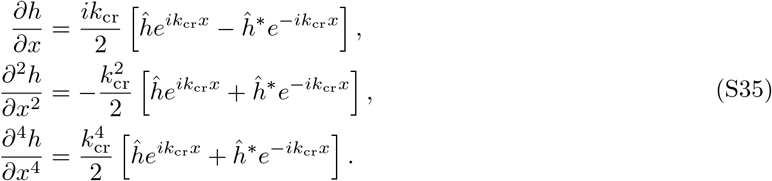

We first evaluate the nonlocal stretching integral. Using Eq. S35,

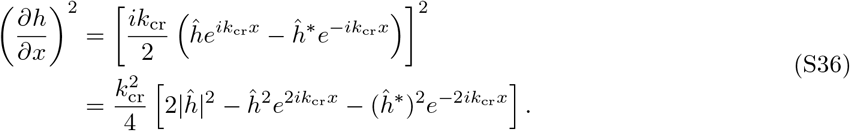

Because the analysis concerns interior buckling modes in a long arterial segment (*k*_cr_*L*_0_ *≫* 1) rather than end effects, the oscillatory terms proportional to 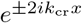 do not contribute to the leading-order evaluation of the integral over the reference interval. Therefore,

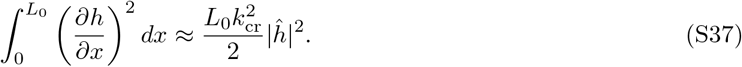

Thus, the nonlocal stretching measure is quadratic in the critical-mode amplitude. Substituting Eqs. S35 and S37 into the nonlinear geometric term gives

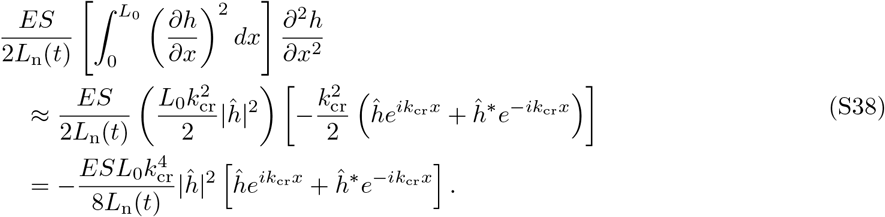

The linear terms on the right-hand side of Eq. 6 give

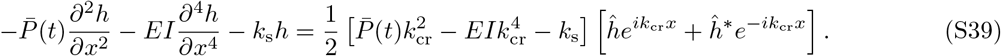

The left-hand side of Eq. 6 is

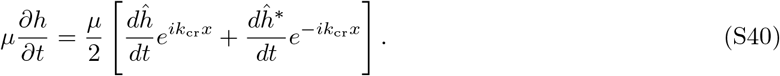

Collecting the coefficients of the 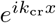 mode, we obtain

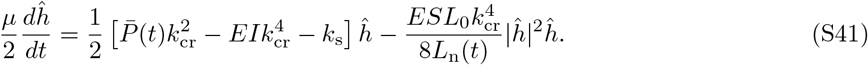

Using the linear growth rate of the critical mode (Eq. S25), this equation can be written as

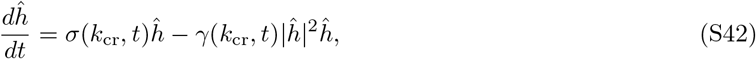

where

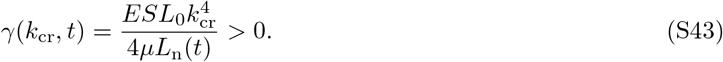

The nonlinear term −*γ*(*k*_cr_, *t*) |*ĥ* | ^2^*ĥ* is cubic in the critical-mode amplitude and opposes its growth.

We next convert the complex amplitude equation into the real amplitude equation. Let 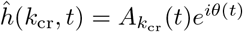, where 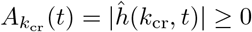 is the physical peak amplitude and *θ*(*t*) denotes the phase of the critical mode. For brevity, we omit the explicit *t*-dependence of 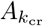 and *θ*. Substituting the polar form into the complex amplitude equation (Eq. S42) gives

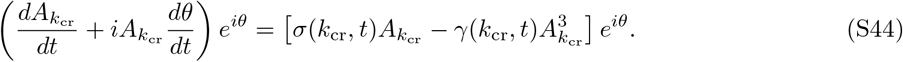

Canceling the common factor 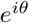 and equating real and imaginary parts yields the real amplitude equation

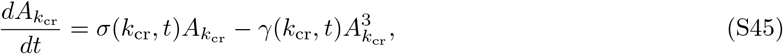

and the phase relation 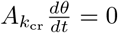. Thus, away from the trivial state 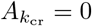, the phase remains constant. The positivity of *γ*(*k*_cr_, *t*) ensures that the cubic term provides a saturation mechanism for growth driven by the linear instability *σ*(*k*_cr_, *t*) *>* 0. With *σ* and *γ* treated as time-independent parameters, the corresponding equation for a signed amplitude has the normal form of a supercritical pitchfork bifurcation (9).

With *σ* and *γ* held fixed, setting 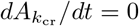 in Eq. S45 gives the nontrivial equilibrium branch:

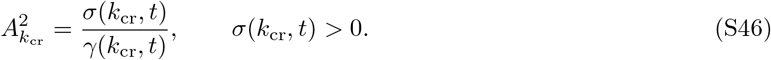

This branch provides a quasi-static reference for the growing system. Using Eqs. S25 and S43, we obtain

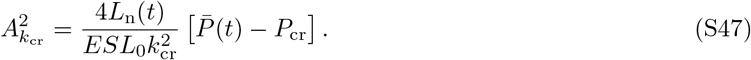

Therefore, along this equilibrium branch and sufficiently close to the critical axial force,

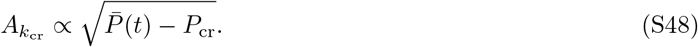

Next, we relate the modal amplitude to the geometrically measurable excess-length ratio *ρ*. Using the small-slope arc-length approximation in Eq. 3, we write

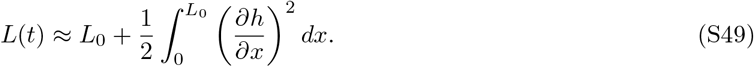

Substituting this approximation and Eq. S37 into the definition of *ρ* (Eq. S60) gives

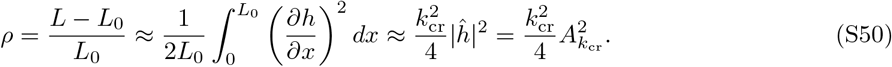

Along this equilibrium branch, using the amplitude relation in Eq. S47, we obtain

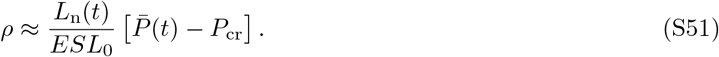

Near the onset, the prefactor can be approximated by its value at *t* = *t*_cr_, namely *L*_n_(*t*_cr_)*/*(*ESL*_0_). Thus, the equilibrium excess-length ratio obeys

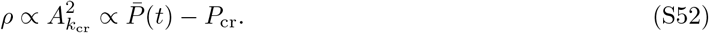

We next interpret the peak of the actual axial force *P* (*t*). Using Eq. S37 and 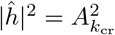, the axial-force relation in Eq. 5 can be approximated as

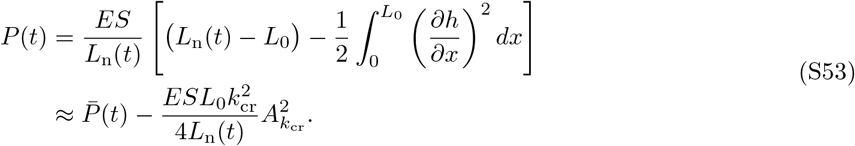

This expression shows that buckling accommodates part of the growth-induced length mismatch through additional arc length, reducing *P* (*t*) relative to the instantaneous straight-state force 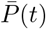. At the force peak, *dP/dt* = 0, so the time derivative of the geometric correction balances 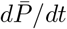.

### S4.3 Post-buckling quasi-static equilibrium and modal dynamics under continued growth

Following the force peak and subsequent relaxation phase, we investigate the late-time behavior of the axial force. We first determine the equilibrium at each fixed value of the natural length *L*_n_. Setting *∂h/∂t* = 0 in the governing equation (Eq. 2) yields the mechanical force balance:

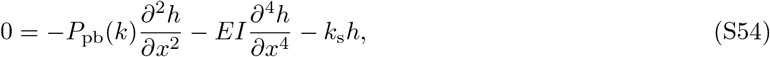

where *P*_pb_(*k*) denotes the equilibrium axial force associated with wavenumber *k*. For a fixed *k*, we consider a nonzero single-mode equilibrium using the representation in Eq. S33, with the time dependence omitted for the equilibrium calculation. Substituting this representation into Eq. S54 and collecting the coefficients of *e*^*ikx*^ gives

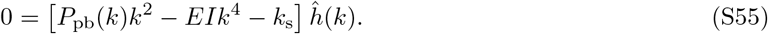

Since *ĥ* (*k*) ≠ 0, the equilibrium axial force is

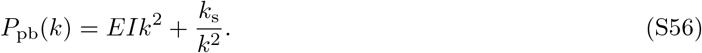

A nonzero single-mode equilibrium requires 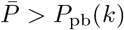, with its amplitude determined by axial compatibility. For fixed *k, EI*, and *k*_s_, *P*_pb_(*k*) is independent of the amplitude and natural length. It equals the critical axial force of the corresponding Fourier mode. For the continuous critical mode, Eq. S17 gives

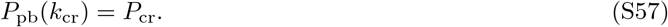

To relate this equilibrium force *P*_pb_(*k*) to the evolving state, we restore the time-derivative term. For a nonzero mode with the same fixed wavenumber *k*, Eq. S13 gives

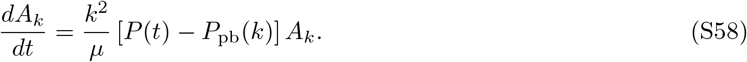

For fixed *k* and *µ*, the relative amplitude growth rate is proportional to the force difference *P* (*t*) − *P*_pb_(*k*). Thus, even a small positive force difference allows continued post-buckling amplification while the actual force remains close to *P*_pb_(*k*).

## S5. Geometric analysis of MRA data

### S5.1 Magnetic resonance imaging acquisition parameters

Magnetic resonance imaging (MRI) was performed using a 3-T scanner (Intera, Philips Medical Systems, Best, The Netherlands). Acquisition parameters are provided in the IXI Dataset documentation (IXI Dataset MRI parameters). Key parameters used in the study are summarized below. Magnetic resonance angiography (MRA) images were acquired using a time-of-flight technique with the following parameters: an echo time (TE) of 5.75 ms, a repetition time (TR) of 16.7 ms, a flip angle of 16^*°*^, an acquisition matrix of 288 × 286, and a reconstructed voxel size of 0.47 × 0.47 × 0.8 mm^3^. The T1-weighted images (T1WI) were acquired with a TE of 4.6 ms, a TR of 9.6 ms, a flip angle of 8^*°*^, an acquisition matrix of 208 × 208, and a reconstructed voxel size of 0.94 × 0.94 × 1.2 mm^3^.

### S5.2 Software information

Custom code for image processing, data extraction, and analysis was implemented in Python (version 3.10). The Python code has been made publicly available in our GitHub repository.

The Python packages used for image processing and geometric analysis included ANTsPy (10), brainextractor (a Python reimplementation of the Brain Extraction Tool), scikit-image (11), the Visualization Toolkit (VTK) (12), PyMeshLab (13), the Vascular Modeling Toolkit (VMTK) (14, 15), and PyVista (16). The Computational Anatomy Toolbox 12 (CAT12, version 12.9) (17), an extension of Statistical Parametric Mapping (SPM12) software, was executed in MATLAB (version 23.2, MathWorks) for brain tissue segmentation, brain surface reconstruction, and morphometric measurements. 3D Slicer (version 5.6) was used for visual inspection and manual procedures (18).

Matplotlib (19) and seaborn (20) were used for data visualization, pandas (21) for data management, and SciPy (22) and statsmodels (23) for statistical analysis.

### S5.3 Image processing

#### Preprocessing, segmentation, and mesh generation

The image processing workflow is summarized in Figure S7. First, both MRA and T1WI volumes were resampled to isotropic voxels of 0.5 mm using B-spline interpolation. For the resampled T1WI volumes, the following steps were automatically performed using the CAT12 standard pipeline: denoising using a spatially adaptive filter, bias-field correction, skull-stripping, brain tissue segmentation, and brain surface reconstruction. CAT12 preprocessing was performed at the input image resolution, with the bias correction strength and preprocessing accuracy parameters both set to 0.75. Surface reconstruction used the extra fine mesh setting (<monospace>vdist</monospace> = 0.5). The reconstructed pial surfaces were used as the brain-surface meshes and were then isotropically remeshed for 10 iterations with a target edge length of 0.33 mm. For MRA volumes, spatially adaptive non-local means denoising with a Gaussian noise model was used, with a patch radius of 1 voxel and a search radius of 3 voxels (24). Low-frequency intensity non-uniformity was then corrected using the N4 bias field correction algorithm (25). Skull-stripping was performed with a fractional intensity threshold of 0.45 (26). The preprocessed MRA volumes were coregistered to the CAT12 intensity-corrected T1WI in native space using affine registration (TRSAA), followed by nonlinear symmetric normalization (SyNOnly) (27). Mattes mutual information was used as the similarity metric in both stages. Spatial alignment was visually inspected in all participants.

We extracted the arterial tree from the large proximal cerebral arteries to their small distal cortical branches while preserving the continuity of the vascular structure. Vessel segmentation was performed on the coregistered, skull-stripped MRA volumes using hysteresis thresholding (28). Major cerebral arteries generally exhibit high signal intensity in time-of-flight MRA owing to the inflow effect (29). The high threshold was set to the 99.9th percentile of the positive voxel intensities in these volumes to extract initial vessel candidates corresponding to major arteries. The low threshold was set to the 98.5th percentile of the same intensity distribution to include distal cortical branches with lower signal intensity. Using three-dimensional (3D) connected component analysis, regions above the low threshold were retained if they contained voxels above the high threshold. Enclosed holes were then filled, and connected components containing fewer than 5,000 voxels (625 mm^3^) were removed.

Vessel segmentations were visually inspected. When necessary, the lumen contours were manually edited to remove misclassified voxels and artificial bridges between adjacent arteries or between arteries and nearby high-signal structures, such as veins. Perforating arteries branching off from the major arteries were also removed. Vascular branches independently detected at the high hysteresis threshold sometimes appeared disconnected from the main arterial tree in the segmentation because of locally reduced signal intensity. In these cases, anatomical continuity was restored manually only when the connection was clearly identifiable on the original MRA images. Manual voxel removal and correction may have introduced small, artifactual bends in the reconstructed vessels. The centerline smoothing described below was applied to reduce the influence of such local irregularities on subsequent geometric measurements. Finally, the MCAs were isolated from the arterial tree for analysis. Anatomical variants with separate origins, including accessory or duplicated MCAs (30), were also included in the analysis (accessory MCA: 3 hemispheres; duplicated MCA: 2 hemispheres).

Vessel surface meshes were generated from segmented vessels using a discrete implementation of the marching cubes algorithm (31) in VTK. Small, isolated components (enclosed volume ≤ 62.5 mm^3^) generated during the marching cubes procedure were removed. The meshes were then smoothed using Taubin’s algorithm with <monospace>lambda_</monospace> = 0.6307 and <monospace>mu</monospace> = − 0.6732 for 10 iterations (32), followed by 2 iterations of Butterfly subdivision applied to all edges to increase mesh density (33). Finally, spatially aligned meshes of the MCAs and the brain surface were obtained.

All visual inspections and manual procedures were performed by an expert with over 10 years of clinical experience in cerebrovascular anatomy and imaging (TA, a board-certified neurosurgeon and board-certified neuroendovascular specialist in Japan).

#### Vessel centerline extraction

To characterize the geometries of the MCAs, we extracted vessel centerlines using the Voronoi diagram of the triangulated surface mesh (14, 15). Local inner (lumen) radii were obtained from the maximum-inscribed-sphere radius values provided by VMTK along the centerlines. Insufficient surface mesh resolution can reduce centerline accuracy, particularly in small distal vessels (34). Across 154 participants, the median of the participant-specific median edge lengths was approximately 0.11 mm. The final vascular surface meshes had a median face density of 192.8 (interquartile range [IQR], 192.2–193.3) triangular faces per mm^2^.

To reduce local irregularities associated with discretization and manual editing, the centerlines were smoothed using a Laplacian filter with a smoothing factor of 1.2 and 100 iterations (15, 35). The smoothed centerlines were then resampled at 0.1 mm intervals along their arc length. These parameter values were chosen with reference to a previous study of internal carotid artery geometry (35). The median total centerline length per hemisphere was 1,051.8 (IQR, 916.6–1,185.3) mm.

### S5.4 Anatomical definitions and M1 geometric measurements

For M1 elongation and tortuosity analysis, measurements were made along anatomically defined M1 paths (main text, Section 3.3).

#### Anatomical definitions and M1 measurements

The MCA was divided into M1 (horizontal or sphenoidal), M2 (insular), M3 (opercular), and M4 (cortical) segments according to anatomical location (Figure 2) (36, 37).

The M1 segment originates at the bifurcation of the internal carotid artery (ICA) into the MCA and the anterior cerebral artery and extends laterally within the deep Sylvian fissure, posterior and roughly parallel to the sphenoid ridge. This segment is typically relatively straight in younger individuals, whereas it tends to adopt a curved configuration with advancing age (38–40). This relatively straight course makes excess centerline length relative to the end-to-end distance easier to interpret than in arteries that curve substantially to follow the surrounding brain anatomy. We therefore selected M1 to assess age-related differences in centerline length, Euclidean distance, and excess-length ratio. The precise landmark that demarcates the transition from the M1 to the M2 segment remains debated (39): classic microsurgical anatomy studies use an anatomical landmark, placing the M1–M2 boundary at the MCA genu at the level of the limen insulae (36, 37), whereas recent neuroendovascular literature often defines M1 as extending to the main MCA bifurcation (30, 41). Moreover, the MCA exhibits substantial anatomical variability (39), and measured M1 length depends on both individual anatomy and the definition of the distal endpoint (36, 39, 42). To establish a consistent measurement boundary, we used a limen insulae-based reference surface to define the distal M1 endpoint, guided by microneurosurgical descriptions (36, 37). Reference points corresponding to the limen insulae were selected in the Montreal Neurological Institute 152 (MNI152) template (43) at (*x, y, z*) = (35, 8, − 18) mm on the right and ( − 35, 8, −18) mm on the left (Figure S9a). Sagittal planes passing through these points were defined in MNI152 space. Each participant’s T1WI was registered to the template, and the inverse transformation was applied to map the entire planes into the participant’s T1WI space. Because the MRA volumes had already been coregistered to T1WI, the mapped reference surfaces were used directly to define the distal M1 endpoints at their intersections with the M1 centerlines. The proximal endpoint was defined at the ICA terminus, where the ICA divides into the anterior cerebral artery and the MCA.

Under this definition, three M1 major branch types were observed, and measurement paths were selected accordingly (Figure S9b). When the main division occurred at or beyond the reference surface, one M1 path was measured (single-trunk type). When the main division occurred before the reference surface and the two trunks were judged comparable in caliber and territorial extent, both paths were measured (codominant-trunk type). When the main division occurred before the reference surface and one trunk was clearly dominant in caliber and territorial extent, only the path following that trunk was measured (dominant-trunk type). For trifurcations (or higher-order divisions), the same dominance-based selection rule was applied. Each measured path extended from the ICA terminus to its corresponding intersection with the reference surface. M1 centerline length, Euclidean distance, and length-based tortuosity metrics were calculated for these anatomical paths without endpoint trimming.

#### Definition of classical tortuosity metrics

The Euclidean distance *L*_0_ is defined as the straight-line distance between the vessel endpoints, and *L* denotes the total centerline length. Tortuosity is summarized by the conventional tortuosity index *T*, defined as:

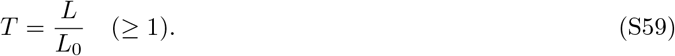

This metric quantifies the departure from straightness and is widely used because it is simple and interpretable (2, 44). The corresponding excess-length ratio *ρ* is defined as (15):

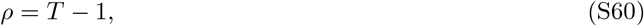

where *ρ* is a dimensionless measure of excess centerline length relative to the current end-to-end Euclidean distance.

We used *ρ* only for the anatomically defined M1 paths. The relatively straight M1 course in younger individuals makes excess length relative to *L*_0_ more readily interpretable than in arteries with pronounced anatomical curvature. Here, *ρ* measures the fractional excess of centerline length over the distance between the vessel endpoints. In the model, this ratio increases as post-buckling deformation progresses. However, these length-based metrics do not characterize periodicity, distinguish planar from nonplanar centerline configurations, or describe the orientation of deviations relative to the brain surface. We therefore performed a complementary 3D moving frame analysis.

### S5.5 Quantification of periodic transverse centerline deviations using a moving frame

Periodic wave analysis used MCA centerline intervals (main text, Section 3.3). The interval-selection and directional-classification workflow is summarized in Figure S3.

#### Definition and anatomical classification of MCA centerline intervals

For analysis of local periodic waves, MCA centerline intervals were defined as continuous branch-to-branch or branch-to-endpoint paths identified in the reconstructed vascular network, with no internal detected branch points. These intervals were defined by the branching structure independently of anatomical compartment labels.

Each interval included in the primary tests of the model predictions was classified as M1, M2–3, or M4 according to the anatomical compartment containing its arc-length midpoint (Figure 2). M2 and M3 segments were analyzed together because both are located within the operculoinsular compartment of the Sylvian fissure (36, 37). M4 corresponded to cortical branches beyond the operculoinsular compartment. Anatomical assignments were visually inspected.

#### Definition of transverse centerline deviations using the Bishop frame

For each interval, the observed 3D centerline was represented as a discrete sequence of points resampled at 0.1-mm arc-length intervals. The sampled positions were denoted by **P**(*s*) *∈* ℝ^3^, where *s ∈* [0, *L*] is the observed-centerline arc-length parameter and *L* is the full observed centerline length. To define a smooth geometric reference, a baseline centerline **P**_0_(*s*) was constructed by least-squares fitting a second-order polynomial to each Cartesian component of **P** as a function of *s* over the full interval (Figure S2a). This baseline was intended to approximate the broad background curvature associated with large-scale brain-surface topography and vessel routing. It served as a geometric reference rather than a mechanically defined pre-buckling configuration. The geometric residual was defined at matching values of the observed arc-length parameter:

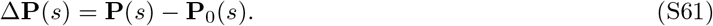

The Bishop (parallel-transport) frame was constructed along **P**_0_ using the unit tangent vector

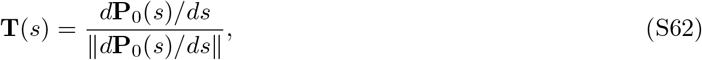

and two orthonormal transverse vectors, **N**_1_(*s*) and **N**_2_(*s*), spanning the plane normal to **T** (45). Here, *s* is the observed centerline arc-length parameter, not the baseline arc-length parameter. In the continuous representation, the transverse vectors satisfy

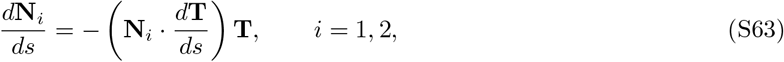

which excludes rotation about the local tangent. For a baseline with *∥d***P**_0_(*s*)*/ds∥ >* 0, the Bishop construction remains well defined at zero curvature and avoids the normal-vector flips that can occur in the Frenet–Serret frame at inflection points. In the discrete implementation, **N**_1_ was initialized as a unit vector orthogonal to the initial tangent. The frame was propagated by minimal rotations aligning successive baseline tangents, with **N**_2_ = **T**× **N**_1_. Brain-surface normals were not used to construct the frame.

The signed transverse components were defined by projection onto the transverse basis (Figure S2b):

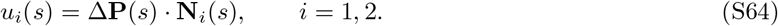

The corresponding transverse deviation vector was given by

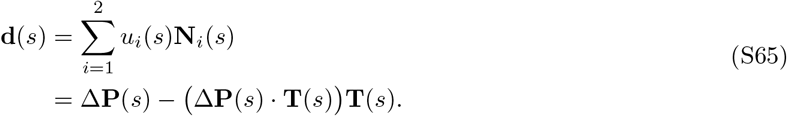

This transverse deviation was used as a geometric measure of vessel waves for comparison with the transverse displacement represented by *h*(*x*) in the one-dimensional model, rather than as a direct measurement of that displacement. After the transverse components were calculated over the full observed interval *s ∈* [0, *L*], the first and last 10 samples were excluded from sinusoid-plus-trend fitting to reduce endpoint effects. This trimming corresponded to approximately 1 mm of arc length at each end of the observed centerline. The same sample indices were retained on the baseline.

For fitting and reconstruction, the arc-length coordinate along the baseline was denoted by *s*_0_ and shifted to zero at the baseline point corresponding to the first retained sample. This coordinate was evaluated at all corresponding baseline samples, including the excluded end samples. The retained interval therefore spanned 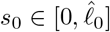, where the estimated baseline length 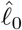 is the arc length along the baseline between the first and last retained samples. In the following, the frame and transverse quantities are indexed by *s*_0_ using the same pointwise correspondence, and *s* denotes the corresponding observed-centerline arc-length coordinate.

#### Single-mode sinusoid-plus-trend fitting

To account for the effect of quadratic baseline subtraction on periodic deviations, candidate transverse sinusoidal modes were subjected to the same quadratic detrending and projection operations used to obtain the observed transverse components. For each candidate wavenumber *k*, four transverse sinusoidal basis functions were constructed using the baseline Bishop frame and assembled as the columns of the matrix

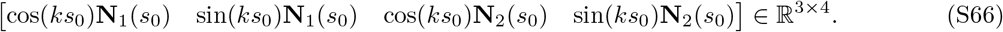

Here, *k* is defined with respect to the baseline arc-length coordinate *s*_0_.

For each column of this matrix, a second-order polynomial was fitted separately to each Cartesian component as a function of *s*, using all samples before end trimming. The fitted polynomials were subtracted from the corresponding Cartesian components of each basis function, and the resulting residual vectors were projected onto **N**_1_(*s*_0_) and **N**_2_(*s*_0_). Only the retained samples were then used for sinusoid-plus-trend fitting. Because the frame varies along the baseline, Cartesian quadratic subtraction need not preserve alignment with the original transverse basis direction. Consequently, each residual vector can contribute to the transverse components along both **N**_1_(*s*_0_) and **N**_2_(*s*_0_).

For notational simplicity, let *Q* denote the operator that returns the componentwise least-squares quadratic approximation of a sampled vector field over the full range of *s*. Its output is indexed by the corresponding baseline coordinate *s*_0_. The scalar transverse components of the candidate sinusoidal mode before quadratic subtraction were written as

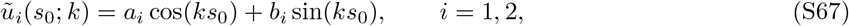

where *k* is the wavenumber shared by both transverse components, and *a*_*i*_ and *b*_*i*_ are the cosine and sine coefficients associated with **N**_*i*_, respectively. The corresponding candidate sinusoidal mode was then given by

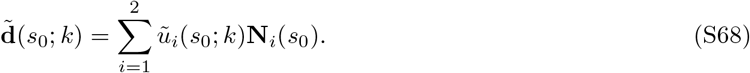

The baseline Bishop frame was held fixed throughout the fitting. Quadratic subtraction and projection are linear operations. Each transverse deviation component on the retained interval was therefore modeled using the processed sinusoidal mode plus a linear trend:

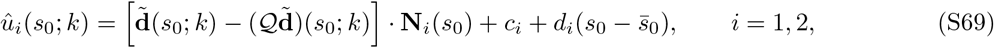

where *c*_*i*_ and *d*_*i*_ are the intercept and slope of the linear trend, respectively, and 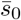 is the mean of *s*_0_ over this interval. The quadratic baseline defined the 3D geometric reference, whereas the additional linear trend accounted for offsets and gradual drift in the projected components.

For each interval, the median inner radius *r*_i_ was calculated over the samples used for sinusoid-plus-trend fitting. To focus on centerline waves while filtering out sub-vessel-scale fluctuations, the lower wavelength bound was set to 4*r*_i_ (twice the inner diameter). To ensure that each candidate mode spanned at least half a spatial cycle over 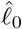, the upper wavelength bound was set to 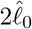. The corresponding wavenumber bounds were 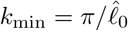 and 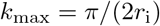. The candidate wavenumber grid started at 0.5 cycles over 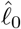, with increments of 1*/*16 cycle, and included the upper wavenumber bound. For each candidate *k*, the sinusoidal and trend coefficients were estimated jointly by ordinary least squares (OLS) using both transverse components. The shared wavenumber 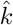 was selected by maximizing the coefficient of determination (*R*^2^) for the two-component sinusoid-plus-trend fit over the numerically valid grid candidates.

#### Reconstruction and geometric characterization of the periodic mode

Using the coefficients estimated at 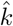, the fitted sinusoidal components before quadratic subtraction were given by

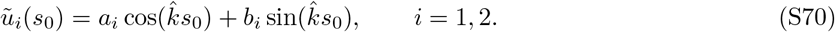

The periodic component of the transverse deviation, hereafter the periodic deviation, was then reconstructed in 3D using the same baseline Bishop frame (Figure S2c):

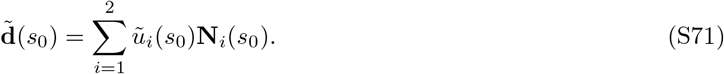

The estimated combined amplitude and wavelength were calculated as

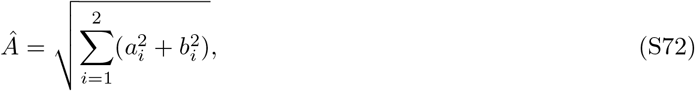

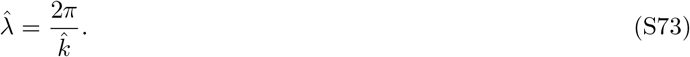

This root-sum-square measure is invariant to a fixed rotation of the Bishop normal-plane basis and to a shift of the phase origin (Figure S2d). It reduces to the conventional scalar sinusoidal amplitude when the periodic deviation is confined to a single transverse direction. Thus, *Â* was used as a rotation-invariant geometric proxy for the magnitude of the transverse periodic mode predicted by the one-dimensional model.

In Bishop transverse coordinates, the fitted sinusoidal pair (*ũ*_1_, *ũ*_2_) parameterizes an elliptical orbit (Figure S2d). Let *M* denote the fitted coefficient matrix:

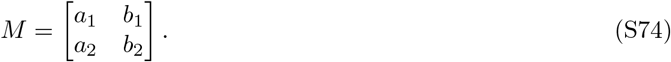

The singular values of *M*, denoted by *Â* _major_ ≥ *Â* _minor_ ≥ 0, represent the major and minor semiaxes, respectively, with 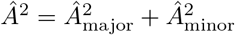. Ellipticity was defined as *Â* _minor_*/ Â* _major_ for *Â* _major_ *>* 0, ranging from 0 for a wave along a single transverse direction to 1 for a circular trajectory in transverse coordinates.

This measure was used to describe the mode geometry.

The orientation of the periodic deviation relative to the brain surface was evaluated separately, as described below.

#### Selection of periodic-fit–qualified intervals

To assess the contribution of the sinusoidal mode beyond the linear trend, we calculated the partial *R*^2^:

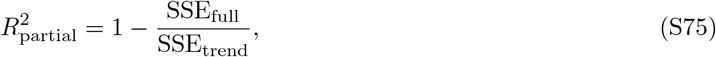

where SSE_full_ and SSE_trend_ are the residual sums of squares for the processed sinusoid-plus-trend model at 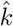 and a separately fitted linear-trend-only model, respectively. Both sums were calculated across the two transverse components using the same fitting samples.

The fitted cycle count was defined as

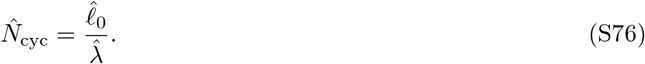

Intervals with numerically valid fits were classified as periodic-fit–qualified when all of the following conditions were met:

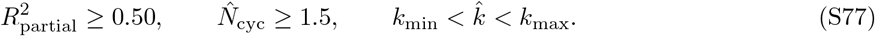

Selection of either search boundary indicated that the wavelength estimate could be limited by the prescribed search range; such fits were therefore excluded. Qualified intervals were retained for subsequent assessment of the orientation of the periodic deviation relative to the brain surface.

#### Brain-surface normal estimation and directional classification

Brain-surface directional classification was used to select intervals suitable for comparison with the tangential waves assumed by the model, with the aim of excluding, as far as possible, intervals whose estimated wavelengths might reflect surface-normal waves following brain-surface topography. This selection does not imply that tangential waves were absent from the other intervals. To characterize the orientation of the periodic deviation relative to the brain surface, local brain-surface geometry was evaluated along the retained vessel centerline of each periodic-fit–qualified interval. Anchor points were placed every 10 centerline points, corresponding to approximately 1 mm, with the final point also included. At each anchor, vertex normals were collected from the brain-surface mesh within a spherical neighborhood centered at the anchor. The neighborhood radius was four times the median inner radius within the interval, bounded to the range 2.5–5.0 mm.

For the *m*_*a*_ neighboring unit vertex normals **n**_*aj*_ at anchor *a*, the sign-invariant orientation tensor was constructed as

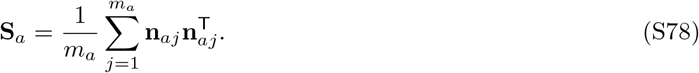

The effective surface-normal axis was represented by a unit eigenvector **ñ**_*a*_ associated with the largest eigenvalue of **S**_*a*_, and the corresponding eigenvalue was defined as the surface-normal coherence *C*_*a*_ (Figure S10a). The coherence *C*_*a*_ ranges from 1*/*3 for an ideal isotropic distribution of normal axes to 1 when all normal axes are aligned, irrespective of vector sign. The vector **ñ**_*a*_ was accepted at anchors with at least 30 neighboring normals and *C*_*a*_ ≥ 0.75 (Figure S10b,c). When two consecutive anchors both satisfied these criteria, their vectors **ñ**_*a*_ were aligned in sign, linearly interpolated along centerline arc length, and normalized to unit length. Gaps containing unaccepted anchors were bridged using only the vectors **ñ**_*a*_ at the accepted anchors on both sides when these anchors were at most 5.0 mm apart along the centerline and the unsigned angle between their axes was at most 45^*°*^. No extrapolation was performed at interval ends.

Directional classification required that points with evaluable normals comprised at least 50% of the retained centerline points and accounted for at least 50% of the summed squared magnitude of the periodic deviation over the interval. The tangential squared-displacement fraction was defined as

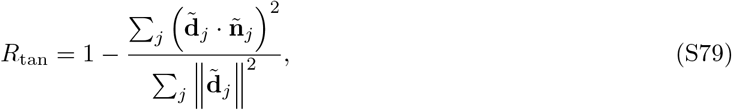

where 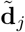 is the periodic deviation (Eq. S71) evaluated at centerline point *j*, and **ñ**_*j*_ is the corresponding unit effective surface-normal vector obtained from the accepted vectors **ñ**_*a*_. Both sums were calculated only over points with evaluable normals, including bridged positions. High values of *R*_tan_ indicate a predominantly tangential orientation of the periodic deviation over the evaluated portion of the interval (Figure S2c).

Qualified intervals satisfying the coverage requirements were classified as having *predominantly tangential periodic deviations* when *R*_tan_ *>* 0.75 and *periodic deviations with a substantial normal component* when *R*_tan_ ≤ 0.75. The criterion *R*_tan_ *>* 0.75 requires the summed squared tangential displacement to exceed three times the summed squared normal displacement. Intervals that failed the coverage requirements or for which *R*_tan_ could not be computed were left unclassified. Qualified intervals with predominantly tangential periodic deviations were used for the primary tests of the model predictions.

#### Validation of the estimation procedure with synthetic centerlines

To assess how accurately the 3D moving frame analysis estimated the amplitude and wavelength of transverse sinusoidal waves, we performed accuracy tests with noise-free synthetic centerlines of the form

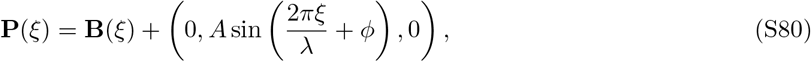

where **B**(*ξ*) = (*x*(*ξ*), 0, *z*(*ξ*)) is the known generating baseline, *ξ* is its arc-length parameter, and *A, λ*, and *ϕ* are the prescribed amplitude, wavelength, and phase, respectively. The displacement was applied in the *y* direction, perpendicular to the local tangent of **B**. Generating baselines were either straight, **B**(*ξ*) = (*ξ*, 0, 0), or obtained by arc-length parameterization of quadratic curves:

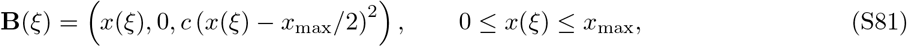

where *c* is the quadratic coefficient of the generating baseline, and *x*_max_ is the axial extent. For curved baselines (*c* ≠ 0), the minimum radius of curvature is 1*/*(2|*c*|). All synthetic centerlines were resampled along their own arc length with a target spacing of 0.1 mm and analyzed using the 3D moving frame analysis described above. The generating baselines were not supplied to the estimation procedure; a quadratic baseline was estimated from each synthetic centerline, as in the analysis of the MRA data. The true baseline length was *ℓ*_0_ = *ξ*_end_ − *ξ*_start_, where *ξ*_start_ and *ξ*_end_ are the arc-length positions of the first and last retained samples on the generating baseline. The corresponding true cycle count was *N*_cyc_ = *ℓ*_0_*/λ*. The fitted cycle count used in the periodic-fit qualification criteria was calculated from the estimated baseline length 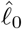 and estimated wavelength 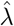 as 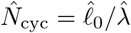. Accuracy was quantified by the absolute relative errors |*Â* − *A*|*/A* and 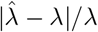, with *Â* and 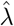defined in Eqs. S72 and S73; because the prescribed displacement was confined to one transverse direction, the true combined amplitude equaled *A*. All accuracy tests used a reference inner diameter of 2*r*_i_ = 1.4 mm, giving a lower wavelength bound of 2.8 mm. The test conditions are summarized in Table S8.

In the accuracy tests across amplitudes, wavelengths, and phases (156 cases), the lengths of the synthetic centerlines were adjusted numerically so that *N*_cyc_ attained its target value. Of these cases, 153 (98.1%) qualified. Among them, the median absolute relative errors were 0.46% (maximum, 6.1%) for combined amplitude and 0.035% (maximum, 1.7%) for wavelength. The three excluded cases had curved generating baselines (*c* = 0.005 mm^*−*1^), *A* = 0.5 mm, and *λ* = 60 mm, one at each phase. Their estimated wavelengths were approximately 270 mm, about 4.5 times the prescribed value, yielding 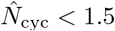. The accuracy tests across background curvatures compared 18 cases at *c* = 0.010 mm^*−*1^ with the 18 matched cases at *c* = 0.005 mm^*−*1^ from the accuracy tests across amplitudes, wavelengths, and phases, all with a target *N*_cyc_ of 3.0. All 18 cases qualified at *c* = 0.005 mm^*−*1^, compared with 16 of 18 at *c* = 0.010 mm^*−*1^. The two excluded cases had *λ* = 40 mm and *A* = 0.5 mm; their estimated wavelengths were approximately 154 mm, about 3.8 times the prescribed value, yielding 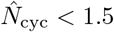. Among the qualified cases at *c* = 0.010 mm^*−*1^, the median absolute relative errors were 1.9% (maximum, 6.2%) for combined amplitude and 0.46% (maximum, 2.1%) for wavelength.

The accuracy tests across cycle counts used 432 windows extracted from longer synthetic centerlines, with target values of *N*_cyc_ from 0.5 to 3.0. Of these cases, 240 qualified, comprising all cases with target values of 1.5–3.0. Among them, the median absolute relative errors were 0.57% (maximum, 4.3%) for combined amplitude and 0.17% (maximum, 4.2%) for wavelength. At target values of 1.0 and 1.25 (48 cases each), the median absolute relative errors remained small (combined amplitude, 1.2% and 0.31%; wavelength, 0.44% and 0.34%), but larger errors occurred in some cases (maximum: combined amplitude, 35% and 12%; wavelength, 11% and 4.0%); all cases at these target values were excluded because 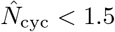. At target values below 1.0 (96 cases), the median absolute relative errors were 22% (maximum, 99%) for combined amplitude and 9.3% (maximum, 46%) for wavelength, and all of these cases were likewise excluded. Thus, the 1.5-cycle requirement served as a conservative acceptance margin that excluded less reliable estimates.

To assess whether estimation and qualification could themselves generate or distort wavelength–diameter scaling, we sampled, with replacement, pairs of measured inner diameter and estimated baseline length from the 11,368 of the 11,808 MCA centerline intervals that had valid values for both before qualification, and used them as the inner diameters and target true baseline lengths of synthetic centerlines, with the pairing either preserved or disrupted by permuting the lengths. Wavelengths were set to *λ* = 17.4 (2*r*_i_) (proportional inputs), where 17.4 is the theoretical ratio (main text, Section 3.2); to *λ* = 17.4 (2*r*_i_) *e*^*Z*^ with *Z ~ N* (0, 0.2^2^) (proportional inputs with scatter); or to proportional wavelengths permuted among intervals (permuted wavelengths). For each combination of scenario, pairing, and amplitude (*A* = 0.5 or 1.0 mm), 20 replicates of 500 straight-baseline centerlines with phases drawn uniformly from [0, 2*π*) were analyzed (120,000 cases), with the lower wavelength bound set to twice each sampled inner diameter. In each replicate, the coefficient of log inner diameter, corresponding to *β* in the analysis of the MRA data, was estimated by OLS regression of log wavelength on log inner diameter using the prescribed wavelengths and the estimated wavelengths before and after qualification. Of the 120,000 cases, 114,956 (95.8%) yielded numerically valid fits and 24,521 (20.4%) qualified. The median coefficients after qualification ranged from 0.996 to 1.000 for proportional inputs, from 0.919 to 1.007 for proportional inputs with scatter, and from 0.025 to 0.056 for permuted wavelengths, whereas those before qualification ranged from 0.627 to 0.775 for the two proportional scenarios and from 0.072 to 0.259 for permuted wavelengths (Table S6). Thus, under these simulated conditions, estimation and qualification preserved proportional scaling when present and did not generate an approximately proportional relationship when it was absent.

These tests assessed the accuracy of wavelength and amplitude estimation and the behavior of the qualification criteria under noise-free, idealized conditions; brain-surface directional classification, measurement noise, and more complex background geometries were not evaluated.

### S5.6 Statistical analysis

All parameters and thresholds, including those for vessel extraction, periodic fitting and qualification, brain-surface normal estimation, and directional classification, were fixed before the statistical analyses. Continuous variables are presented as median and IQR. For descriptive statistics, geometric metrics were summarized at the segment or interval level without participant-level aggregation. All logarithmic transformations used natural logarithms. Segment- and interval-level data were analyzed using OLS regression. To account for within-participant correlation from multiple vascular measurements, cluster-robust standard errors were calculated at the participant level (46). Regression coefficients are reported with 95% confidence intervals (CIs) based on these standard errors. For outcomes modeled on the log scale, coefficients for age were expressed as estimated relative differences per year, 100(*e*^*β*^ − 1)%. There were no missing values in the variables used in the regression analyses. Statistical significance for these models was assessed using two-sided Wald tests. A *P* value *<* 0.05 was considered statistically significant. *P* values from analyses explicitly described as exploratory were interpreted descriptively.

Ethnicity in the IXI Dataset was classified as White, Asian, Black, Chinese, and Other. For analysis, the “Asian” and “Chinese” categories were combined into a single “Asian” group (*n* = 20). Seven participants shared an ethnicity code that was not defined in the dataset documentation. These participants were retained in the analysis and assigned to an “Unknown” category. Because the Black, Other, and Unknown categories each included fewer than 10 participants, these categories were combined into a single “Black/Other/Unknown” group to improve model stability. This combined category was used in all regression models that included ethnic group as a covariate.

#### M1 elongation and tortuosity

The primary M1 geometric outcomes were centerline length, Euclidean distance, and excess-length ratio. Univariable regressions were fitted separately for age and each covariate. Multivariable linear regressions included age and the following covariates selected *a priori* based on their potential association with intracranial arterial morphology: sex, ethnic group, the cube root of total intracranial volume (TIV^1*/*3^), side, and M1 major branch type. Age was modeled as a single linear term.

Additional exploratory analyses were performed to assess potential nonlinearity in the association between age and M1 excess-length ratio *ρ*. First, we used a covariate-adjusted natural cubic spline function for age with two degrees of freedom. The covariates were the same as those used in the corresponding multivariable single-slope model. To describe how the rate of change in *ρ* varied with age, we numerically estimated the age derivative, *dρ/d*Age, from the fitted spline function. Second, we performed piecewise linear regression analyses using candidate breakpoints every 5 years from ages 30 to 55. For each candidate breakpoint, the model included age, a slope-change term, max(0, Age − breakpoint), and the same covariates as the multivariable single-slope model. The resulting Wald *P* values for the slope-change term were considered exploratory and were not adjusted for testing across multiple candidate breakpoints. Third, we used participant-level cluster bootstrap resampling (1,000 replicates) to describe the stability of selected breakpoint locations. In each bootstrap replicate, participants were sampled with replacement, and all M1 segment-level observations from the selected participants were retained. Candidate piecewise linear models were fitted within each replicate, and the breakpoint with the minimum Akaike information criterion was recorded as a descriptive summary. These breakpoint analyses were considered exploratory and were not interpreted as confirmatory evidence for a discrete onset or emergence age.

#### Periodic arterial waves

All analyses of periodic waves were restricted to intervals that met the periodic-fit qualification criteria and showed predominantly tangential periodic deviations. Each outcome was analyzed in univariable and multivariable models; all multivariable models were adjusted for sex, side, ethnic group, TIV^1*/*3^, and anatomical MCA segment.

For the wavelength–diameter scaling analysis, the log-transformed estimated wavelength was modeled as a function of the log-transformed inner diameter, with additional adjustment for age in the multivariable model. The estimated coefficient of log inner diameter and its 95% CI were compared with 1, the value expected for proportional scaling. The wavelength-to-inner-diameter ratio was summarized as the median and IQR. As a supplementary analysis, the log-transformed ratio was modeled as a function of age. For the amplitude analysis, the log-transformed combined amplitude was modeled as a function of age. The multivariable model additionally included the log-transformed wavelength and log-transformed inner diameter.

## Supplementary Figures

**Figure S1.**
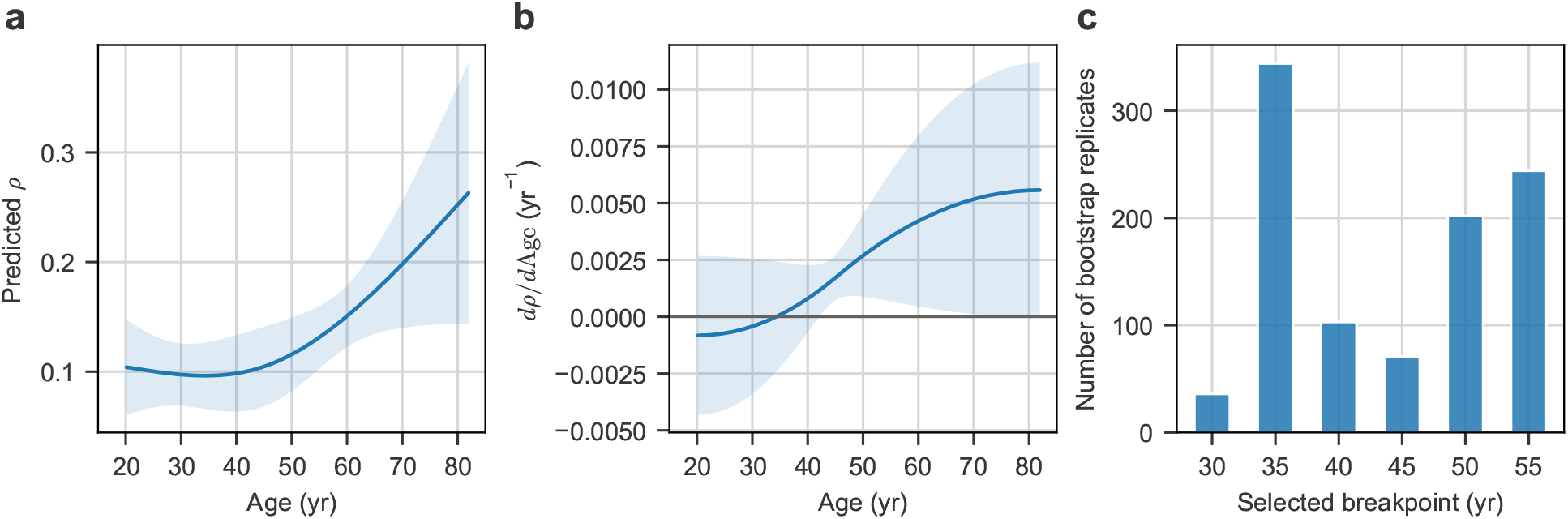
Exploratory analysis of age-related patterns in excess-length ratio *ρ*. (a) Covariate-adjusted natural cubic spline (df = 2) showing the relationship between age and excess-length ratio *ρ*. The solid blue line indicates the predicted mean, and the shaded area represents the pointwise 95% confidence interval (CI). The adjusted spline curve suggested that *ρ* was relatively flat in young adulthood and increased progressively thereafter. (b) Age-specific slope derived from the spline function, representing the rate of change in *ρ* with respect to age, *dρ/d*Age. The solid blue line indicates the estimated slope, and the shaded area represents the pointwise 95% CI. The slope estimates shifted toward positive values from early middle age (approximately 35–45 years) onward. (c) Distribution of selected breakpoint locations from participant-level cluster bootstrap analysis (1,000 replicates). In each replicate, the breakpoint was selected by minimizing the Akaike information criterion across candidate piecewise linear models. The most frequently selected breakpoint was 35 years, although the selected breakpoints were distributed across the candidate range.

**Figure S2.**
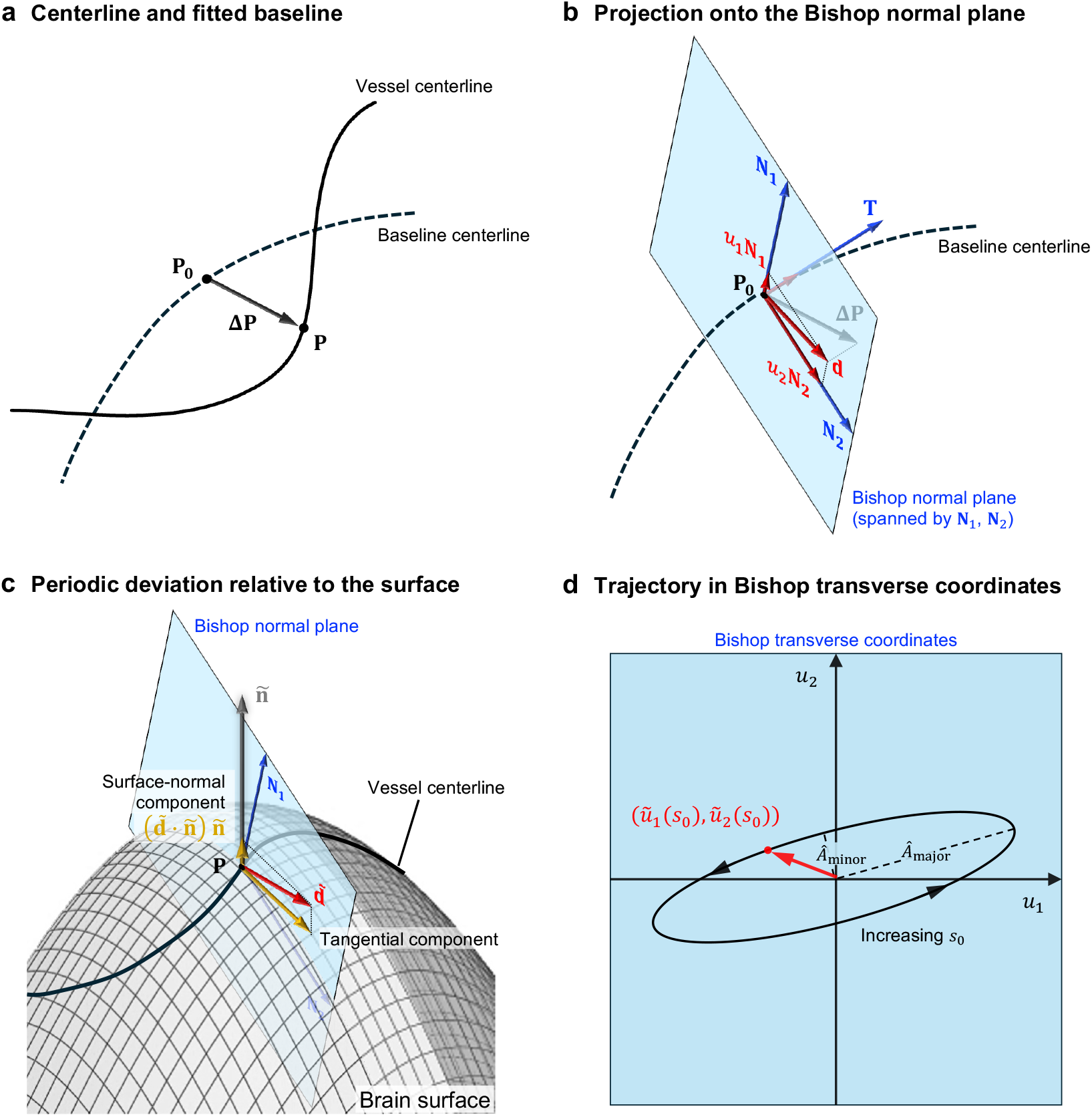
Schematic of the geometric residual and its transverse projection using the Bishop frame. (a) The pointwise geometric residual is defined as Δ**P**(*s*) = **P**(*s*) − **P**_0_(*s*) between corresponding points on the vessel centerline (solid line) and its fitted baseline centerline (dashed line). (b) In the local Bishop frame along the baseline centerline, Δ**P** is orthogonally projected onto the normal plane spanned by **N**_1_ and **N**_2_, yielding **d** = *u*_1_**N**_1_ + *u*_2_**N**_2_. **T** denotes the local unit tangent of the baseline. (c) The periodic deviation 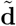, defined in the baseline Bishop frame, is compared with the effective surface-normal vector **ñ** at the corresponding vessel centerline point **P**. The surface-normal component 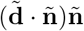 is used to calculate the tangential squared-displacement fraction *R*_tan_ over points with evaluable normals. (d) The fitted sinusoidal coordinate pair *ũ*_1_(*s*_0_), *ũ*_2_(*s*_0_) traces an elliptical locus in Bishop transverse coordinates as the baseline arc-length parameter *s*_0_ varies. Its semiaxis lengths are the singular values *Á*_major_ and *Á*_minor_ of the fitted coefficient matrix *M*; the combined amplitude is 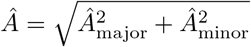. Ellipticity is defined as *Á*_minor_*/Á*_major_.

**Figure S3.**
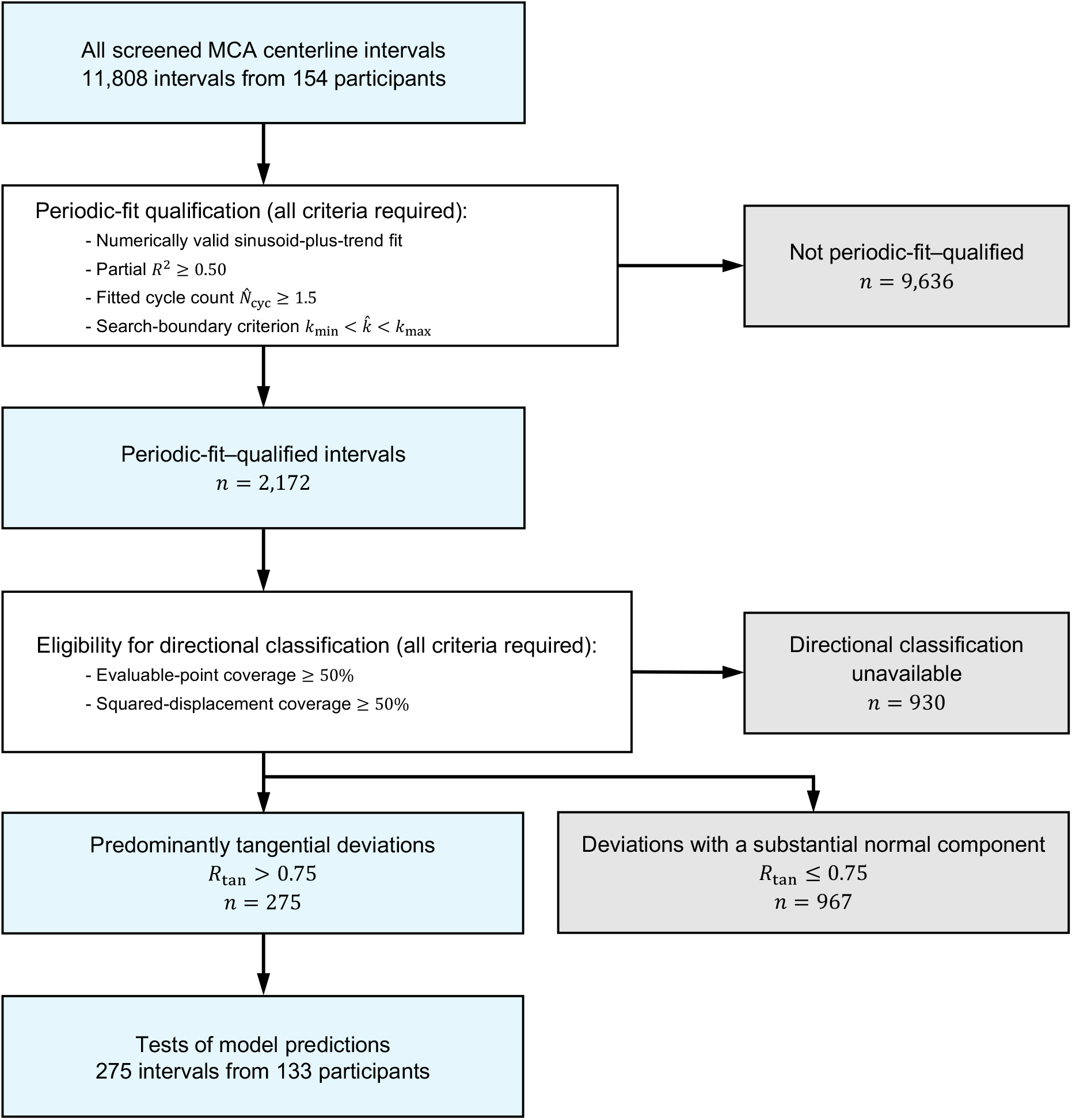
Workflow for periodic-fit qualification and directional classification. Screened middle cerebral artery (MCA) intervals underwent periodic-fit qualification followed by directional classification of the periodic deviations relative to the brain surface using the criteria shown. Intervals with insufficient coverage or without a valid tangential squared-displacement fraction *R*_tan_ were left unclassified. Qualified intervals with predominantly tangential periodic deviations were used for the primary tests of the model predictions.

**Figure S4.**
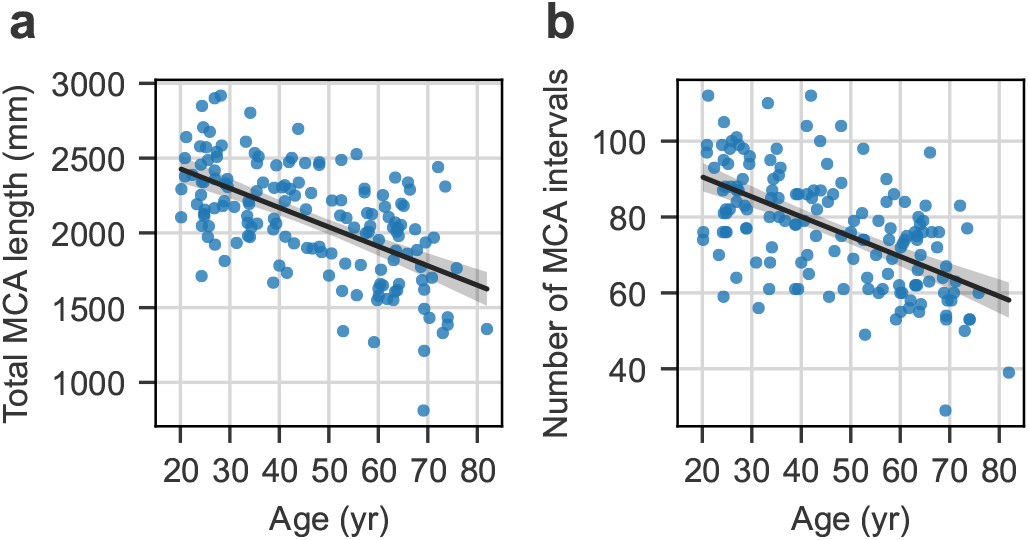
Association between age and arterial depiction on magnetic resonance angiography. Exploratory analyses of age associations with (a) total reconstructed middle cerebral artery (MCA) length and (b) the total number of detected MCA intervals. Each point represents one participant (*n* = 154). Black lines indicate univariable ordinary least-squares regression fits, and gray shaded bands represent pointwise 95% confidence intervals (CIs) for the fitted mean responses. The unadjusted age coefficients were −12.97 mm*/*yr (95% CI, − 15.85 to − 10.08; *P* < 0.001) for total MCA length and −0.524 intervals/yr (95% CI, − 0.643 to −0.405; *P* < 0.001) for interval count. Multivariable models were adjusted for sex, ethnic group, and the cube root of total intracranial volume, with additional adjustment for total MCA length in the interval-count model. The adjusted age coefficients were −12.78 mm*/*yr (95% CI, −15.53 to −10.02; *P* < 0.001) and −0.133 intervals/yr (95% CI, −0.230 to 0.037; *P* = 0.007), respectively. All analyses were performed at the participant level, and *P* values for the age coefficients were obtained from two-sided *t* tests.

**Figure S5.**
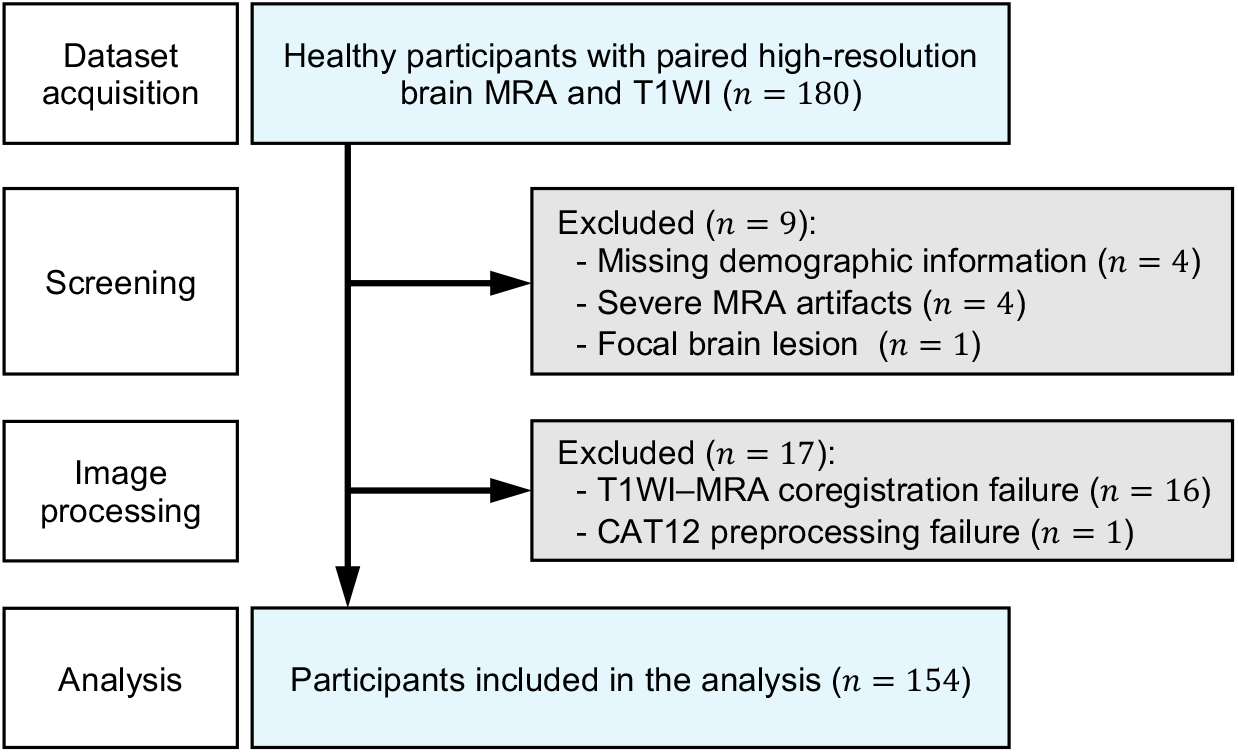
Flowchart illustrating the participant selection process. *MRA, magnetic resonance angiography; T1WI, T1-weighted images; CAT12, Computational Anatomy Toolbox 12*.

**Figure S6.**
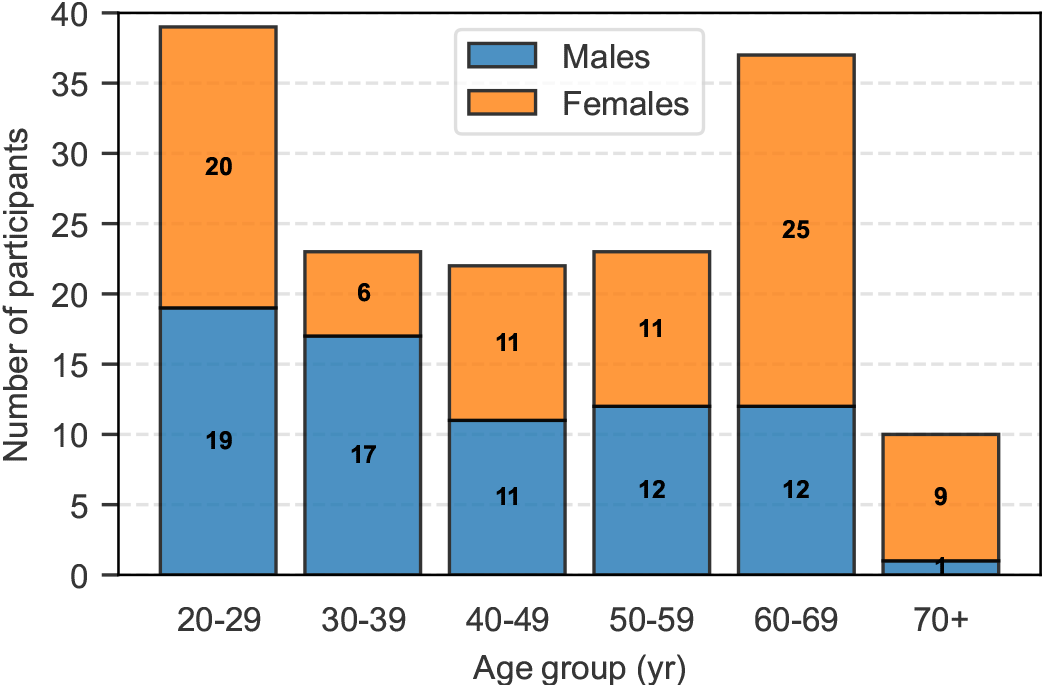
Age and sex distribution of the study participants. Participants ranged in age from 20 to 81 years, covering a broad span of adulthood. Bars show the number of male and female participants in each age group, with counts shown inside each bar (*n* = 154).

**Figure S7.**
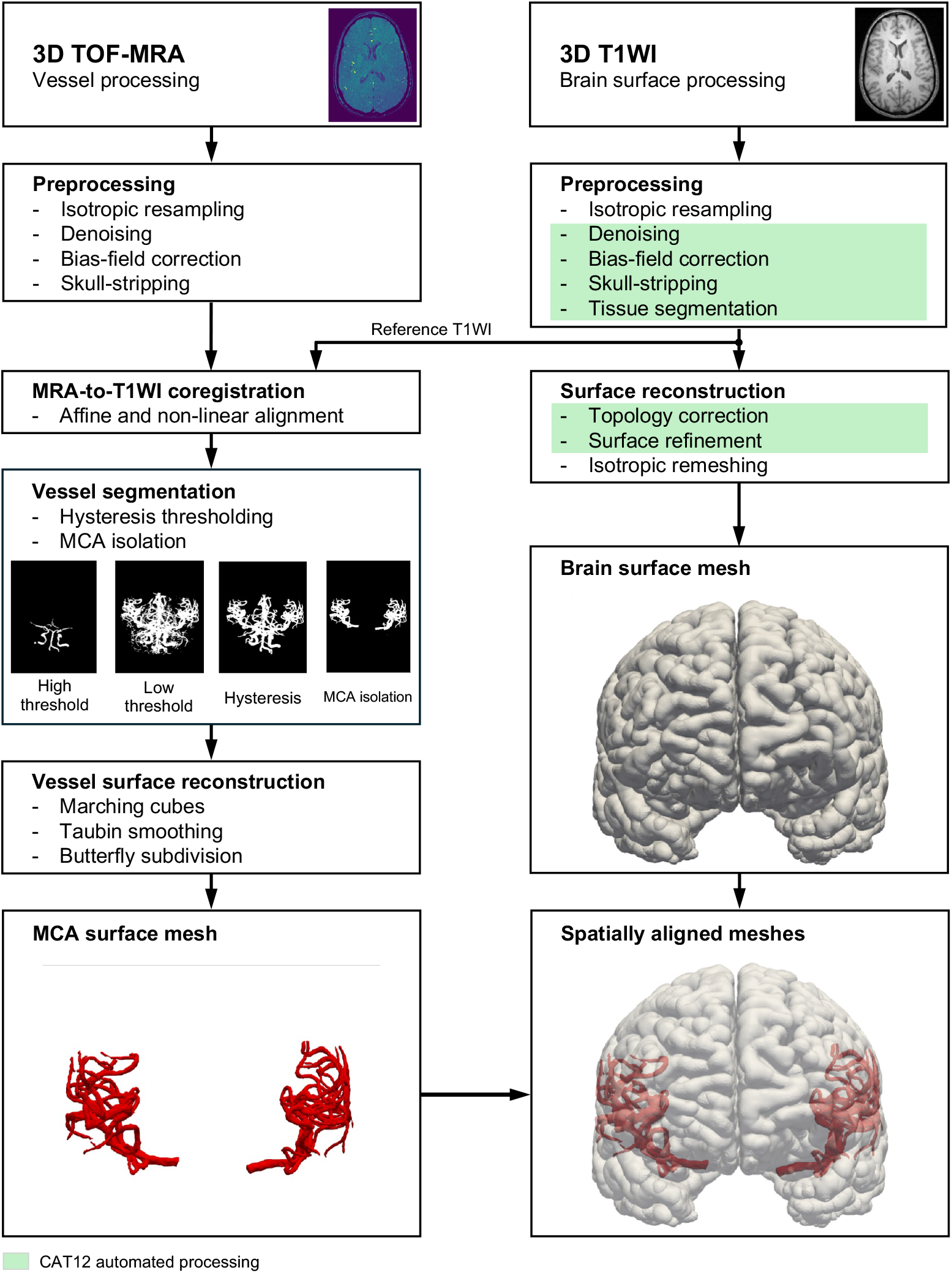
Image processing pipeline for cerebral artery and brain surface meshes. MRA volumes were coregistered to preprocessed T1WI to obtain spatially aligned MCA and brain surface meshes. The green boxes indicate automated processing steps performed using CAT12. *3D, three-dimensional; TOF-MRA, time-of-flight magnetic resonance angiography; T1WI, T1-weighted images; CAT12, Computational Anatomy Toolbox 12; MCA, middle cerebral artery*.

**Figure S8.**
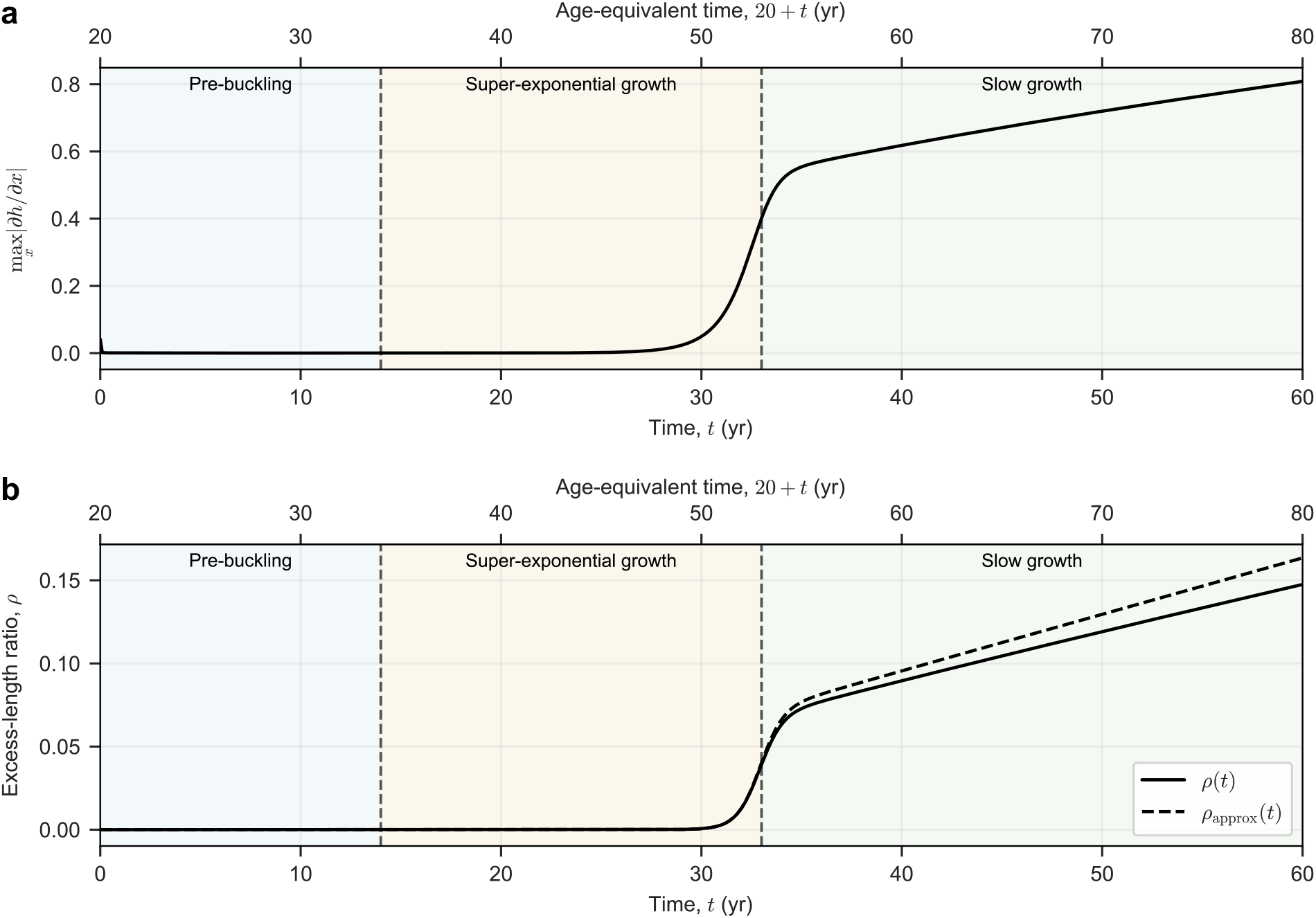
*A posteriori* assessment of the small-slope approximation during numerical simulation. (a) Maximum local slope, max_*x*_ |*∂h/∂x*|. (b) Comparison of the excess-length ratio, *ρ*(*t*) (solid line), calculated using the exact arc-length formula, with its quadratic approximation, *ρ*_approx_(*t*) (dashed line), used in the axial-force relation. Both were evaluated from the same simulated profile. The increase of max_*x*_ *∂h/∂x* to order unity and the widening separation between the two excess-length ratios during the late slow-growth stage indicate that the small-slope condition max_*x*_ |*∂h/∂x*| *≪* 1 is no longer satisfied. Colored backgrounds and vertical dashed lines indicate the three stages defined in Figure 3.

**Figure S9.**
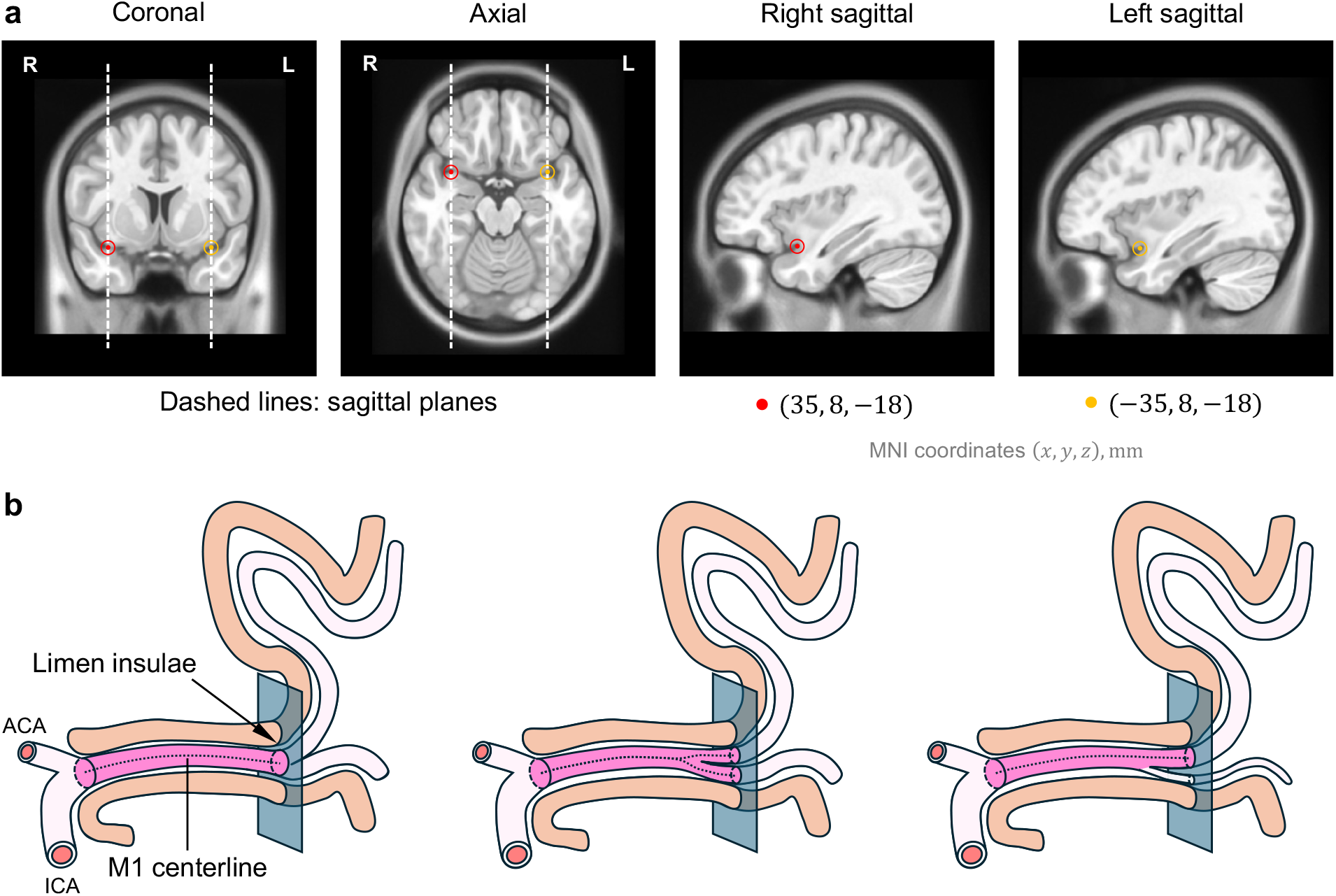
Limen insulae reference points and M1 major branch types. (a) Limen insulae reference points and sagittal planes. From left to right: coronal, axial, right sagittal, and left sagittal views. Red and yellow markers indicate reference points for the right and left limen insulae at Montreal Neurological Institute (MNI) coordinates (*x, y, z*) = (35, 8, −18) and ( −35, 8, −18) mm, respectively. Dashed lines in the coronal and axial views indicate the sagittal planes through these points. The MNI-defined planes were mapped into each participant’s native T1-weighted image space to obtain the reference surfaces used for M1 measurements. (b) M1 major branch types and measurement paths. Left, single-trunk type; middle, codominant-trunk type; right, dominant-trunk type. Paths extend from the internal carotid artery (ICA) terminus to the limen insulae-based reference surface (light-blue slab). Pink highlights indicate the measured paths, and dotted lines indicate their centerlines. *ACA, anterior cerebral artery*.

**Figure S10.**
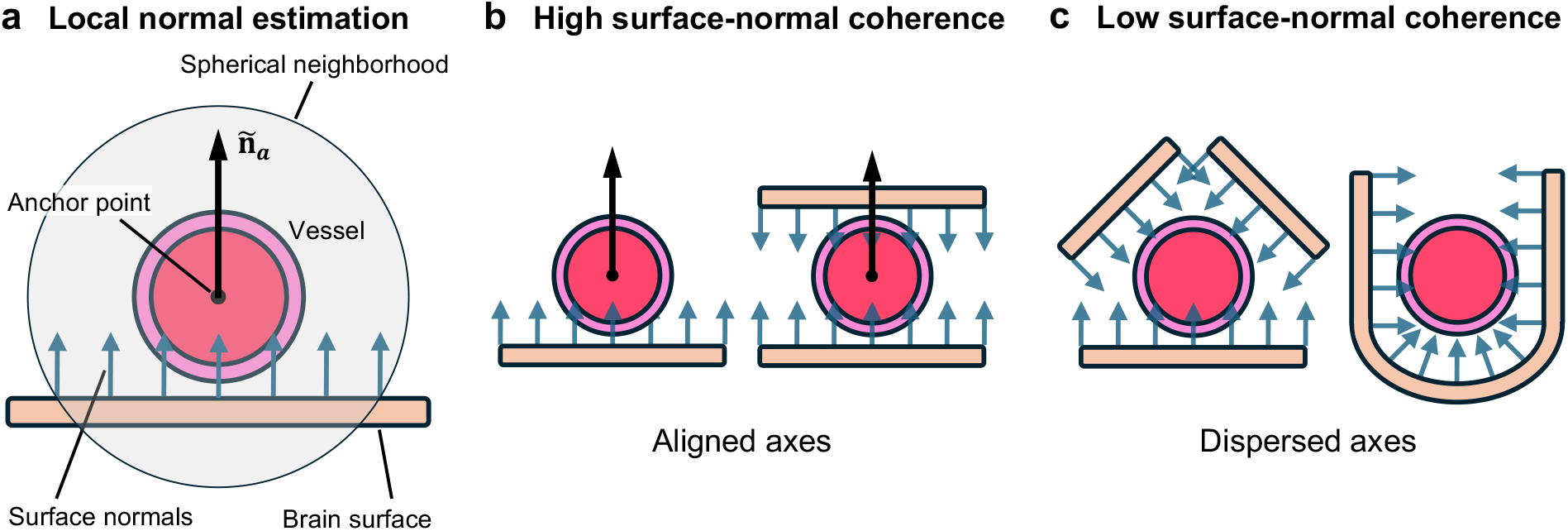
Schematic of local brain-surface normal estimation and surface-normal coherence. (a) Brain-surface normals are sampled within a local spherical neighborhood around each vessel centerline anchor point. Their dominant orientation defines the effective surface-normal axis, represented by the unit vector **ñ**_*a*_, and their degree of alignment defines the surface-normal coherence *C*_*a*_. (b) A single surface or two approximately parallel surfaces can yield high *C*_*a*_ because their normal axes are closely aligned, regardless of vector sign. (c) Complex local surface configurations, such as multiple nonparallel brain-surface patches and sulcal fundi, can yield widely dispersed normal axes and therefore low *C*_*a*_. Low-*C*_*a*_ estimates are not used directly for directional evaluation.

## Supplementary Tables

**Table S1.**
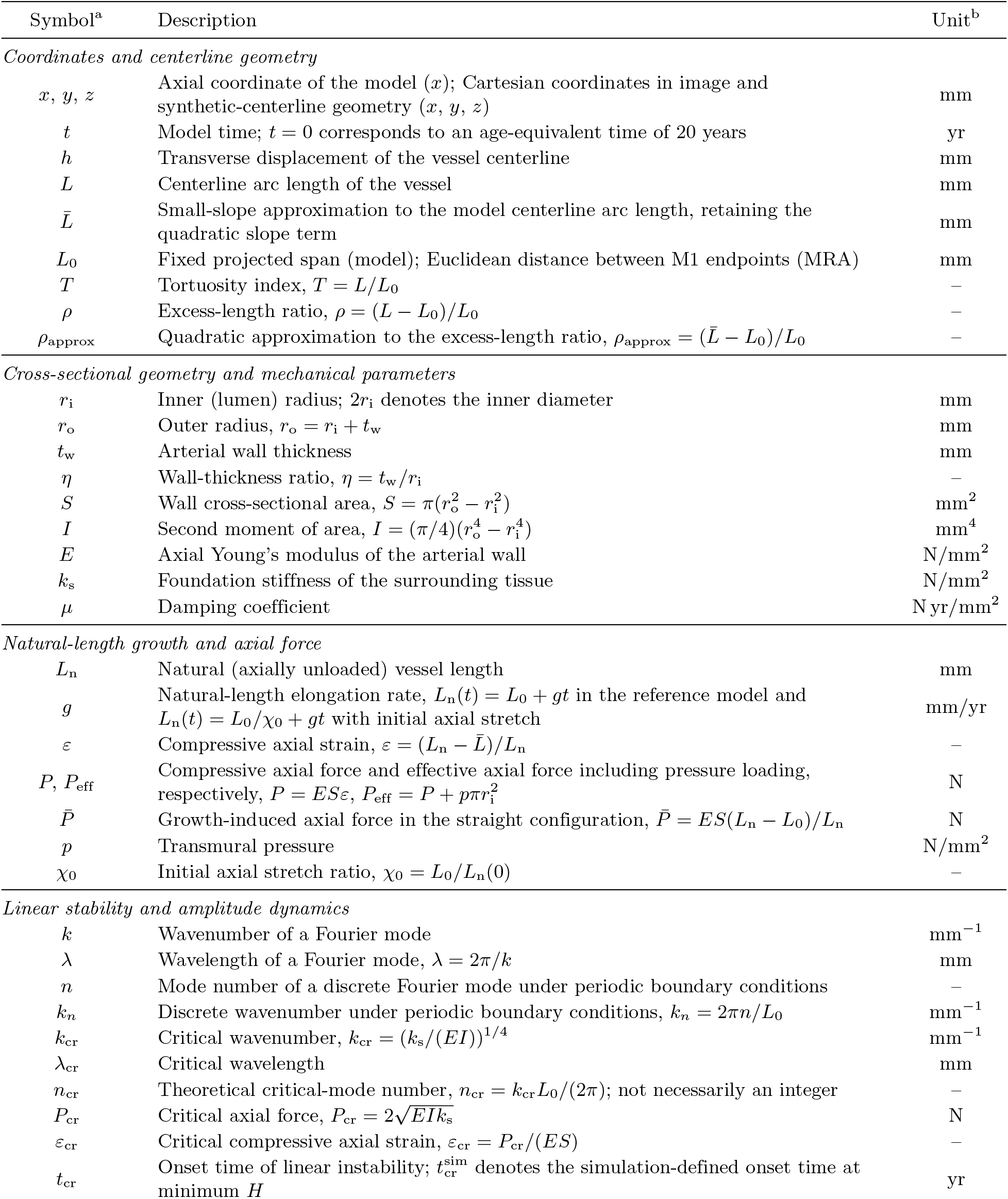
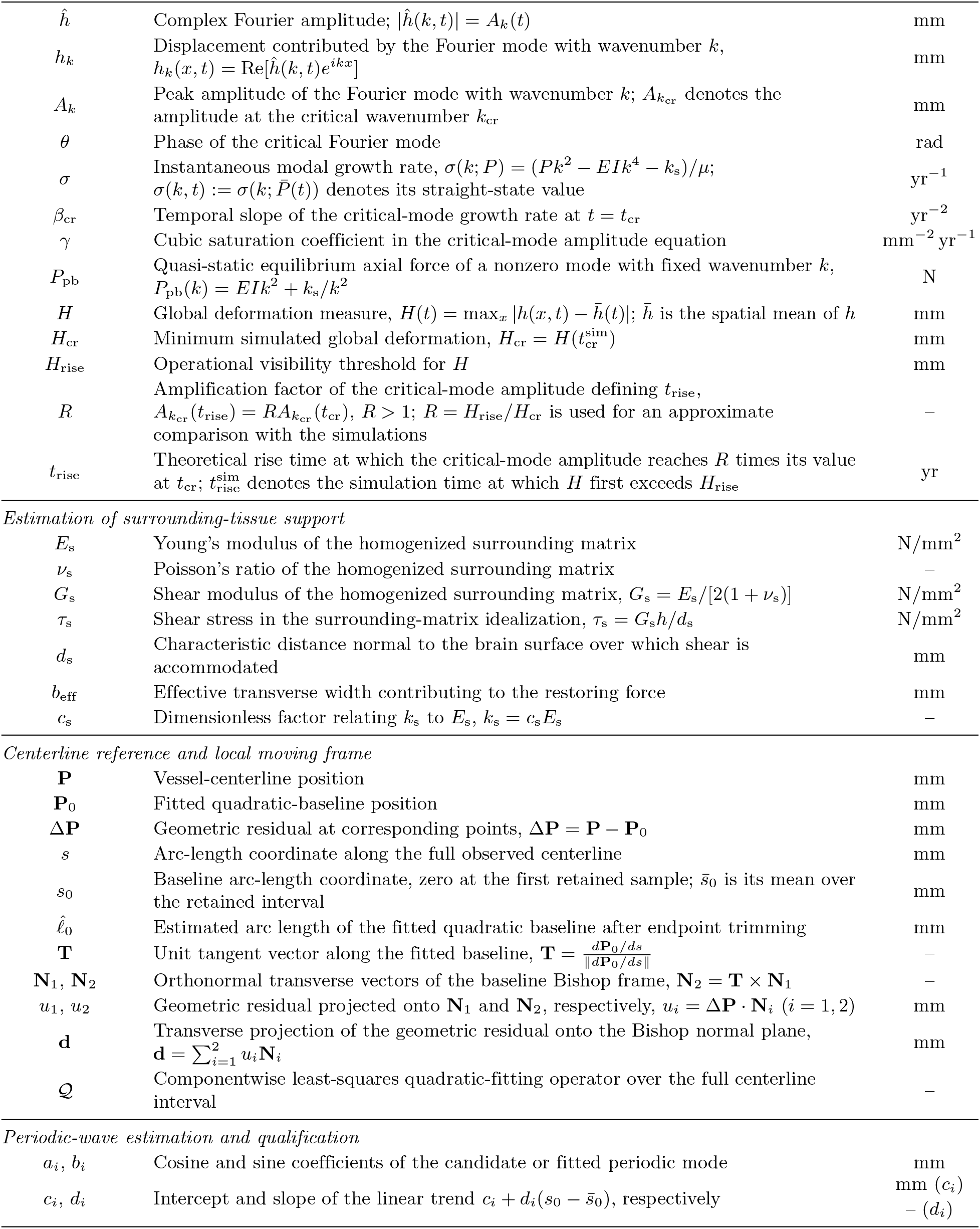
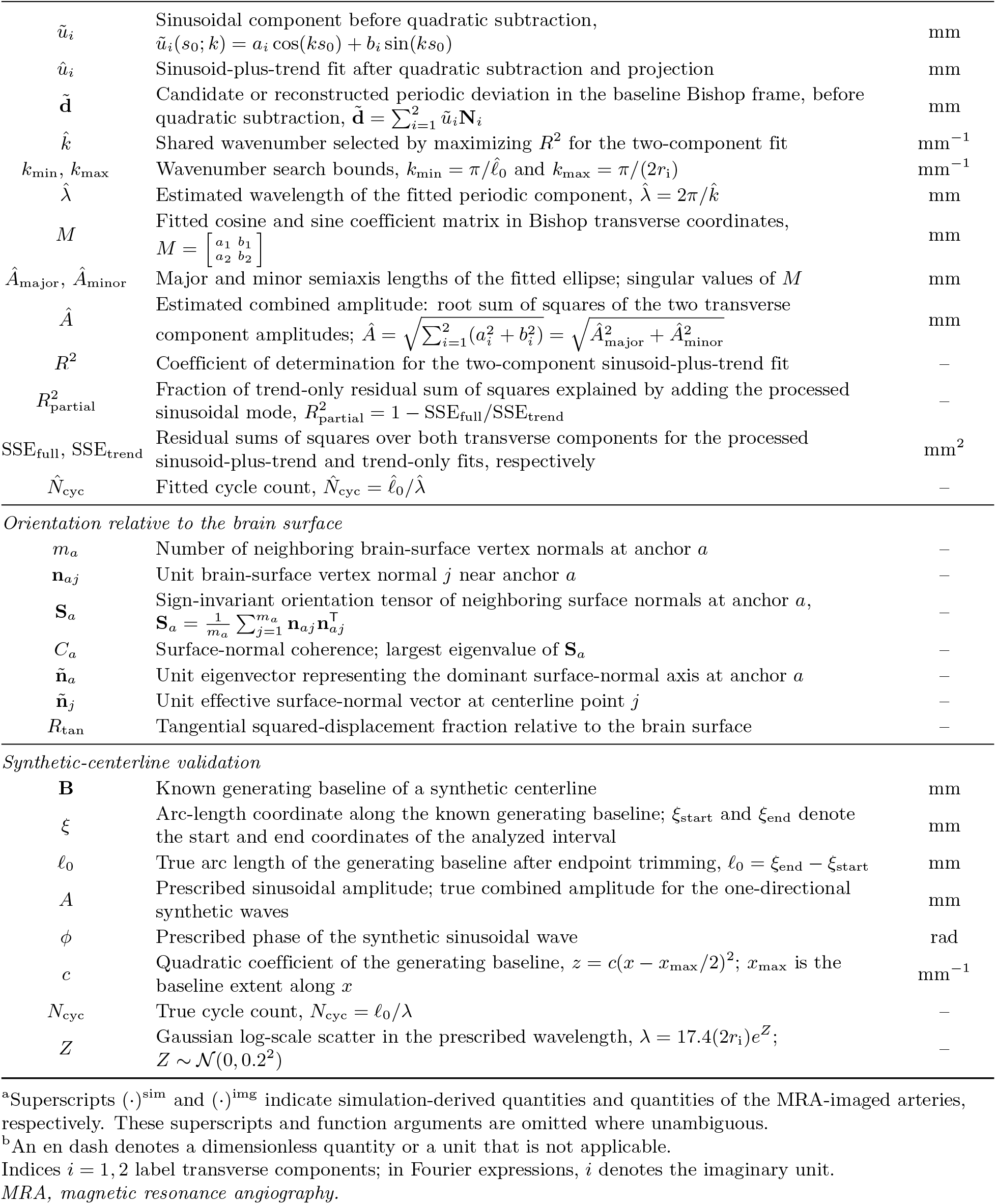
List of variables and parameters used in the SI Appendix.

| Symbol<sup>a</sup> | Description | Unit<sup>b</sup> |
|---|---|---|
| *Coordinates and centerline geometry* |  |  |
| x, y, z | Axial coordinate of the model (x); Cartesian coordinates in image and synthetic-centerline geometry (x, y, z) | mm |
| t | Model time; t = 0 corresponds to an age-equivalent time of 20 years | yr |
| h | Transverse displacement of the vessel centerline | mm |
| L | Centerline arc length of the vessel | mm |
| L̄ | Small-slope approximation to the model centerline arc length, retaining the quadratic slope term | mm |
| L<sub>0</sub> | Fixed projected span (model); Euclidean distance between M1 endpoints (MRA) | mm |
| T | Tortuosity index, T = L/L<sub>0</sub> | – |
| ρ | Excess-length ratio, ρ = (L − L<sub>0</sub>)/L<sub>0</sub> | – |
| ρ<sub>approx</sub> | Quadratic approximation to the excess-length ratio, ρ<sub>approx</sub> = (L̄ − L<sub>0</sub>)/L<sub>0</sub> | – |
| *Cross-sectional geometry and mechanical parameters* |  |  |
| r<sub>i</sub> | Inner (lumen) radius; 2r<sub>i</sub> denotes the inner diameter | mm |
| r<sub>o</sub> | Outer radius, r<sub>o</sub> = r<sub>i</sub> + t<sub>w</sub> | mm |
| t<sub>w</sub> | Arterial wall thickness | mm |
| η | Wall-thickness ratio, η = t<sub>w</sub>/r<sub>i</sub> | – |
| S | Wall cross-sectional area, S = π(r<sub>o</sub><sup>2</sup> − r<sub>i</sub><sup>2</sup>) | mm<sup>2</sup> |
| I | Second moment of area, I = (π/4)(r<sub>o</sub><sup>4</sup> − r<sub>i</sub><sup>4</sup>) | mm<sup>4</sup> |
| E | Axial Young’s modulus of the arterial wall | N/mm<sup>2</sup> |
| k<sub>s</sub> | Foundation stiffness of the surrounding tissue | N/mm<sup>2</sup> |
| μ | Damping coefficient | N yr/mm<sup>2</sup> |
| *Natural-length growth and axial force* |  |  |
| L<sub>n</sub> | Natural (axially unloaded) vessel length | mm |
| g | Natural-length elongation rate, L<sub>n</sub>(t) = L<sub>0</sub> + gt in the reference model and L<sub>n</sub>(t) = L<sub>0</sub>/χ<sub>0</sub> + gt with initial axial stretch | mm/yr |
| ε | Compressive axial strain, ε = (L<sub>n</sub> − L̄)/L<sub>n</sub> | – |
| P, P<sub>eff</sub> | Compressive axial force and effective axial force including pressure loading, respectively, P = ESε, P<sub>eff</sub> = P + pπr<sub>i</sub><sup>2</sup> | N |
| P̄ | Growth-induced axial force in the straight configuration, P̄ = ES(L<sub>n</sub> − L<sub>0</sub>)/L<sub>n</sub> | N |
| p | Transmural pressure | N/mm<sup>2</sup> |
| χ<sub>0</sub> | Initial axial stretch ratio, χ<sub>0</sub> = L<sub>0</sub>/L<sub>n</sub>(0) | – |
| *Linear stability and amplitude dynamics* |  |  |
| k | Wavenumber of a Fourier mode | mm<sup>−1</sup> |
| λ | Wavelength of a Fourier mode, λ = 2π/k | mm |
| n | Mode number of a discrete Fourier mode under periodic boundary conditions | – |
| k<sub>n</sub> | Discrete wavenumber under periodic boundary conditions, k<sub>n</sub> = 2πn/L<sub>0</sub> | mm<sup>−1</sup> |
| k<sub>cr</sub> | Critical wavenumber, k<sub>cr</sub> = (k<sub>s</sub>/(EI))<sup>1/4</sup> | mm<sup>−1</sup> |
| λ<sub>cr</sub> | Critical wavelength | mm |
| n<sub>cr</sub> | Theoretical critical-mode number, n<sub>cr</sub> = k<sub>cr</sub>L<sub>0</sub>/(2π); not necessarily an integer | – |
| P<sub>cr</sub> | Critical axial force, $P_{\mathrm{cr}}=2\sqrt{EIk_{\mathrm{s}}}$ | N |
| ε<sub>cr</sub> | Critical compressive axial strain, ε<sub>cr</sub> = P<sub>cr</sub>/(ES) | – |
| t<sub>cr</sub> | Onset time of linear instability; t<sub>cr</sub><sup>sim</sup> denotes the simulation-defined onset time at minimum H | yr |
| ĥ | Complex Fourier amplitude; \|ĥ(k, t)\| = A<sub>k</sub>(t) | mm |
| h<sub>k</sub> | Displacement contributed by the Fourier mode with wavenumber k, h<sub>k</sub>(x, t) = Re[ĥ(k, t)e<sup>ikx</sup>] | mm |
| A<sub>k</sub> | Peak amplitude of the Fourier mode with wavenumber k; A<sub>k<sub>cr</sub></sub> denotes the amplitude at the critical wavenumber k<sub>cr</sub> | mm |
| θ | Phase of the critical Fourier mode | rad |
| σ | Instantaneous modal growth rate, σ(k; P) = (Pk<sup>2</sup> − EIk<sup>4</sup> − k<sub>s</sub>)/μ; σ(k, t) := σ(k; P̄(t)) denotes its straight-state value | yr<sup>−1</sup> |
| β<sub>cr</sub> | Temporal slope of the critical-mode growth rate at t = t<sub>cr</sub> | yr<sup>−2</sup> |
| γ | Cubic saturation coefficient in the critical-mode amplitude equation | mm<sup>−2</sup> yr<sup>−1</sup> |
| P<sub>pb</sub> | Quasi-static equilibrium axial force of a nonzero mode with fixed wavenumber k, P<sub>pb</sub>(k) = EIk<sup>2</sup> + k<sub>s</sub>/k<sup>2</sup> | N |
| H | Global deformation measure, H(t) = max<sub>x</sub> \|h(x, t) − h̄(t)\|; h̄ is the spatial mean of h | mm |
| H<sub>cr</sub> | Minimum simulated global deformation, H<sub>cr</sub> = H(t<sub>cr</sub><sup>sim</sup>) | mm |
| H<sub>rise</sub> | Operational visibility threshold for H | mm |
| R | Amplification factor of the critical-mode amplitude defining t<sub>rise</sub>, A<sub>k<sub>cr</sub></sub>(t<sub>rise</sub>) = RA<sub>k<sub>cr</sub></sub>(t<sub>cr</sub>), R > 1; R = H<sub>rise</sub>/H<sub>cr</sub> is used for an approximate comparison with the simulations | – |
| t<sub>rise</sub> | Theoretical rise time at which the critical-mode amplitude reaches R times its value at t<sub>cr</sub>; t<sub>rise</sub><sup>sim</sup> denotes the simulation time at which H first exceeds H<sub>rise</sub> | yr |
| *Estimation of surrounding-tissue support* |  |  |
| E<sub>s</sub> | Young’s modulus of the homogenized surrounding matrix | N/mm<sup>2</sup> |
| ν<sub>s</sub> | Poisson’s ratio of the homogenized surrounding matrix | – |
| G<sub>s</sub> | Shear modulus of the homogenized surrounding matrix, G<sub>s</sub> = E<sub>s</sub>/[2(1 + ν<sub>s</sub>)] | N/mm<sup>2</sup> |
| τ<sub>s</sub> | Shear stress in the surrounding-matrix idealization, τ<sub>s</sub> = G<sub>s</sub>h/d<sub>s</sub> | N/mm<sup>2</sup> |
| d<sub>s</sub> | Characteristic distance normal to the brain surface over which shear is accommodated | mm |
| b<sub>eff</sub> | Effective transverse width contributing to the restoring force | mm |
| c<sub>s</sub> | Dimensionless factor relating k<sub>s</sub> to E<sub>s</sub>, k<sub>s</sub> = c<sub>s</sub>E<sub>s</sub> | – |
| *Centerline reference and local moving frame* |  |  |
| **P** | Vessel-centerline position | mm |
| **P**<sub>0</sub> | Fitted quadratic-baseline position | mm |
| **ΔP** | Geometric residual at corresponding points, **ΔP** = **P** − **P**<sub>0</sub> | mm |
| s | Arc-length coordinate along the full observed centerline | mm |
| s<sub>0</sub> | Baseline arc-length coordinate, zero at the first retained sample; s̄<sub>0</sub> is its mean over the retained interval | mm |
| ℓ̂<sub>0</sub> | Estimated arc length of the fitted quadratic baseline after endpoint trimming | mm |
| **T** | Unit tangent vector along the fitted baseline, $\mathbf{T}=\frac{d\mathbf{P}_0/ds}{\lVert d\mathbf{P}_0/ds\rVert}$ | – |
| **N**<sub>1</sub>, **N**<sub>2</sub> | Orthonormal transverse vectors of the baseline Bishop frame, **N**<sub>2</sub> = **T** × **N**<sub>1</sub> | – |
| u<sub>1</sub>, u<sub>2</sub> | Geometric residual projected onto **N**<sub>1</sub> and **N**<sub>2</sub>, respectively, u<sub>i</sub> = **ΔP** · **N**<sub>i</sub> (i = 1, 2) | mm |
| **d** | Transverse projection of the geometric residual onto the Bishop normal plane, $\mathbf{d}=\sum_{i=1}^{2}u_i\mathbf{N}_i$ | mm |
| 𝒬 | Componentwise least-squares quadratic-fitting operator over the full centerline interval | – |
| *Periodic-wave estimation and qualification* |  |  |
| a<sub>i</sub>, b<sub>i</sub> | Cosine and sine coefficients of the candidate or fitted periodic mode | mm |
| c<sub>i</sub>, d<sub>i</sub> | Intercept and slope of the linear trend c<sub>i</sub> + d<sub>i</sub>(s<sub>0</sub> − s̄<sub>0</sub>), respectively | mm (c<sub>i</sub>)<br>– (d<sub>i</sub>) |
| ũ<sub>i</sub> | Sinusoidal component before quadratic subtraction, ũ<sub>i</sub>(s<sub>0</sub>; k) = a<sub>i</sub> cos(ks<sub>0</sub>) + b<sub>i</sub> sin(ks<sub>0</sub>) | mm |
| û<sub>i</sub> | Sinusoid-plus-trend fit after quadratic subtraction and projection | mm |
| **d̃** | Candidate or reconstructed periodic deviation in the baseline Bishop frame, before quadratic subtraction, $\tilde{\mathbf{d}}=\sum_{i=1}^{2}\tilde{u}_i\mathbf{N}_i$ | mm |
| k̂ | Shared wavenumber selected by maximizing R<sup>2</sup> for the two-component fit | mm<sup>−1</sup> |
| k<sub>min</sub>, k<sub>max</sub> | Wavenumber search bounds, k<sub>min</sub> = π/ℓ̂<sub>0</sub> and k<sub>max</sub> = π/(2r<sub>i</sub>) | mm<sup>−1</sup> |
| λ̂ | Estimated wavelength of the fitted periodic component, λ̂ = 2π/k̂ | mm |
| M | Fitted cosine and sine coefficient matrix in Bishop transverse coordinates, $M=\begin{bmatrix}a_1&b_1\\a_2&b_2\end{bmatrix}$ | mm |
| Â<sub>major</sub>, Â<sub>minor</sub> | Major and minor semiaxis lengths of the fitted ellipse; singular values of M | mm |
| Â | Estimated combined amplitude: root sum of squares of the two transverse component amplitudes; $\hat{A}=\sqrt{\sum_{i=1}^{2}(a_i^2+b_i^2)}=\sqrt{\hat{A}_{\mathrm{major}}^2+\hat{A}_{\mathrm{minor}}^2}$ | mm |
| R<sup>2</sup> | Coefficient of determination for the two-component sinusoid-plus-trend fit | – |
| R<sup>2</sup><sub>partial</sub> | Fraction of trend-only residual sum of squares explained by adding the processed sinusoidal mode, R<sup>2</sup><sub>partial</sub> = 1 − SSE<sub>full</sub>/SSE<sub>trend</sub> | – |
| SSE<sub>full</sub>, SSE<sub>trend</sub> | Residual sums of squares over both transverse components for the processed sinusoid-plus-trend and trend-only fits, respectively | mm<sup>2</sup> |
| N̂<sub>cyc</sub> | Fitted cycle count, N̂<sub>cyc</sub> = ℓ̂<sub>0</sub>/λ̂ | – |
| *Orientation relative to the brain surface* |  |  |
| m<sub>a</sub> | Number of neighboring brain-surface vertex normals at anchor a | – |
| **n**<sub>aj</sub> | Unit brain-surface vertex normal j near anchor a | – |
| **S**<sub>a</sub> | Sign-invariant orientation tensor of neighboring surface normals at anchor a, $\mathbf{S}_a=\frac{1}{m_a}\sum_{j=1}^{m_a}\mathbf{n}_{aj}\mathbf{n}_{aj}^{\mathsf{T}}$ | – |
| C<sub>a</sub> | Surface-normal coherence; largest eigenvalue of **S**<sub>a</sub> | – |
| **ñ**<sub>a</sub> | Unit eigenvector representing the dominant surface-normal axis at anchor a | – |
| **ñ**<sub>j</sub> | Unit effective surface-normal vector at centerline point j | – |
| R<sub>tan</sub> | Tangential squared-displacement fraction relative to the brain surface | – |
| *Synthetic-centerline validation* |  |  |
| **B** | Known generating baseline of a synthetic centerline | mm |
| ξ | Arc-length coordinate along the known generating baseline; ξ<sub>start</sub> and ξ<sub>end</sub> denote the start and end coordinates of the analyzed interval | mm |
| ℓ<sub>0</sub> | True arc length of the generating baseline after endpoint trimming, ℓ<sub>0</sub> = ξ<sub>end</sub> − ξ<sub>start</sub> | mm |
| A | Prescribed sinusoidal amplitude; true combined amplitude for the one-directional synthetic waves | mm |
| ϕ | Prescribed phase of the synthetic sinusoidal wave | rad |
| c | Quadratic coefficient of the generating baseline, z = c(x − x<sub>max</sub>/2)<sup>2</sup>; x<sub>max</sub> is the baseline extent along x | mm<sup>−1</sup> |
| N<sub>cyc</sub> | True cycle count, N<sub>cyc</sub> = ℓ<sub>0</sub>/λ | – |
| Z | Gaussian log-scale scatter in the prescribed wavelength, λ = 17.4(2r<sub>i</sub>)e<sup>Z</sup>; Z ∼ 𝒩(0, 0.2<sup>2</sup>) | – |

<sup>a</sup>Superscripts (·)<sup>sim</sup> and (·)<sup>img</sup> indicate simulation-derived quantities and quantities of the MRA-imaged arteries, respectively. These superscripts and function arguments are omitted where unambiguous.
<sup>b</sup>An en dash denotes a dimensionless quantity or a unit that is not applicable.
Indices i = 1, 2 label transverse components; in Fourier expressions, i denotes the imaginary unit.
*MRA, magnetic resonance angiography.*

**Table S2.** Morphological characteristics of M1 segments.

| Characteristic (*n* = 349) | Values |
|---|---|
| Centerline length, *L* (mm) | 20.1 (18.0–22.8) |
| Euclidean distance, *L*<sub>0</sub> (mm) | 18.0 (16.5–19.6) |
| Tortuosity index, *T* | 1.10 (1.06–1.16) |
| Excess-length ratio, *ρ* | 0.10 (0.06–0.16) |
| Inner radius, *r*<sub>i</sub> (mm) | 1.55 (1.46–1.63) |

Data are presented as median (interquartile range) at the segment level. The 349 M1 segments were obtained from 154 participants; some hemispheres contributed more than one M1 segment under the anatomical definition used here (Section S5.4). Multiple segments from the same participant were retained to characterize the segment-level distribution.

**Table S3.**
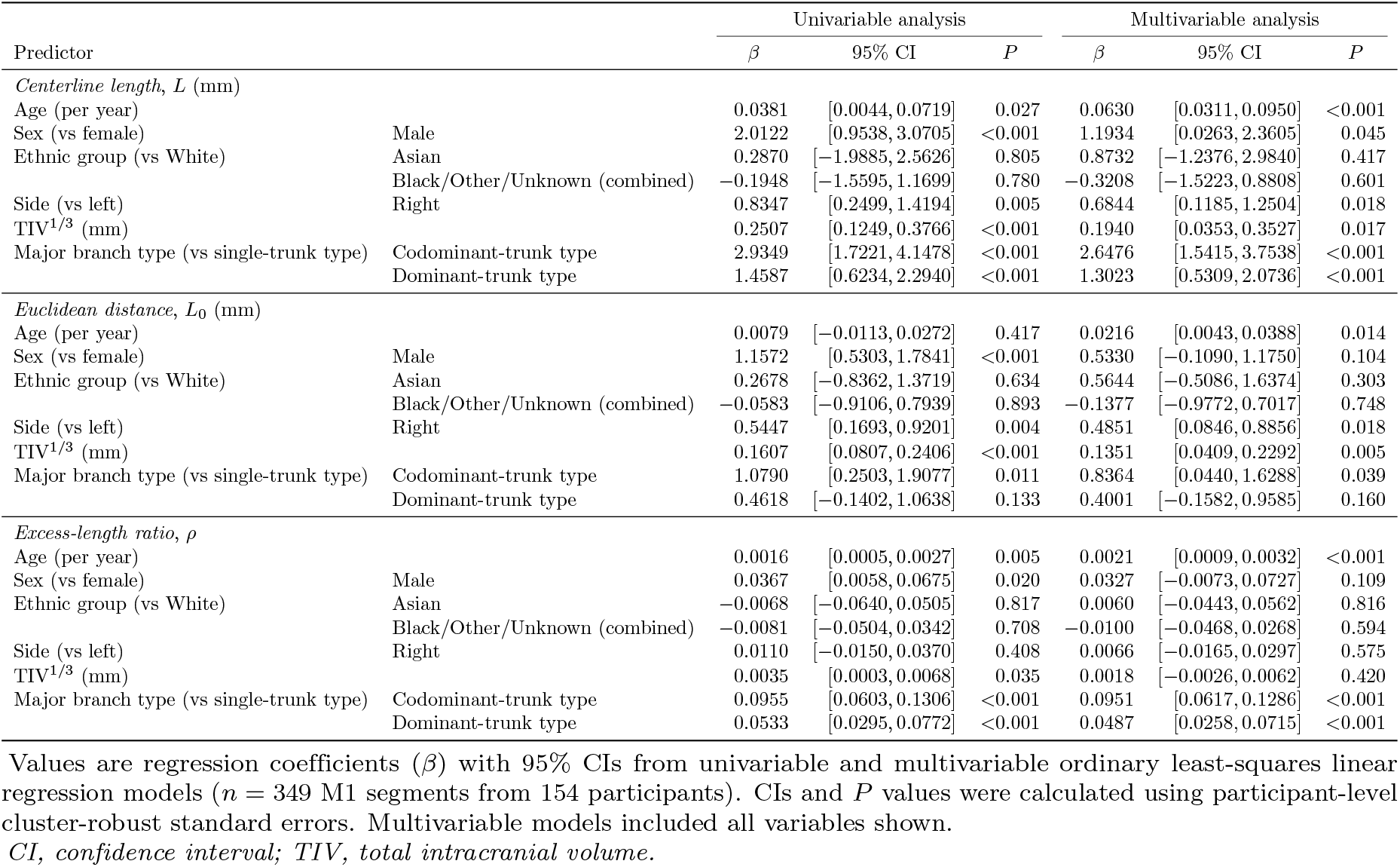
Univariable and multivariable linear regression models for M1 morphological parameters.

| Predictor |  | Univariable analysis |  |  | Multivariable analysis |  |  |
|---|---|---|---|---|---|---|---|
|  |  | *β* | 95% CI | *P* | *β* | 95% CI | *P* |
| *Centerline length*, *L* (mm) |  |  |  |  |  |  |  |
| Age (per year) |  | 0.0381 | [0.0044, 0.0719] | 0.027 | 0.0630 | [0.0311, 0.0950] | <0.001 |
| Sex (vs female) | Male | 2.0122 | [0.9538, 3.0705] | <0.001 | 1.1934 | [0.0263, 2.3605] | 0.045 |
| Ethnic group (vs White) | Asian | 0.2870 | [−1.9885, 2.5626] | 0.805 | 0.8732 | [−1.2376, 2.9840] | 0.417 |
|  | Black/Other/Unknown (combined) | −0.1948 | [−1.5595, 1.1699] | 0.780 | −0.3208 | [−1.5223, 0.8808] | 0.601 |
| Side (vs left) | Right | 0.8347 | [0.2499, 1.4194] | 0.005 | 0.6844 | [0.1185, 1.2504] | 0.018 |
| TIV<sup>1/3</sup> (mm) |  | 0.2507 | [0.1249, 0.3766] | <0.001 | 0.1940 | [0.0353, 0.3527] | 0.017 |
| Major branch type (vs single-trunk type) | Codominant-trunk type | 2.9349 | [1.7221, 4.1478] | <0.001 | 2.6476 | [1.5415, 3.7538] | <0.001 |
|  | Dominant-trunk type | 1.4587 | [0.6234, 2.2940] | <0.001 | 1.3023 | [0.5309, 2.0736] | <0.001 |
| *Euclidean distance*, *L*<sub>0</sub> (mm) |  |  |  |  |  |  |  |
| Age (per year) |  | 0.0079 | [−0.0113, 0.0272] | 0.417 | 0.0216 | [0.0043, 0.0388] | 0.014 |
| Sex (vs female) | Male | 1.1572 | [0.5303, 1.7841] | <0.001 | 0.5330 | [−0.1090, 1.1750] | 0.104 |
| Ethnic group (vs White) | Asian | 0.2678 | [−0.8362, 1.3719] | 0.634 | 0.5644 | [−0.5086, 1.6374] | 0.303 |
|  | Black/Other/Unknown (combined) | −0.0583 | [−0.9106, 0.7939] | 0.893 | −0.1377 | [−0.9772, 0.7017] | 0.748 |
| Side (vs left) | Right | 0.5447 | [0.1693, 0.9201] | 0.004 | 0.4851 | [0.0846, 0.8856] | 0.018 |
| TIV<sup>1/3</sup> (mm) |  | 0.1607 | [0.0807, 0.2406] | <0.001 | 0.1351 | [0.0409, 0.2292] | 0.005 |
| Major branch type (vs single-trunk type) | Codominant-trunk type | 1.0790 | [0.2503, 1.9077] | 0.011 | 0.8364 | [0.0440, 1.6288] | 0.039 |
|  | Dominant-trunk type | 0.4618 | [−0.1402, 1.0638] | 0.133 | 0.4001 | [−0.1582, 0.9585] | 0.160 |
| *Excess-length ratio*, *ρ* |  |  |  |  |  |  |  |
| Age (per year) |  | 0.0016 | [0.0005, 0.0027] | 0.005 | 0.0021 | [0.0009, 0.0032] | <0.001 |
| Sex (vs female) | Male | 0.0367 | [0.0058, 0.0675] | 0.020 | 0.0327 | [−0.0073, 0.0727] | 0.109 |
| Ethnic group (vs White) | Asian | −0.0068 | [−0.0640, 0.0505] | 0.817 | 0.0060 | [−0.0443, 0.0562] | 0.816 |
|  | Black/Other/Unknown (combined) | −0.0081 | [−0.0504, 0.0342] | 0.708 | −0.0100 | [−0.0468, 0.0268] | 0.594 |
| Side (vs left) | Right | 0.0110 | [−0.0150, 0.0370] | 0.408 | 0.0066 | [−0.0165, 0.0297] | 0.575 |
| TIV<sup>1/3</sup> (mm) |  | 0.0035 | [0.0003, 0.0068] | 0.035 | 0.0018 | [−0.0026, 0.0062] | 0.420 |
| Major branch type (vs single-trunk type) | Codominant-trunk type | 0.0955 | [0.0603, 0.1306] | <0.001 | 0.0951 | [0.0617, 0.1286] | <0.001 |
|  | Dominant-trunk type | 0.0533 | [0.0295, 0.0772] | <0.001 | 0.0487 | [0.0258, 0.0715] | <0.001 |

Values are regression coefficients (*β*) with 95% CIs from univariable and multivariable ordinary least-squares linear regression models (*n* = 349 M1 segments from 154 participants). CIs and *P* values were calculated using participant-level cluster-robust standard errors. Multivariable models included all variables shown.
*CI, confidence interval; TIV, total intracranial volume.*

**Table S4.**
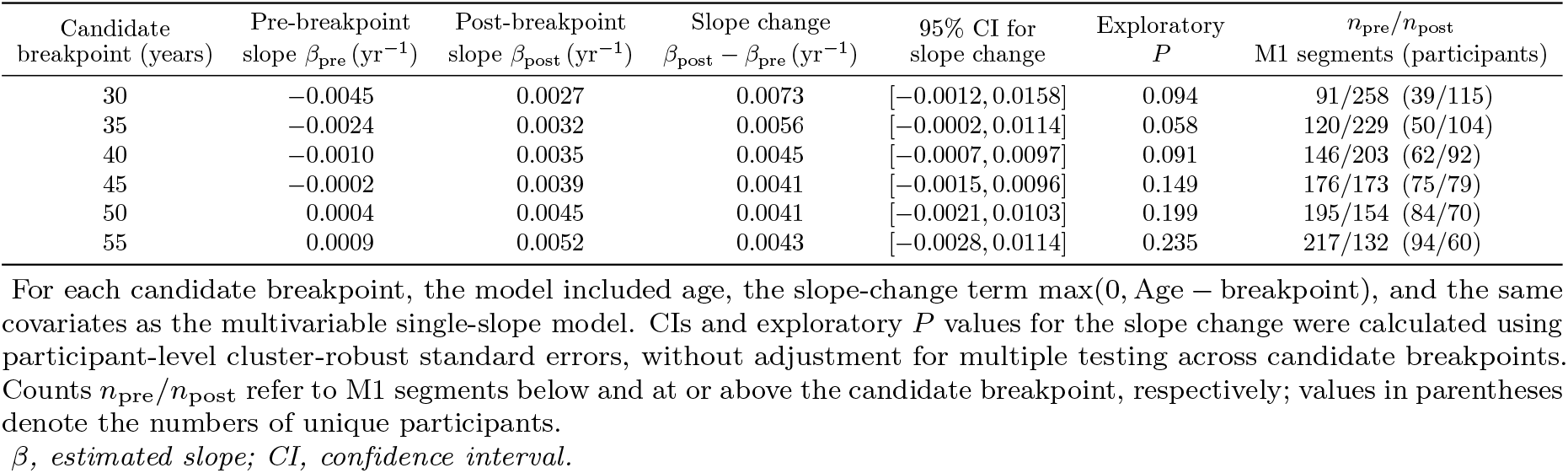
Exploratory piecewise models for excess-length ratio *ρ*.

| Candidate<br>breakpoint (years) | Pre-breakpoint<br>slope β<sub>pre</sub> (yr<sup>−1</sup>) | Post-breakpoint<br>slope β<sub>post</sub> (yr<sup>−1</sup>) | Slope change<br>β<sub>post</sub> − β<sub>pre</sub> (yr<sup>−1</sup>) | 95% CI for<br>slope change | Exploratory<br>*P* | *n*<sub>pre</sub>/*n*<sub>post</sub><br>M1 segments (participants) |
|---|---|---|---|---|---|---|
| 30 | −0.0045 | 0.0027 | 0.0073 | [−0.0012, 0.0158] | 0.094 | 91/258 (39/115) |
| 35 | −0.0024 | 0.0032 | 0.0056 | [−0.0002, 0.0114] | 0.058 | 120/229 (50/104) |
| 40 | −0.0010 | 0.0035 | 0.0045 | [−0.0007, 0.0097] | 0.091 | 146/203 (62/92) |
| 45 | −0.0002 | 0.0039 | 0.0041 | [−0.0015, 0.0096] | 0.149 | 176/173 (75/79) |
| 50 | 0.0004 | 0.0045 | 0.0041 | [−0.0021, 0.0103] | 0.199 | 195/154 (84/70) |
| 55 | 0.0009 | 0.0052 | 0.0043 | [−0.0028, 0.0114] | 0.235 | 217/132 (94/60) |

For each candidate breakpoint, the model included age, the slope-change term max(0, Age − breakpoint), and the same covariates as the multivariable single-slope model. CIs and exploratory *P* values for the slope change were calculated using participant-level cluster-robust standard errors, without adjustment for multiple testing across candidate breakpoints. Counts *n*<sub>pre</sub>/*n*<sub>post</sub> refer to M1 segments below and at or above the candidate breakpoint, respectively; values in parentheses denote the numbers of unique participants.
*β, estimated slope; CI, confidence interval.*

**Table S5.**
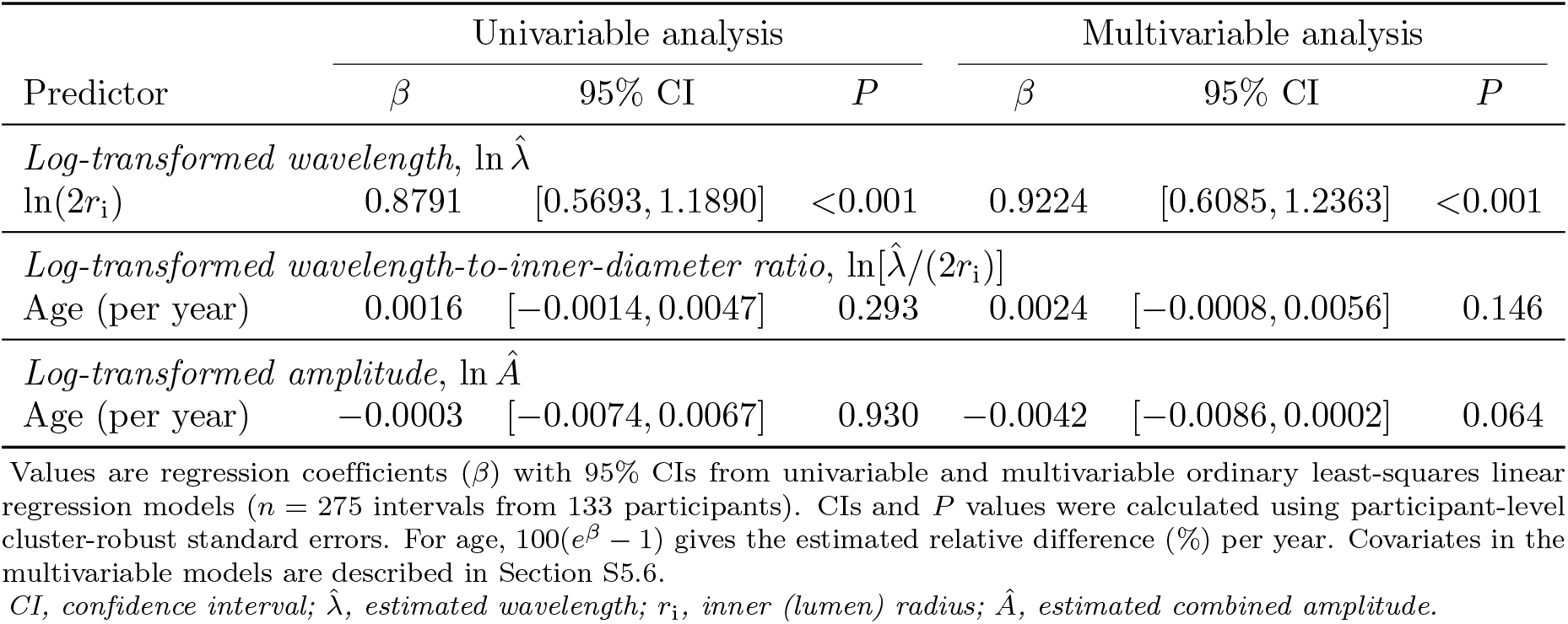
Univariable and multivariable linear regression models for periodic arterial wave parameters.

|  | Univariable analysis |  |  | Multivariable analysis |  |  |
|---|---|---|---|---|---|---|
| Predictor | *β* | 95% CI | *P* | *β* | 95% CI | *P* |
| *Log-transformed wavelength*, ln λ̂ |  |  |  |  |  |  |
| ln(2*r*<sub>i</sub>) | 0.8791 | [0.5693, 1.1890] | <0.001 | 0.9224 | [0.6085, 1.2363] | <0.001 |
| *Log-transformed wavelength-to-inner-diameter ratio*, ln[λ̂/(2*r*<sub>i</sub>)] |  |  |  |  |  |  |
| Age (per year) | 0.0016 | [−0.0014, 0.0047] | 0.293 | 0.0024 | [−0.0008, 0.0056] | 0.146 |
| *Log-transformed amplitude*, ln Â |  |  |  |  |  |  |
| Age (per year) | −0.0003 | [−0.0074, 0.0067] | 0.930 | −0.0042 | [−0.0086, 0.0002] | 0.064 |

Values are regression coefficients (*β*) with 95% CIs from univariable and multivariable ordinary least-squares linear regression models (*n* = 275 intervals from 133 participants). CIs and *P* values were calculated using participant-level cluster-robust standard errors. For age, 100(*e*<sup>*β*</sup> − 1) gives the estimated relative difference (%) per year. Covariates in the multivariable models are described in Section S5.6.
*CI, confidence interval; λ̂, estimated wavelength; r*<sub>i</sub>*, inner (lumen) radius; Â, estimated combined amplitude.*

**Table S6.**
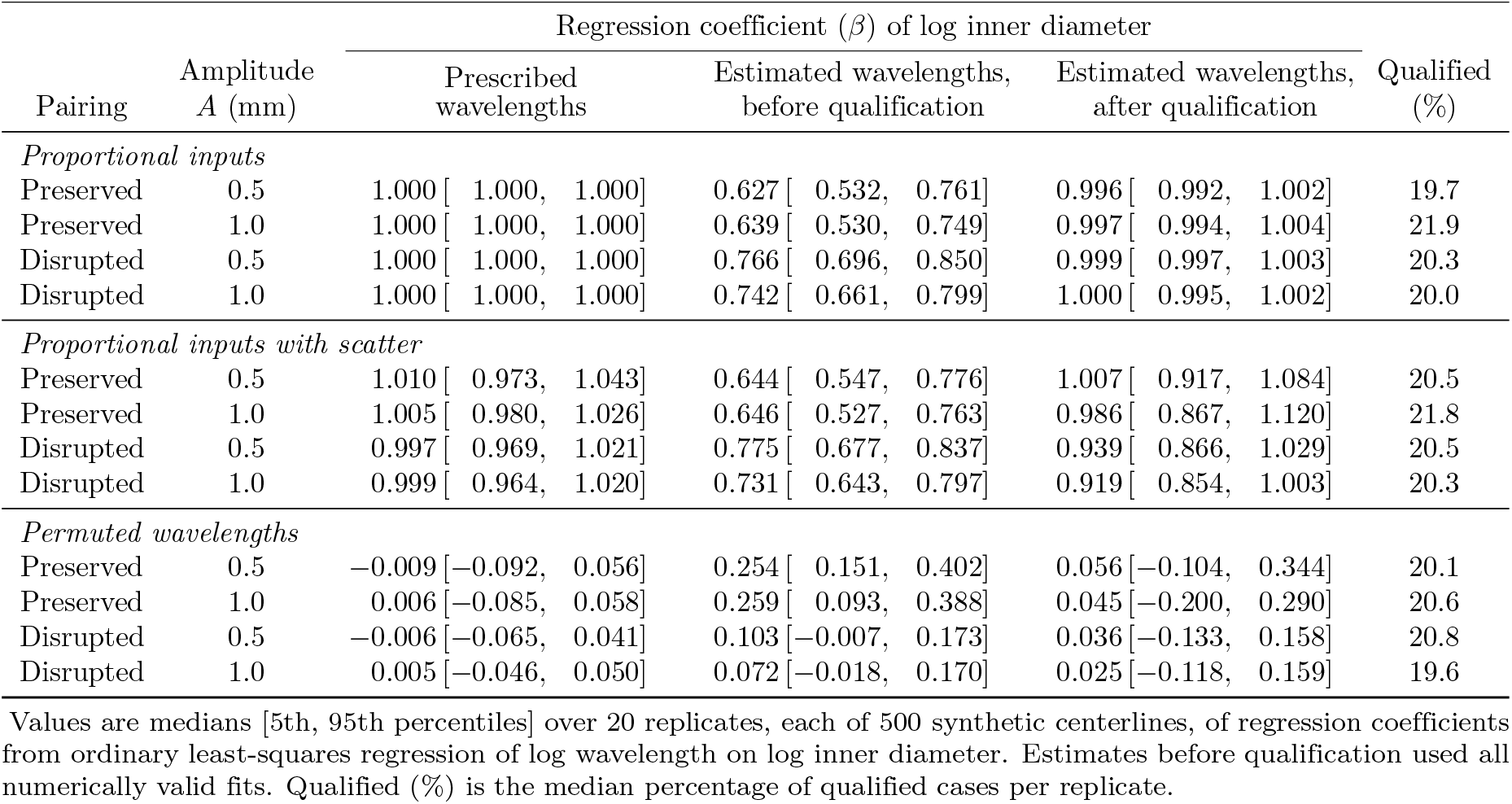
Coefficients of log inner diameter in the simulations of procedure-induced bias.

|  |  | Regression coefficient (*β*) of log inner diameter |  |  |  |
|---|---|---|---|---|---|
| Pairing | Amplitude *A* (mm) | Prescribed wavelengths | Estimated wavelengths, before qualification | Estimated wavelengths, after qualification | Qualified (%) |
| *Proportional inputs* |  |  |  |  |  |
| Preserved | 0.5 | 1.000 [ 1.000, 1.000] | 0.627 [ 0.532, 0.761] | 0.996 [ 0.992, 1.002] | 19.7 |
| Preserved | 1.0 | 1.000 [ 1.000, 1.000] | 0.639 [ 0.530, 0.749] | 0.997 [ 0.994, 1.004] | 21.9 |
| Disrupted | 0.5 | 1.000 [ 1.000, 1.000] | 0.766 [ 0.696, 0.850] | 0.999 [ 0.997, 1.003] | 20.3 |
| Disrupted | 1.0 | 1.000 [ 1.000, 1.000] | 0.742 [ 0.661, 0.799] | 1.000 [ 0.995, 1.002] | 20.0 |
| *Proportional inputs with scatter* |  |  |  |  |  |
| Preserved | 0.5 | 1.010 [ 0.973, 1.043] | 0.644 [ 0.547, 0.776] | 1.007 [ 0.917, 1.084] | 20.5 |
| Preserved | 1.0 | 1.005 [ 0.980, 1.026] | 0.646 [ 0.527, 0.763] | 0.986 [ 0.867, 1.120] | 21.8 |
| Disrupted | 0.5 | 0.997 [ 0.969, 1.021] | 0.775 [ 0.677, 0.837] | 0.939 [ 0.866, 1.029] | 20.5 |
| Disrupted | 1.0 | 0.999 [ 0.964, 1.020] | 0.731 [ 0.643, 0.797] | 0.919 [ 0.854, 1.003] | 20.3 |
| *Permuted wavelengths* |  |  |  |  |  |
| Preserved | 0.5 | −0.009 [−0.092, 0.056] | 0.254 [ 0.151, 0.402] | 0.056 [−0.104, 0.344] | 20.1 |
| Preserved | 1.0 | 0.006 [−0.085, 0.058] | 0.259 [ 0.093, 0.388] | 0.045 [−0.200, 0.290] | 20.6 |
| Disrupted | 0.5 | −0.006 [−0.065, 0.041] | 0.103 [−0.007, 0.173] | 0.036 [−0.133, 0.158] | 20.8 |
| Disrupted | 1.0 | 0.005 [−0.046, 0.050] | 0.072 [−0.018, 0.170] | 0.025 [−0.118, 0.159] | 19.6 |

Values are medians [5th, 95th percentiles] over 20 replicates, each of 500 synthetic centerlines, of regression coefficients from ordinary least-squares regression of log wavelength on log inner diameter. Estimates before qualification used all numerically valid fits. Qualified (%) is the median percentage of qualified cases per replicate.

**Table S7.** Baseline characteristics of the study population.

| Characteristic (*n* = 154) |  | Value |
|---|---|---|
| Age (years) |  | 45.5 (29.9–61.7) |
| Sex | Male | 72 (46.8) |
|  | Female | 82 (53.2) |
| Ethnic group | White | 117 (76.0) |
|  | Asian | 20 (13.0) |
|  | Black | 6 (3.9) |
|  | Other | 4 (2.6) |
|  | Unknown | 7 (4.5) |
| Body mass index (kg/m<sup>2</sup>) |  | 24.3 (22.2–26.9) |
| Total intracranial volume (mL) |  | 1452 (1323–1562) |

Data are presented as number (%) or median (interquartile range). Total intracranial volume was estimated using the Computational Anatomy Toolbox 12.

**Table S8.**
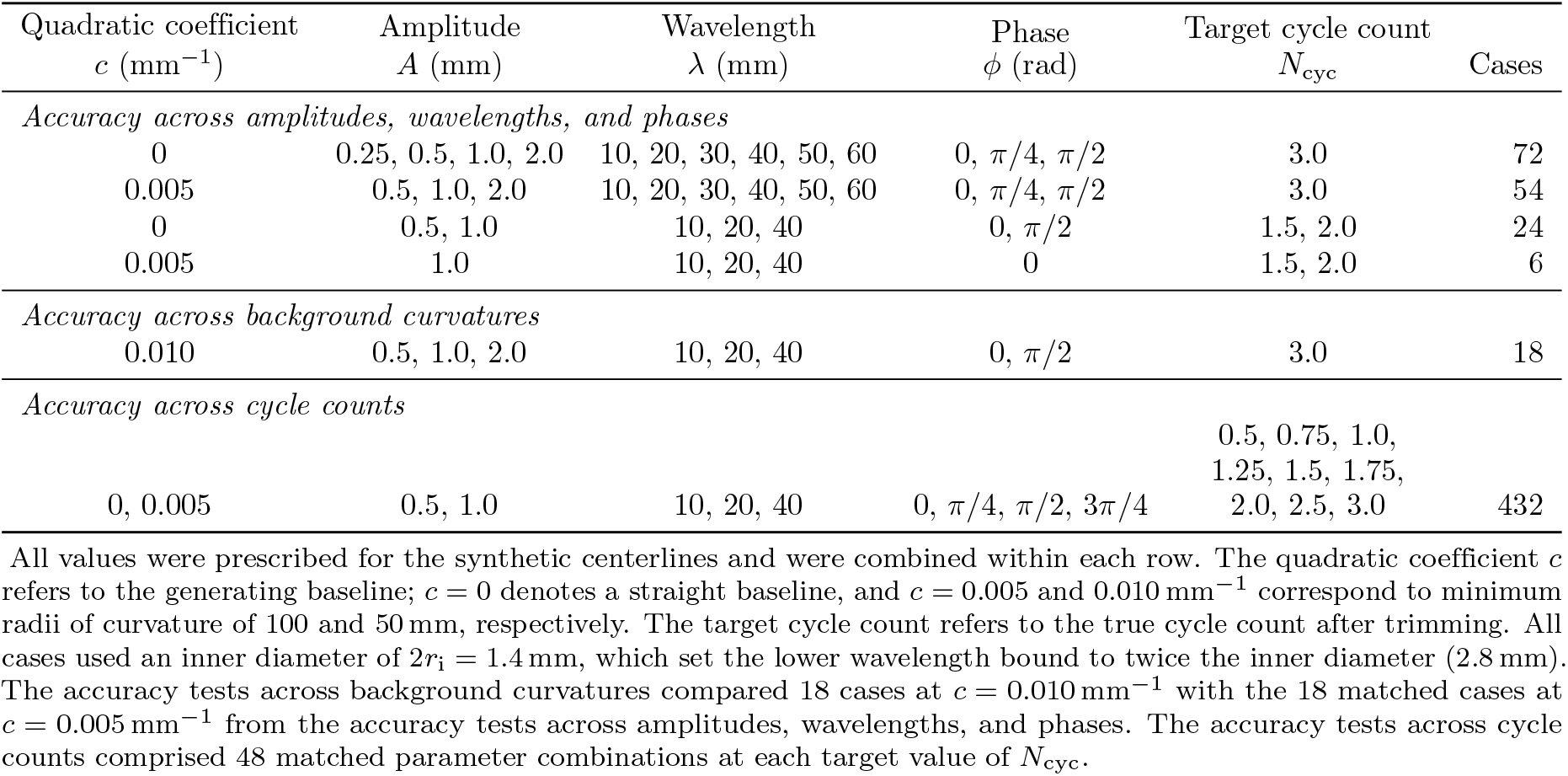
Conditions used in the accuracy tests with synthetic centerlines.

| Quadratic coefficient<br>*c* (mm<sup>−1</sup>) | Amplitude<br>*A* (mm) | Wavelength<br>*λ* (mm) | Phase<br>*ϕ* (rad) | Target cycle count<br>*N*<sub>cyc</sub> | Cases |
|---|---|---|---|---|---|
| *Accuracy across amplitudes, wavelengths, and phases* |  |  |  |  |  |
| 0 | 0.25, 0.5, 1.0, 2.0 | 10, 20, 30, 40, 50, 60 | 0, π/4, π/2 | 3.0 | 72 |
| 0.005 | 0.5, 1.0, 2.0 | 10, 20, 30, 40, 50, 60 | 0, π/4, π/2 | 3.0 | 54 |
| 0 | 0.5, 1.0 | 10, 20, 40 | 0, π/2 | 1.5, 2.0 | 24 |
| 0.005 | 1.0 | 10, 20, 40 | 0 | 1.5, 2.0 | 6 |
| *Accuracy across background curvatures* |  |  |  |  |  |
| 0.010 | 0.5, 1.0, 2.0 | 10, 20, 40 | 0, π/2 | 3.0 | 18 |
| *Accuracy across cycle counts* |  |  |  |  |  |
| 0, 0.005 | 0.5, 1.0 | 10, 20, 40 | 0, π/4, π/2, 3π/4 | 0.5, 0.75, 1.0, 1.25, 1.5, 1.75, 2.0, 2.5, 3.0 | 432 |

All values were prescribed for the synthetic centerlines and were combined within each row. The quadratic coefficient *c* refers to the generating baseline; *c* = 0 denotes a straight baseline, and *c* = 0.005 and 0.010 mm<sup>−1</sup> correspond to minimum radii of curvature of 100 and 50 mm, respectively. The target cycle count refers to the true cycle count after trimming. All cases used an inner diameter of 2*r*<sub>i</sub> = 1.4 mm, which set the lower wavelength bound to twice the inner diameter (2.8 mm). The accuracy tests across background curvatures compared 18 cases at *c* = 0.010 mm<sup>−1</sup> with the 18 matched cases at *c* = 0.005 mm<sup>−1</sup> from the accuracy tests across amplitudes, wavelengths, and phases. The accuracy tests across cycle counts comprised 48 matched parameter combinations at each target value of *N*<sub>cyc</sub>.

